# Design, synthesis and pharmacological characterization of the first photoswitchable small-molecule agonist for the Atypical Chemokine Receptor 3

**DOI:** 10.1101/2024.03.20.585914

**Authors:** Sophie Bérenger, Justyna M. Adamska, Francesca Deflorian, Chris de Graaf, Christie B. Palmer, Martyna Szpakowska, Andy Chevigné, Iwan J. P. de Esch, Barbara Zarzycka, Henry F. Vischer, Maikel Wijtmans, Rob Leurs

## Abstract

Photopharmacology offers the promise of optical modulation of cellular signaling in a spatially and temporally controlled fashion with light-sensitive molecules. This study presents the first small-molecule photoswitchable agonist for an atypical G protein-coupled receptor (GPCR), the atypical chemokine receptor 3 (ACKR3). Inspired by a known benzylpiperidine-based ACKR3 agonist scaffold, 12 photoswitchable azobenzene-containing analogs were synthesized and characterized for their interaction with ACKR3. After analysis of Structure-Photochemistry and Structure-Affinity Relationships (SAR), compound **3e** was selected as the best photoswitchable ACKR3 agonist in the series. Compound **3e** can be effectively switched from its thermodynamically stable *trans* state to the less active *cis*-isomer with a PhotoStationary State of 96 %. The thermodynamically less stable *cis-***3e** only slowly switches back to the *trans* state (t_1/2,37 °C_ = 15 days), and *trans-***3e** binds and activates ACKR3 at 10-fold lower concentrations compared to its *cis*-isomer. Compound **3e** demonstrates selectivity for ACKR3 within in a panel of chemokine receptors. Using the recently published ACKR3 cryo-EM structures in computational studies, a binding mode for *trans-***3e** is proposed and is perfectly in line with the observed SAR and the loss of interaction with ACKR3 upon photoswitching. ACKR3 agonist **3e** (VUF25471) is the first photoswitchable ligand for an atypical GPCR and will be a useful tool to investigate the role of ACKR3 in biological settings.

## 2. Introduction

Photopharmacology uses photo-responsive ligands to obtain spatiotemporal control of protein function via optical modulation^1–3^. Indeed, light is, in principle, ideal for spatiotemporal modulation owing to its ease of dosing and non-invasive nature^4^, and in the last decade, this feature has been successfully applied in the field of optogenetics^5,6^. The field of photopharmacology is complementary to optogenetics and offers optical modulation of native protein function via the use of photocaged or photoswitchable molecules^1–3^. Molecules containing a photoswitchable moiety can reversibly isomerize upon illumination at specific wavelengths resulting in a reversible configurational change. Ideally, the isomerization of a photoswitch is accompanied by a distinct change in the interaction of the molecule with the targeted protein. Photoswitchable ligands have been developed for a number of biologically relevant protein classes, including photoswitches acting on enzymes, ion channels and G protein-coupled receptors (GPCR)^1–3^. To date, various types of photoswitchable ligands for Class A GPCRs have been described^1,7–9^. These studies underscore how a balance needs to be found between the availability of a suitable template ligand, the synthetic feasibility and photochemical properties of the photoswitchable ligand, and the differential recognition of the isomers by the target GPCR^1^.

The first and so far only photoswitchable ligand for a GPCR was described in 2018 and involved a small-molecule antagonist-to-agonist efficacy switch for the chemokine receptor CXCR3^10,11^. Chemokine receptors form a large subfamily of at least 23 class A GPCRs that orchestrate immune cell trafficking and have been identified as important targets for several disorders^12–14^. Lately, a subfamily of at least five *atypical* chemokine receptors (ACKRs) has been suggested as key regulators of chemokine function^14–16^. These atypical GPCRs all bind a variety of chemokines, but do not signal following the canonical G-protein pathways. In contrast, the ACKRs act as β-arrestin-biased GPCRs and are involved in the regulation of extracellular chemokine levels mostly via chemokine internalization and breakdown^14–16^.

In the current study, we report on the first photoswitchable ligand for such an intrinsically β-arrestin-biased ACKR, the atypical chemokine receptor 3 (ACKR3). Initially discovered as RDC-1^17^ and renamed first CXCR7 and thereafter ACKR3, this atypical GPCR binds and scavenges the chemokines CXCL11 and CXCL12, thereby controlling their availability for their classical receptors, CXCR3 and CXCR4, respectively^17–20^. Moreover, overexpression of ACKR3 has been observed in immune, cardiovascular, neurodegenerative, and oncological disorders^21,22^. Consequently, small-molecule ACKR3 ligands have been developed to explore the biology of this atypical GPCR and provide starting points for novel therapeutics^23,24,33–40,25–32^.

To provide complementary chemical biology tools from a photopharmacology approach, this study reports on a series of photoswitchable ACKR3 ligands (**3a-l**). These ligands are based on a scaffold patented by Actelion Pharmaceuticals^41^, and have been obtained following an azologization strategy, *i.e.* incorporating the light-sensitive azobenzene moiety in the main scaffold. The azobenzene moiety is a popular photoswitchable moiety^1–3^ and is known for its small size, synthetic accessibility, robust photoswitching character and the possibility to modify its thermal relaxation rate and absorption characteristics by substitution^2,42–44^. The introduction of the azobenzene unit resulted in the successful generation of a number of photoswitchable molecules from which **3e** (VUF25471) emerges as a robust photoswitchable small-molecule agonist to modulate β-arrestin2 recruitment to ACKR3 with light.

## 3. Results and Discussion

### 3.1. Design

In the search for a suitable template for a photoswitchable ACKR3 ligand, we evaluated a 2013 patent from Actelion Pharmaceuticals^41^ where 428 substituted benzylpiperidines (see Markush structure **1**) are described as ACKR3 ligands (Fig. 1A). These compounds are reported to display nanomolar to micromolar agonist potencies, as determined in a PathHunter® β-arrestin2 recruitment assay. *Meta*-benzamide derivatives were amongst the most potent derivatives explored. Seeking for a scaffold with good synthetic feasibility to develop photoswitchable ACKR3 ligands, a *N*-(*tert*-butyl)carboxamide moiety in conjunction with a *meta-*phenyl-benzamide, attached to the piperidine via a one-carbon linker, was selected. This scaffold is exemplified by the *para* F-substituted analogue **2** (Fig. 1A), which was reported in the patent to recruit β-arrestin2 with sub-nanomolar potency (pEC50 = 9.3)^41^.

**Figure 1.**
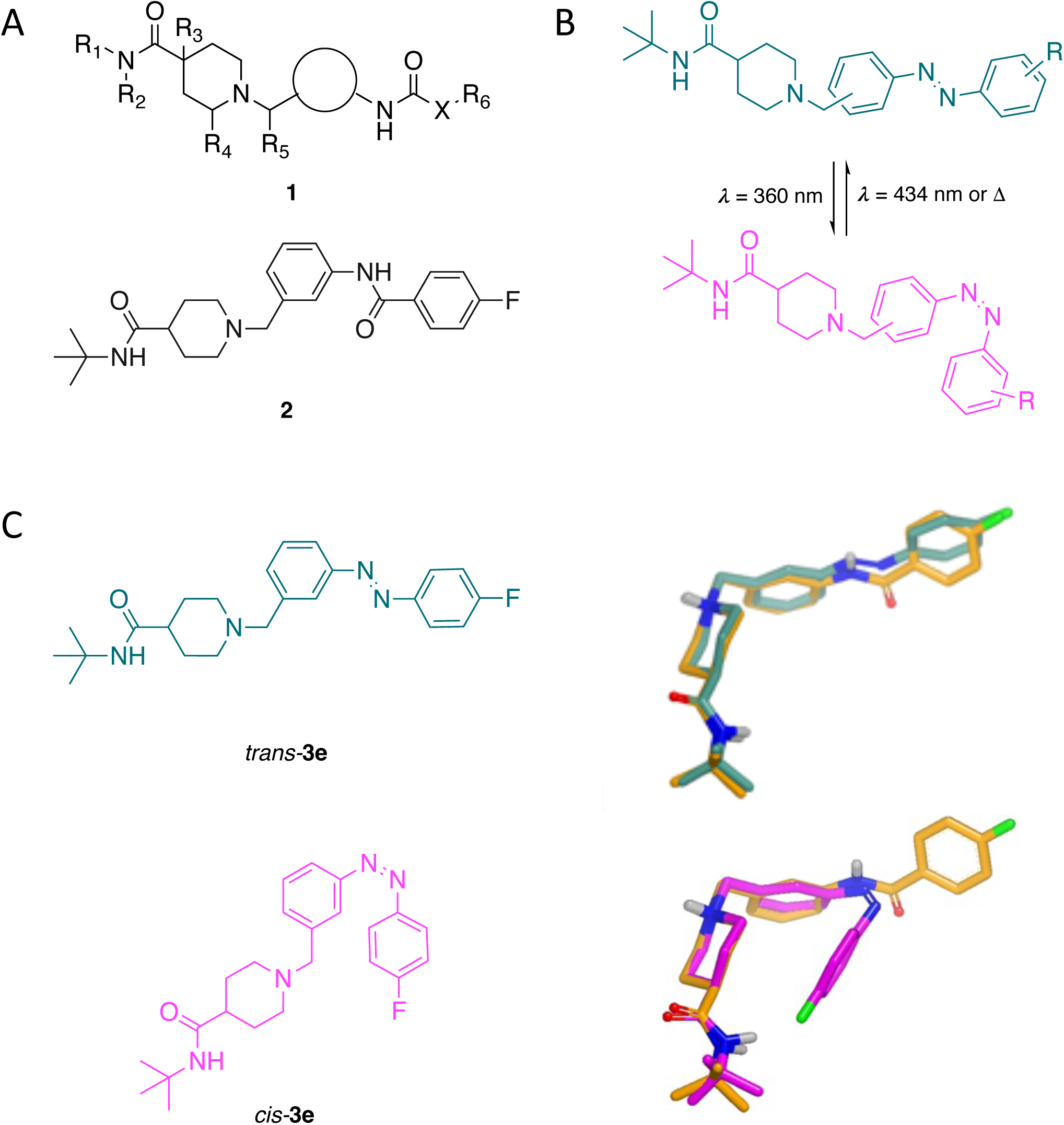
General design strategy. (A) Markush structure from patent US9428456^41^ (1) and structure of a representative member (**2**) of the series. (B) Azologization strategy based on **2**: isomerization of the *trans* analogs leads to the *cis* isomers after illumination at 360 nm. The switch is reversible by illumination at 434 nm or by thermal relaxation. (C) Structures of *trans*- and *cis*-**3e** (green and magenta, respectively), superposition of the minimum energy conformation of **2** (orange carbon atoms) and that of *trans*-**3e** (green carbon atoms, top right) and superposition of the minimum energy conformation of **2** (orange carbon atoms) and that of *cis*-**3e** (magenta carbon atoms, bottom right).

The azosteric amide moiety between the two aryl rings of **2** appears to be a suitable candidate for an azologization strategy^2,3^. It was reasoned that the substitution of the bisarylamide in **2** by an azobenzene and a subsequent concise Structure-Activity Relationship (SAR) exploration could provide a suitable photoswitchable (tool) compound (Fig. 1B). To support this assumption, a pharmacophore alignment of energy-minimized structures was performed on template ligand **2**, and the azologization-derived analogue **3e** (Fig. 1C). Superposition of low-energy conformations of **2** and **3e** highlights a 3D conformational similarity between **2** and *trans*-**3e**, with the azobenzene moiety of *trans*-**3e** overlapping with the benzamide moiety of **2**, suggesting that both molecules can adopt similar binding modes in the ACKR3 binding site. In contrast, the folded azobenzene conformation adopted by *cis*-**3e** in its minimum energy conformation shows no overlap of the terminal phenyl moiety with that of **2**. Indeed, it is well established that the *cis*- azobenzene moiety is not planar but folded with the distance between the two *ortho*-carbons being around 3.3 Å^45^. Based on this superposition, the *trans* isomer of the photoswitchable *meta*-scaffold was expected to interact better with ACKR3 than its corresponding *cis* isomer. A focused SAR study was foreseen to explore this hypothesis. Inserting the azobenzene moiety at the *ortho*, *meta* or *para* position with respect to the methylene substituent induces significant shape modifications of the molecules. To modulate the electron density of the azobenzene moiety, electron-donating and -withdrawing groups were introduced at several positions of the azobenzene core. Analysis of the patent on the template scaffold^41^ suggests that the substitution of the internal aryl ring does not bode well for significant improvements in affinity.

Therefore, modification of this ring was not pursued. Instead, according to the patent data, substitution at the *ortho*, *meta* and *para* position of the terminal aryl ring is tolerated, with *meta* and *para* substitution preferred. Analogues exploring this side of the scaffold were thus included in the present SAR study.

### 3.2. Synthesis

Two routes were followed to synthesize the photoswitchable ligands (Scheme 1). The first one allowed the introduction of substituents at the terminal phenyl moiety in the final step. Acid **4** was transformed to (its) acid chloride before undergoing a substitution by *tert*-butylamine to afford **5**. The piperidine nitrogen atom was deprotected and the resulting amine **6** was reacted in a reductive amination with commercially available **7a-c**. The resulting *ortho*, *meta*, and *para*-anilines **8a-c** were reacted with nitroso compounds **10a- e**, made via the commercially available corresponding anilines **9a-e** under acidic Mills conditions. Ten photoswitchable ligands **3a-f,h-j,l** were obtained through this strategy. Template **2** was obtained by acylation of aniline **8b** with 4-fluorobenzoyl chloride. The second route was designed to provide compounds **3g** and **3k**, as the Mills reactions in the first strategy gave low conversion and difficult work-up for these two substitution patterns. Aniline **11** was oxidized with Oxone^TM^ to give nitroso compound **12**, which was directly used in a Mills reaction with substituted anilines to afford azobenzenes **13a-b.** These were reduced to alcohols **14a-b** followed by an oxidation, either using Dess-Martin reagent (**15a**) or PCC (**15b**). Aldehydes **15a-b** were reacted with **6** in a reductive amination to afford **3g** and **3k**.

**Scheme 1:**
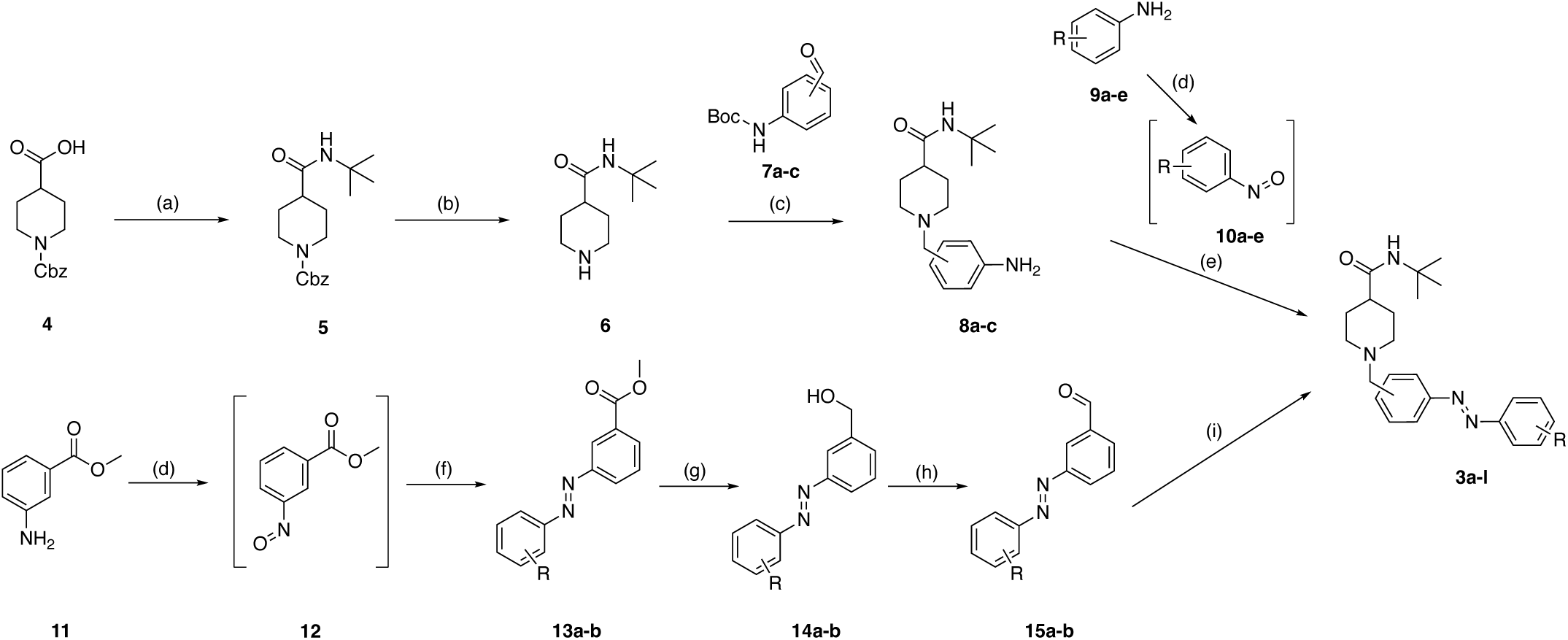
*Preparation of compounds.* Reagents and conditions: (a) (i) (COCl)2, DMF, DCM, rt, 4 h, (ii) *t*BuNH2, TEA, DCM, rt, on, 87%. (b) H2, Pd/C, MeOH, rt, 4 h, 93%. (c) (i) NaBH(OAc)3, DCM, rt, on. (ii) TFA, DCM, rt, on, 42-64%. (d) Oxone^TM^, DCM/H2O, 1-18 h, rt, used as such. (e) AcOH, DCM, rt - 95 °C, 16-72 h, 4-89%. (f) **9g/f**, AcOH, DCM, rt, 16h, 68-73%. (g) DIBAL-H, *c*Hex, 0 °C - rt, 16 h, 52-71%. (h) **15a**: Dess-Martin reagent, DCM, rt, 2 h, 80%; **15b**: PCC, DCM, rt, 2h, 96 %. (i) **6**, NaBH(OAc)3, DCM, rt, 16 h, 9-18%. For the structures to which the letters in **3a-l** correspond, see Table 1. For the structures to which the letters in intermediates correspond and for detailed experimental procedures, see Supporting Information.

### 3.3. Structure–Photochemistry and Structure–Activity Relationships

Photochemical and pharmacological characterisation of ligands **3a-l** were performed in parallel. This allowed for the simultaneous exploration of structure***–***photochemistry and structure***–***activity relationships (SAR), which are both crucial to select a key ligand. The photochemical parameters (Table 1, Fig. S1) were measured in 100% DMSO. These include the absorption maxima (λ_max_) of the *trans*- and *cis*-isomers to determine the wavelengths of illumination as well as the PhotoStationary State (PSS_*cis*_) values to estimate the efficiency of the *trans*-to-*cis* conversion. As expected, virtually all compounds show the classic absorption pattern of azobenzenes with a large band around 320 nm in the dark (π–π* transition of the *trans* isomer) and a weaker band around 420 nm after illumination (n–π* transition of the *cis* isomer). This absorption pattern can be explained by the terminal phenyl ring of the azobenzene having substituents that lack pronounced *π*-donating properties, the only exception being **3g** in which the MeO-group shifts the λ_max_,*trans* value to 356 nm. Interestingly, **3a** and **3d**, carrying an azobenzene *ortho* to the piperidine, show photodecomposition after a few minutes of illumination at 360 nm. The decomposition products could not be identified by LCMS or NMR. PSS_*cis*_ values were generally high (>90 %) except for CF3-analogue **3j** (84 %). The thermal half-lives were measured qualitatively for all compounds by LC-MS analysis. All half-lives were within the range of days at 25 °C, meaning that pharmacological studies could reliably be undertaken with preilluminated samples.

**Table 1.**
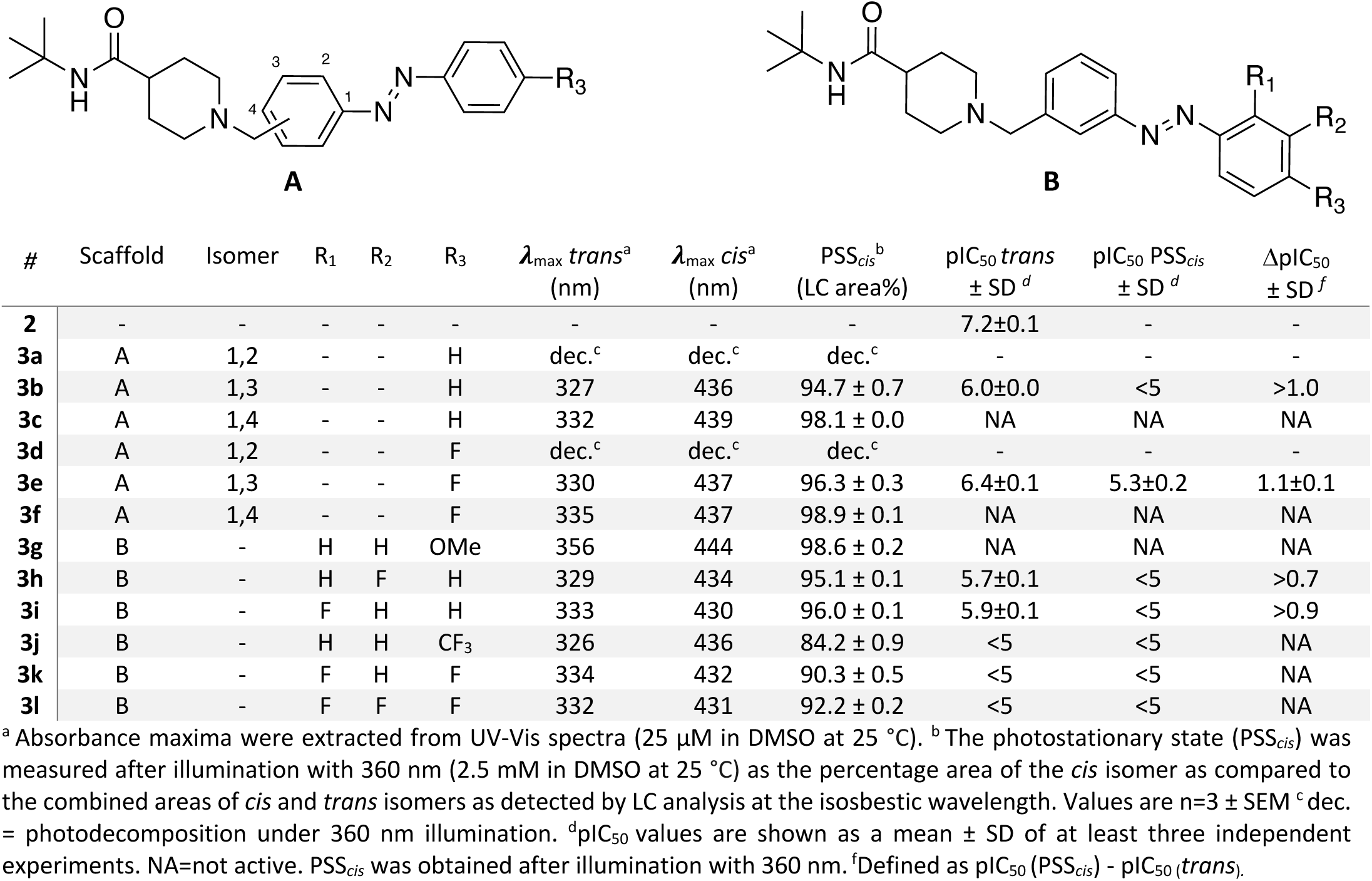
Structure−Photochemistry Relationship and Structure–Activity Relationship of photoswitchable ACKR3 ligands for displacing fluorescently labeled CXCL12 (A647®) to NLuc-ACKR3.

| # | Scaffold | Isomer | R<sub>1</sub> | R<sub>2</sub> | R<sub>3</sub> | $\lambda_{\max}^{trans^a}$<br>(nm) | $\lambda_{\max}^{cis^a}$<br>(nm) | PSS<sub>cis</sub><sup>b</sup><br>(LC area%) | pIC<sub>50</sub> *trans*<br>± SD<sup>d</sup> | pIC<sub>50</sub> PSS<sub>cis</sub><br>± SD<sup>d</sup> | ΔpIC<sub>50</sub><br>± SD<sup>f</sup> |
| --- | --- | --- | --- | --- | --- | --- | --- | --- | --- | --- | --- |
| **2** | - | - | - | - | - | - | - | - | 7.2±0.1 | - | - |
| **3a** | A | 1,2 | - | - | H | dec.<sup>c</sup> | dec.<sup>c</sup> | dec.<sup>c</sup> | - | - | - |
| **3b** | A | 1,3 | - | - | H | 327 | 436 | 94.7 ± 0.7 | 6.0±0.0 | <5 | >1.0 |
| **3c** | A | 1,4 | - | - | H | 332 | 439 | 98.1 ± 0.0 | NA | NA | NA |
| **3d** | A | 1,2 | - | - | F | dec.<sup>c</sup> | dec.<sup>c</sup> | dec.<sup>c</sup> | - | - | - |
| **3e** | A | 1,3 | - | - | F | 330 | 437 | 96.3 ± 0.3 | 6.4±0.1 | 5.3±0.2 | 1.1±0.1 |
| **3f** | A | 1,4 | - | - | F | 335 | 437 | 98.9 ± 0.1 | NA | NA | NA |
| **3g** | B | - | H | H | OMe | 356 | 444 | 98.6 ± 0.2 | NA | NA | NA |
| **3h** | B | - | H | F | H | 329 | 434 | 95.1 ± 0.1 | 5.7±0.1 | <5 | >0.7 |
| **3i** | B | - | F | H | H | 333 | 430 | 96.0 ± 0.1 | 5.9±0.1 | <5 | >0.9 |
| **3j** | B | - | H | H | CF<sub>3</sub> | 326 | 436 | 84.2 ± 0.9 | <5 | <5 | NA |
| **3k** | B | - | F | H | F | 334 | 432 | 90.3 ± 0.5 | <5 | <5 | NA |
| **3l** | B | - | F | F | F | 332 | 431 | 92.2 ± 0.2 | <5 | <5 | NA |

<sup>a</sup> Absorbance maxima were extracted from UV-Vis spectra (25 μM in DMSO at 25 °C). <sup>b</sup> The photostationary state (PSS<sub>cis</sub>) was measured after illumination with 360 nm (2.5 mM in DMSO at 25 °C) as the percentage area of the *cis* isomer as compared to the combined areas of *cis* and *trans* isomers as detected by LC analysis at the isosbestic wavelength. Values are n=3 ± SEM <sup>c</sup> dec. = photodecomposition under 360 nm illumination. <sup>d</sup> pIC<sub>50</sub> values are shown as a mean ± SD of at least three independent experiments. NA=not active. PSS<sub>cis</sub> was obtained after illumination with 360 nm. <sup>f</sup> Defined as pIC<sub>50</sub> (PSS<sub>cis</sub>) - pIC<sub>50</sub> (*trans*).

Initially, all *trans-* and PSS_*cis*_ states were tested for their binding to human ACKR3, using a NanoBRET binding assay employing fluorescently labelled CXCL12 (A647®) and membranes from HEK293T cells, transiently expressing human ACKR3 fused at its N-terminus to NanoLuciferase (NLuc-ACKR3).^46^ In this assay, CXCL12 (A647®) binds NLuc-ACKR3 with high affinity (0.5 ± 0.0 nM, n =3). Template compound **2** inhibited binding of CXCL12 (A647®) to NLuc-ACKR3 with a pIC50 value of 7.2 ± 0.1 (Table 1, Fig. 3A), which is 100-fold lower than its potency to recruit β-arrestin2 as reported in the patent describing **2**^41^.

In the SAR study (Table 1), compounds **3a-f** serve to explore the optimal position of the azo moiety with R3 being either H (**3a-c**) or F (**3d-f**) to mimic template **2** more closely. Only compounds **3b** and **3e**, with the azo moiety at the *meta* position, inhibited the binding of CXCL12 (A647®) to NLuc-ACKR3. A comparison between **3b** and **3e** highlights the benefit of the EWG fluorine atom at the terminal phenyl ring, with increased pIC50 values for both isomers of compound **3e**. In contrast, an EDG substituent at the *para* position (MeO-compound **3g**) abolished the ability to displace CXCL12 (A647®) from NLuc-ACKR3.

The next step aimed to determine the optimal position of the fluorine atom at the terminal phenyl ring. *Meta* and *ortho* fluorine-substituted azobenzenes **3h** and **3i** were examined, but neither of the two analogs bound to NLuc-ACKR3 as well as **3e**, revealing the *para*-position as the preferred anchor. However, introducing a *p*-CF3 group (**3j**) was detrimental for the interaction with ACKR3. Finally, while keeping an F atom at the *para*-position (**3k-l**), additional fluorine atoms were introduced on the terminal aryl ring, but none of the derivatives was as active as **3e**.

### 3.4. In-depth photochemistry of key compound 3e

Following the photochemical and pharmacological profiling, **3e** emerged as having the best profile in terms of PSS_*cis*_ value (96.3 %), absolute binding affinity for the *trans* isomer (6.4) and βpIC50 (1.1). An in-depth photochemical characterisation was performed (Fig. 2). In HBSS buffer with 1 % DMSO, *trans*-**3e** (Fig. 2A) shows a major band with a maximum absorption λ_max_ of 326 nm corresponding to the π–π* transition (Fig. 2B). Upon illumination at 360 nm, the compound switches to the PSS_*cis*_ state which consists mainly of the *cis*-**3e** isomer as characterized by an emerging small absorption band with a maximum λ_max_ of 424 nm, corresponding to the n–π* transition of *cis*-**3e** (Fig. 2B). The switch can be reversed using a lower-energy wavelength of illumination. Indeed, by applying 434 nm light to the PSS_*cis*_ state of **3e**, the PSS*trans* state can be obtained (Fig. 2B). The *trans*-to-*cis* switching was measured at different time points upon illumination at 360 nm (Fig. 2C). A gradual decrease of the π–π* band of *trans*-**3e** with a concomitant increase of the n– π* band of *cis*-**3e** indicated the establishment of the PSS_*cis*_ state within 32 s. The reversible switch was quantified by ^1^H NMR spectroscopy and LCMS analysis in the dark, after 360 and 434 nm illumination, respectively. By observing the ppm shift of the two protons at the *ortho* position of the terminal aryl group of the azobenzene moiety (Fig. 2D, Fig. S2), the PSS_*cis*_ and PSS*trans* values were quantified to be 87 and 78 mole %, respectively. The PSS states could also be observed by LC-MS analysis (Fig. 2E) and approximated as a percentage area at the isosbestic point (381 nm). The PSS_*cis*_ value after 360 nm illumination is 96.3 ± 0.3 %, while the PSS*trans* value after 434 nm illumination amounts to 77.3 ± 0.1 %. The lower PSS*trans* value compared to the PSS_*cis*_ value is expected for azobenzenes as the n–π* absorption bands of the *trans*- and *cis*-isomer significantly overlap^1–3^. Detection at 254 nm was also used (Fig. S3) to ensure that no unwanted photodecomposition occurs during the switching process. The thermal relaxation of the PSS_*cis*_ state of **3e** in HBSS buffer was assessed at 60 (Fig. 2F), 70 and 80 °C with detection at 320 nm, because the π–π* transition bands of both isomers absorb significantly differently at that wavelength. An Arrhenius plot was used to extrapolate the thermodynamic half-life to biologically relevant temperatures (Fig. S4)^47^. This afforded an estimated half-life of 216 and 15 days at 25 and 37 °C, respectively. Dynamic and repeated isomerization in buffer was studied under alternating illumination, using 320 nm detection. This confirmed robust and reversible switching over 10 cycles (Fig. 2G).

**Figure 2.**
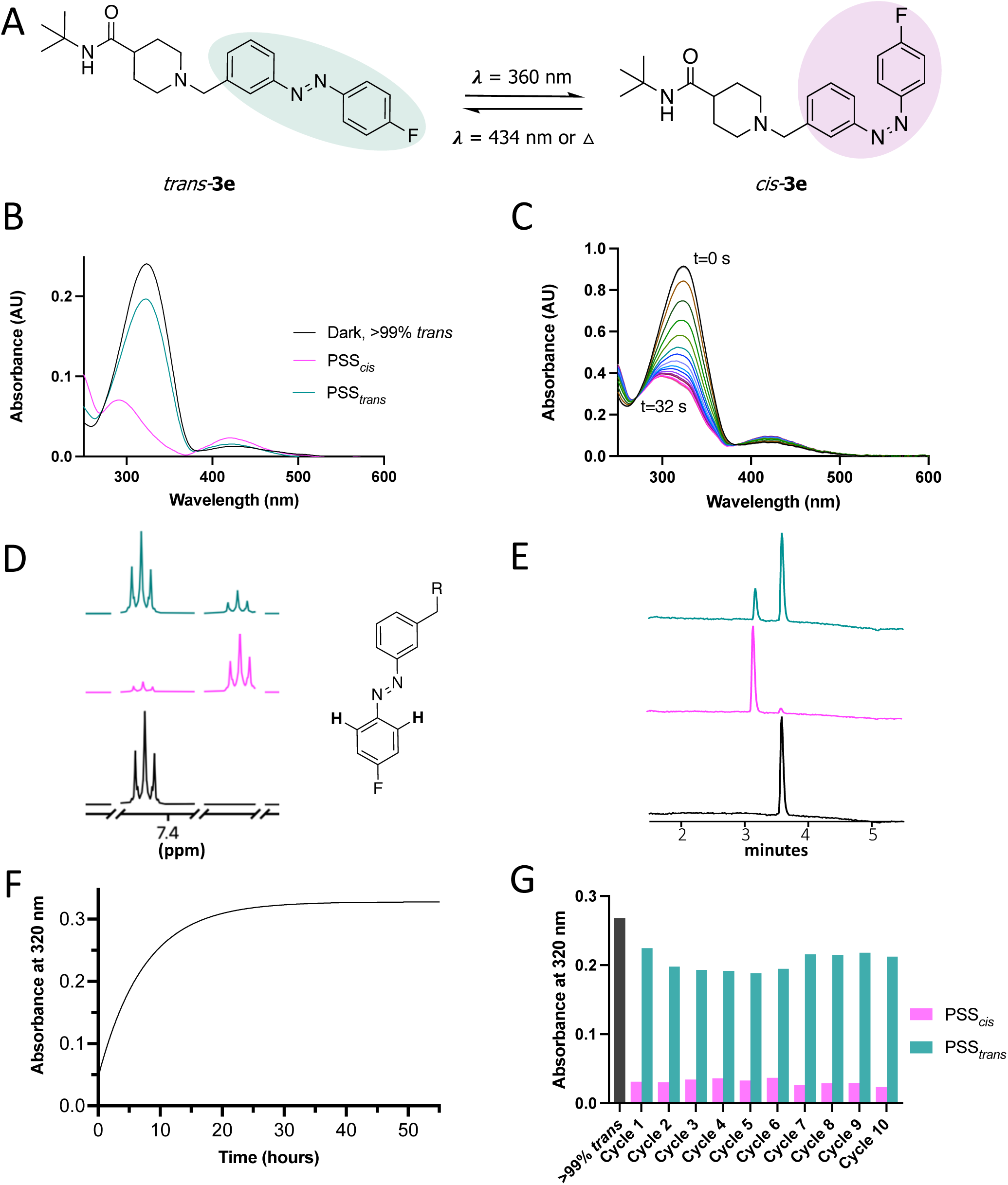
Photocharacterisation of key compound 3e. (A) Photoisomerization between the *trans* and *cis* isomers of **3e**. (B) UV-Vis spectra of **3e**. The black, magenta and green curves represent the absorption spectra (25 µM in HBSS buffer with 1 % DMSO at 25 °C) in the dark as >99% *trans* isomer, as PSS_*cis*_ after illumination of 5 min using 360 nm light, and as PSS*trans* after subsequent illumination of 5 min using 434 nm light, respectively. (C) Time-resolved UV/Vis absorption spectra of **3e** (50 µM in HBSS buffer with 1 % DMSO at 25 °C) upon illumination to PSS_*cis*_. Spectra were measured every 2 s for 32 s upon 360 nm illumination. (D) A representative part of ^1^H NMR spectra in d6-DMSO (50 mM) in the dark, under 360 nm light (illumination for 90 min) and 434 nm light (illumination for 90 min) at 25 °C. The presented peak belongs to the hydrogen atoms explicitly drawn in the structure accompanying the spectrum. Full spectra are available in Fig. S2. (E) LC-MS chromatograms belonging to a dark, PSS_*cis*_ or PSS*trans* sample (2.5 mM in DMSO at 25 °C ). Illumination with 360 nm (10 min) and 434 nm (10 min) at 25 °C was used. The detection was performed at 381 nm, i.e. the isosbestic point of **3e.** The full chromatograms with 254 nm detection are available in Fig. S3. (F) Thermal relaxation of **3e** at PSS_*cis*_ (25 µM in HBSS buffer with 1 % DMSO) at 60 °C measured by absorbance at 320 nm (the wavelength at which the difference in absorbance between *trans*-**3e** and *cis*-**3e** is maximal). UV. (G) Dynamic switching of **3e** (25 µM in HBSS buffer with 1 % DMSO at 25 °C) as measured by absorbance at 320 nm. UV-Vis spectra were obtained at 10-time intervals of 10 minutes each by alternating the irradiation wavelengths from 360 nm to 434 nm.

**Figure 3.**
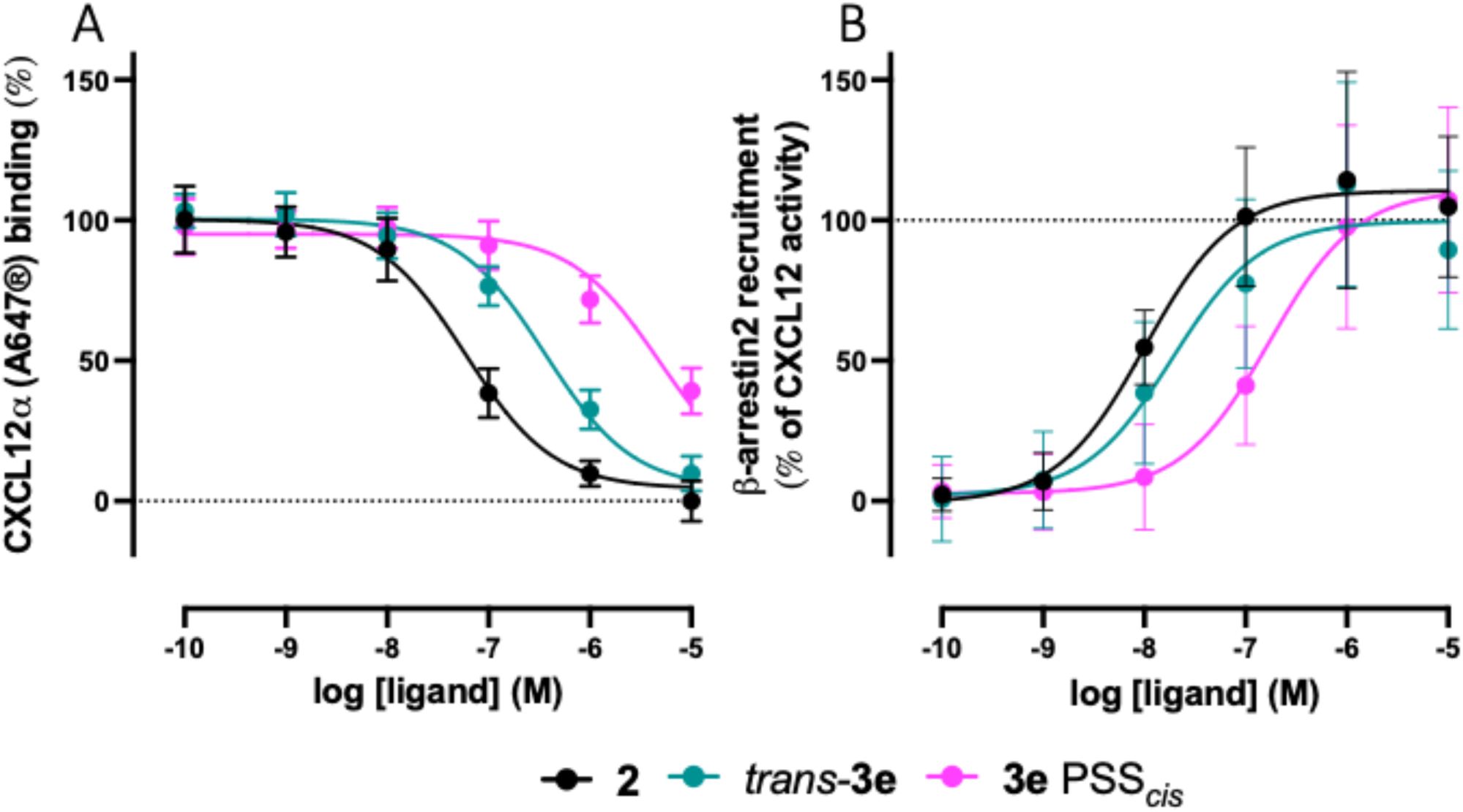
Pharmacological characterization of template compound 2 and photoswitchable key compound 3e. (A) HEK293T membranes transiently expressing NLuc-ACKR3 were used to measure concentration-dependent inhibition of fluorescently labeled CXCL12 (A647®) binding by **3e** (*trans* and PSS_*cis*_) and **2**. Data are shown as the BRET-ratio (BRET signal at >610 nm divided by NanoLuciferase signal at 460 nm). (B) β-arrestin2 recruitment to the ACKR3 in response to **3e** (*trans* and PSS_*cis*_) and **2** was measured using a NanoBiT complementation assay Data are normalized as (A) percentage of top and bottom plateau of the data for **2**, or (B) maximum luminescence induced by CXCL12 at 1 µM, which is defined as 100%. Data are shown as the mean ± SD of at least 3 independent experiments.

### 3.5. Pharmacology of key compound 3e

Template compound **2** and the two isomers of the key photoswitchable compound **3e** were pharmacologically characterized in more detail. To address any potential interfering compound precipitation, nephelometry was used to approximate the solubility of **2** and **3e** in HBSS buffer^48, 49^. The results (Fig. S5) show that **2**, *trans*-**3e** and **3e** at PSS_*cis*_ show no signs of aggregation up to 10^-5^ - 10^-4.5^, and also reflect the well-known difference in solubility between *trans*- and *cis*-azobenzenes ^1,2^.

The compounds were subsequently tested for their ability to induce β-arrestin2 recruitment to ACKR3 in a NanoBiT-based assay in living cells. In this assay, human ACKR3, C-terminally fused to SmBiT was transiently co-expressed in HEK293T-cells with N-terminally tagged LgBiT-β-arrestin2. Upon ACKR3 activation with CXCL12, β-arrestin2 is recruited to ACKR3 and the SmBiT and LgBiT protein fragments reconstitute a functional NanoLuciferase enzyme. In this assay CXCL12 induces β-arrestin2 recruitment to ACKR3 with a pEC50 value of 8.2 ± 0.3 (Fig. S6A). As expected, **3** also acts as a full agonist in this functional assay with a pEC50 value of 8.0 ± 0.1 and an intrinsic activity (α) of 1.1 ± 0.2 (Fig. 3B). The pEC50 value of **2** in this assay is approximately 20-fold lower than the reported potency (pEC50 = 9.3) in the patent that used the PathHunter™ β-arrestin2 recruitment assay^41^. As shown in Fig. 3B, both states of **3e** induced concentration-dependent β-arrestin2 recruitment to ACKR3 with similar intrinsic activity (α = 0.9 ± 0.3 and 1.1 ± 0.4, respectively) as **2** and CXCL12. The potency of *trans*-**3e** (pEC50 = 7.8 ± 0.2) to induce β-arrestin2 recruitment was 10-fold higher as compared to **3e** at PSS_*cis*_ (pEC50 = 6.8 ± 0.1), which is in line with the observed loss in binding affinity for ACKR3 upon *trans*-*cis* isomerization (Fig. 3A). Additionally, compound **3e** was tested in receptor trafficking assay to confirm ACKR3 activation and its subsequent internalisation. To this end, a BRET assay was used to measure the ligand-induced proximity between ACKR3 (NLuc-tagged) and the fluorescent protein mNeonGreen membrane-anchored via N-terminal palmitoylation/myristoylation signal of the Lyn protein. Upon stimulation with *trans*-**3e** (pEC50 = 8.2 ± 0.1). ACKR3 delocalizes from the membrane with 10-fold higher potency as compared to **3e** at PSS_*cis*_ (pEC50 = 7.2 ± 0.1) (Fig. S6B). Obtained results are in line with the observed loss of potency in β-arrestin2 recruitment.

### Pharmacological specificity of key compound 3e towards chemokine receptors

Verifying whether key compound **3e** (*trans* and PSS_*cis*_) is specific to ACKR3 is crucial information for future application of **3e** in studies related to ACKR3. The NanoBiT-based β-arrestin2 recruitment assay was employed to assess the specificity of **3e** for ACKR3 in comparison with a wide panel of chemokine receptors. This assay provides a functional readout to measure chemokine receptor activity. HEK293T cells transiently co-expressing β-arrestin2 N-terminally fused to LgBiT and the chemokine receptors C-terminally fused to SmBiT were used.^50^ (Fig. 4). In agonist mode, β-arrestin2 recruitment was measured upon treatment with 1 μM of compound **3e** (*trans* or PSS_*cis*_). As expected, **3e**-induced recruitment was observed for ACKR3. Yet, none was observed for any of the other tested chemokine receptors. This result indicates the high specificity of compound **3e** (*trans* or PSS_*cis*_) to activate ACKR3 (Fig. 4A). Additionally, potential antagonistic properties of compound **3e** (*trans* or PSS_*cis*_) were investigated. To assess this, chemokines activating each of the tested chemokine receptors were added after incubation with 1 μM of **3e** (*trans* or PSS_*cis*_). The antagonism of activated receptors by 1 μM of compound **3e** (*trans* or PSS_*cis*_) was less than 25% for all tested chemokine receptors (Fig. 4B). Thus, compound **3e** (*trans* or PSS_*cis*_) did not have a significant effect on chemokine receptors other than ACKR3.

**Figure 4.**
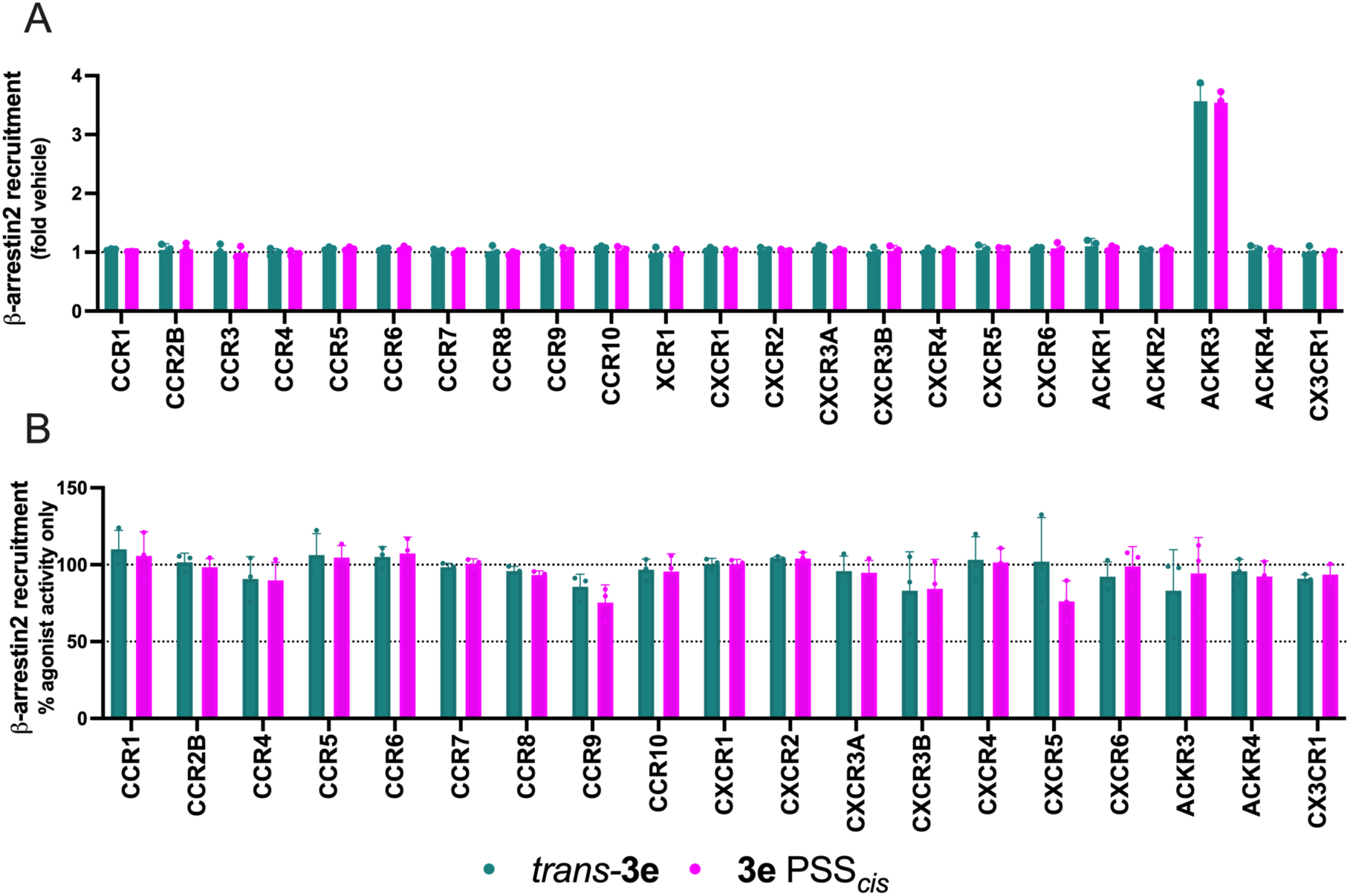
Characterization of pharmacological specificity of photoswitchable key compound 3e. HEK293T cells were transfected to co-express β-arrestin2 N-terminally fused to LgBiT and one of the chemokine receptors C-terminally fused to SmBiT. For CXCR4, β-arrestin2 was N-terminally fused to SmBiT and CXCR4 was C-terminally fused to LgBiT (A) β-arrestin2 recruitment to the indicated chemokine receptors in response to 1 µM of compound **3e** (*trans* or PSS_*cis*_). Data are shown as fold-increase in luminescence over the vehicle. (B) Evaluation of antagonistic properties of compound **3e** (*trans* and PSS_*cis*_). Ability of inhibition of agonist-induced β-arrestin2 recruitment to indicated chemokine receptors was determined after pre-stimulation with 1 µM of **3e** (*trans* or PSS_*cis*_). Data represents the signal obtained for wells stimulated with **3e** after treatment with the respective chemokine, normalized as % to the signal obtained for wells treated with endogenous agonist only (100 nM for CCR1/CCR9/CCR10, 50 nM for CXCR3B, 20 nM for other receptors). For the list of chemokines used as positive control for each receptor, see Supporting information. Data are shown as the mean ± SD of 3 independent experiments.

### 3.6. Characterization of the binding mode of 3e

The predicted binding modes of **2**, **3e** and selected derivatives were analyzed in light of the recently published cryo-EM structures of ACKR3 (PDB ID: 7SK4, 7SK9)^51^. A druggability assessment of the ACKR3 orthosteric pocket using GRID^53,54^ indicates that the ACKR3 binding pocket within the seven transmembrane domains (TMs) adapts remarkably to the bound ligand, changing size and shape to accommodate various ligands optimally. In both peptide and small-molecule complexes, two lipophilic hotspots can be identified in the orthosteric pocket, one in the major pocket (formed by TM3 to TM7) and one in the minor pocket (formed by TM1, TM2, TM3, and TM7). The lipophilic hotspot in the minor pocket, subpocket I (P I), is formed by side chains of Y51^1.39^, I97^2.57^, W100^2.60^, L104^2.64^, F124^3.32^, and L305^7.43^ (Ballesteros–Weinstein numbering in superscript)^55^. P I accommodates the CXCL12LRHQ His^2^ side chain residue (Fig. 5A, PDB ID: 7SK4), but interestingly this subpocket is not filled with any side chain of CXCL12WT. In the ACKR3-CCX662 structure (Fig. S7, PDB ID: 7SK9), P I shows induced fit, and a bigger lipophilic hotspot accommodates the quinoline moiety of CCX662 small molecule. The lipophilic hotspot in the major pocket, subpocket II (P II), is formed by L128^3.36^, F129^3.37^, I132^3.40^, S216^5.42^, W265^6.48^, Y268^6.51^, H269^6.52^ (Fig. 5A) and unoccupied by CXCL12WT side chains in the ACKR3-CXCL12WT structure, while Leu^0^ of the extended N-terminus of CXCL12LRHQ and the 4-hydroxypiperidine group of CCX662 moiety (Fig. S7) are reaching deeper in P II.

**Figure 5.**
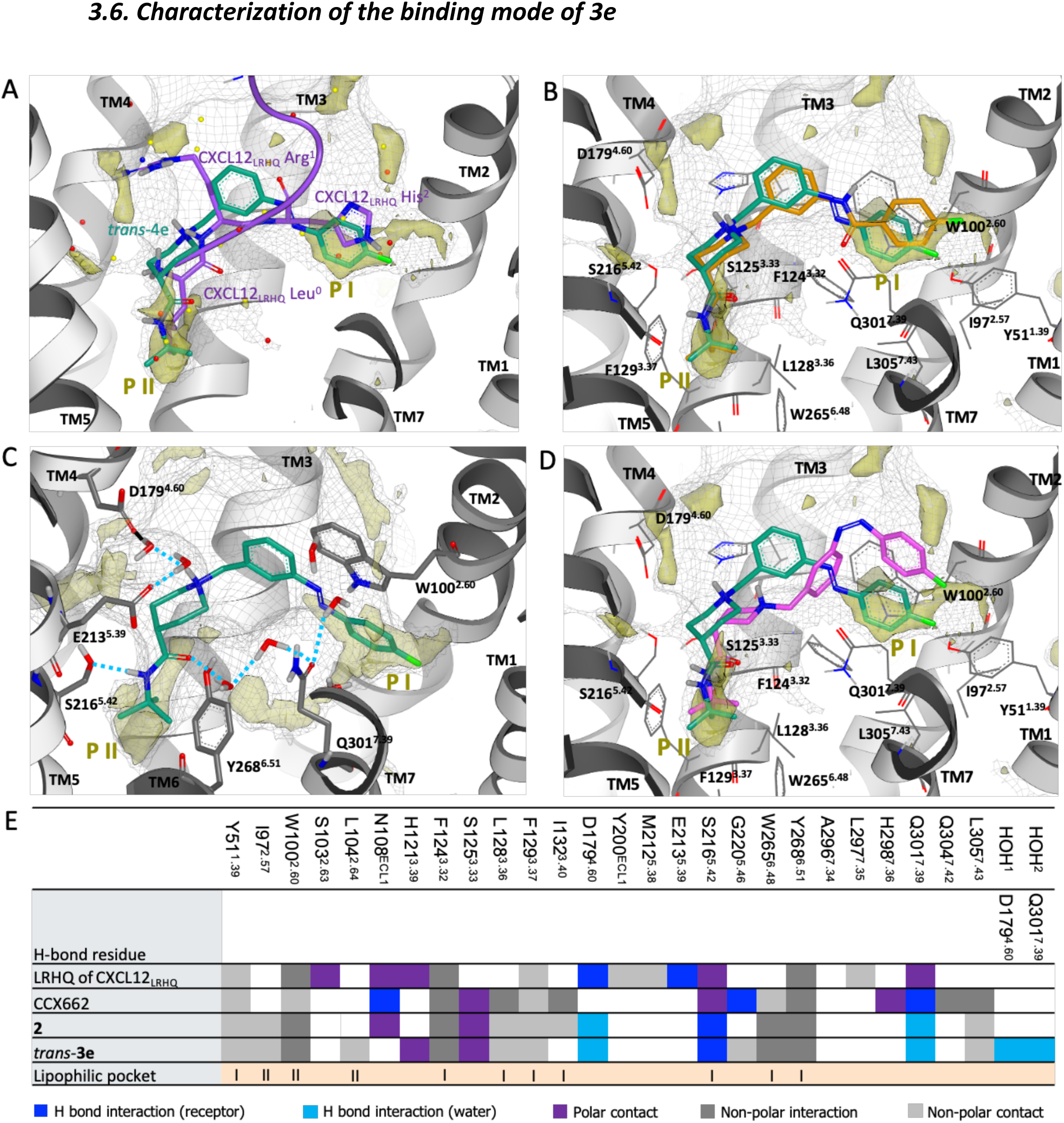
Overview of the binding mode of *trans-*3e in the ACKR3 binding pocket. (**A**) The binding site of ACKR3 (shown as grey cartoon) and CXCL12LRHQ N-terminus (shown in purple) with side chains Leu^0^, Arg^1^, His^2^ (depicted in stick representation in purple) as determined by cryo-EM (PDB ID: 7SK4)^51^. The docked pose of *trans-***3e** is shown in stick representation in green. In all figure panels, physicochemical properties of the binding sites characterized with GRID: GRID C3 surface to define the pocket surface in terms of how close a ligand carbon atom can reside (1.0 kcal mol^-1^; grey mesh) and C1= lipophilic hotspots (-2.8 kcal mol^-1^; yellow transparent solid surface) are shown. Key lipophilic hotspots, subpocket I and II, are labeled P I and P II, respectively. (**B**) Docking poses of **2** and *trans-***3e** (depicted in stick representation in orange and green, respectively) highlight overlapping binding orientations and occupation of key lipophilic hotspots in subpockets P I and P II by terminal moieties of the ligands. (**C**) The final trajectory frame from a 100 ns molecular dynamics simulation of a fully solvated membrane-embedded complex of ACKR3 and *trans*-**3e**. Hydrogen bonds are depicted in blue. (**D**) Docking poses of *trans-***3e** and *cis-***3e** (depicted in stick representation with green and pink, respectively) highlighting different binding orientations and occupation of key lipophilic hotspots in subpocket P I and P II by terminal moieties of the ligands. (**E**) Summary of interactions observed in ACKR3 structures in complex with CXCL12LRHQ (PDB ID: 7SK4) and CCX662 (PDB ID: 7SK9)^51^ and for docking poses of **2** and *trans-***3e** to the ACKR3-CXCL12LRHQ structure (PDB ID: 7SK4), including key water molecules from predicted water networks in the binding site. Subpockets I and II (labeled P I and P II) are defined, and interactions are represented according to the different interaction types. Hydrogen bond interactions with the receptor or with water molecules are shown in dark and light blue, respectively. Polar contacts are shown in purple, while non-polar interactions or contacts are depicted in dark and light grey, respectively. The figure was generated with Vida (OpenEye).^52^

Docking poses of **2** and *trans-***3e** to the ACKR3-CXCL12LRHQ (PDB ID: 7SK4) orthosteric binding site show an extended conformation reaching both subpockets, favored by the amide piperidine conformation (Fig. 5B). The fluorobenzene moiety of *trans-***3e** and **2** occupies P I, while their terminal *N*-*tert*-butyl moiety fills P II, resembling the shape of the backbone of the bound CXCL12LRHQ N-terminus, and as expected, in overlapping orientations for *trans-***3e** and **2** (Fig. 5B). To validate the docking poses of *trans-***3e**, the complex of ACKR3-*trans-***3e** was also subjected to 100 ns of MD (molecular dynamics) simulations in a fully solvated membrane-embedded system. The MD results indicate that only one direct hydrogen bond interaction was formed between the docked pose of the ligand and the receptor, namely between the amide N-H moiety and S216^5.42^ (Fig. 5C). Predicted water networks in the binding site of the complex highlight water-mediated interactions between the ligand and polar side chains in the binding site, including D179^4.60^ and Q301^7.39^, in addition to favorable van der Waals interactions with apolar side chains (Fig. 5C and S8). In P I, W100^2.60^ is anchoring the azosteric amide moiety of **2** and the azobenzene of *trans-***3e** via ν-ν stacking (Fig. 5C), similarly to the ν-ν stacking observed for CXCL12LRHQ His^2^. The MD simulations suggest that the predicted binding mode (Fig.5B) is stable. This is illustrated by the key hydrogen bond interactions (direct or water-mediated ones) being maintained over the 100 ns of simulation (Fig.S8 and 5C).

Docking poses and ligand–receptor interactions for *trans-***3e** (pEC50 = 7.8 ± 0.2) were supported by SAR of selected analogues (Table 1). For example, compound *trans-***3f** (pEC50 = NA), with a *para*-substituted inner ring of the azobenzene moiety instead of a *meta* one, cannot adopt a conformation to reach P I (Fig. S9), indeed losing the ability to bind ACKR3. The bulkier substituent of *trans-***3g** (pEC50 = NA) at the terminal aryl ring would not fit optimally in P I, clashing with Y51^1.39^ or losing the optimal ligand conformation to fit P I (Fig. S9). Importantly, *cis-***3e** was docked in the ACKR3 binding site and subsequently, subjected to MD simulations. Differently from *trans-***3e**, the non-planarity of the azobenzene moiety of the *cis* isomer does not allow the compound to fill the key lipophilic hotspot in P I (Fig. 5D), contributing to a weaker binding of the ligand in the binding site (Table 1) and fluctuations of the ligand orientation throughout MD simulations (Fig. S10).

A final summary of the various interactions of the four N-terminal residues (LRHQ) of CXCL12LRHQ analog, CCX662, **2** and *trans*-**3e** with key amino acids and (predicted) water molecules in the ACKR3 binding pocket is given in Fig. 5E. While all these ligand chemotypes occupy the subpockets P I and P II, they nonetheless exhibit unique interaction patterns as derived from the published cryo-EM studies and our modelling studies with **2** and *trans*-**3e**.

These data are either based on the published cryo-EM structures (CXCL12LRHQ analog and CCX662^51^) or on the final binding modes of **2** and *trans*-**3e** (this study). The comparison highlights unique interaction patterns for the various ligands, despite the fact that all occupy the same subpockets P I and P II.

## Conclusion

This study describes the design and synthesis of the first photoswitchable ligand for the atypical chemokine receptor ACKR3. A library of azobenzene analogs based on the azologization of a small-molecule ACKR3 agonist from patent literature was synthesized and characterized photochemically and pharmacologically. The key photoswitchable agonist **3e** offers a 10-fold decrease in both the inhibition of CXCL12 binding and stimulation of β-arrestin2 recruitment upon illumination with 360 nm. Compound **3e** exhibits selectivity for ACKR3 within a large panel of chemokine receptors. The recently published cryo-EM structure of ACKR3 was used to propose a binding mode of **3e** and to explain the observed SAR. Compound **3e** (VUF25471) is the first photoswitchable ligand for an intrinsically β-arrestin-biased GPCR and might be a valuable photopharmacology tool to investigate the spatial and temporal effects of ACKR3 signaling in biological processes such as cell migration and cancer development.

## ASSOCIATED CONTENT

Experimental procedures and additional figures for photochemical, physicochemical, pharmacological and modeling studies. Experimental details and NMR analyses for the synthesis of all studied compounds.

## AUTHOR INFORMATION

### Boomi Author

- **Maikel Wijtmans** - *Division of Medicinal Chemistry, Amsterdam Institute of Molecular and Life Sciences, Faculty of Science, Vrije Universiteit Amsterdam, De Boelelaan 1108, 1081 HZ Amsterdam, The Netherlands*;
- **Rob Leurs** - *Division of Medicinal Chemistry, Amsterdam Institute of Molecular and Life Sciences, Faculty of Science, Vrije Universiteit Amsterdam, De Boelelaan 1108, 1081 HZ Amsterdam, The Netherlands*; http://orcid.org/0000-0003-1354-2848;

### Author Contributions

- **Sophie Bérenger** - *Division of Medicinal Chemistry, Amsterdam Institute of Molecular and Life Sciences, Faculty of Science, Vrije Universiteit Amsterdam, De Boelelaan 1108, 1081 HZ Amsterdam, The Netherlands*
- **Justyna M. Adamska** - *Division of Medicinal Chemistry, Amsterdam Institute of Molecular and Life Sciences, Faculty of Science, Vrije Universiteit Amsterdam, De Boelelaan 1108, 1081 HZ Amsterdam, The Netherland*
- **Francesca Deflorian** – Sosei Heptares, Cambridge CB21 6DG, U.K.
- **Henry F. Vischer** - *Division of Medicinal Chemistry, Amsterdam Institute of Molecular and Life Sciences, Faculty of Science, Vrije Universiteit Amsterdam, De Boelelaan 1108, 1081 HZ Amsterdam, The Netherlands*
- **Chris de Graaf** – Sosei Heptares, Cambridge CB21 6DG, U.K.
- **Barbara Zarzycka** - *Division of Medicinal Chemistry, Amsterdam Institute of Molecular and Life Sciences, Faculty of Science, Vrije Universiteit Amsterdam, De Boelelaan 1108, 1081 HZ Amsterdam, The Netherlands*
- **Iwan J. P. de Esch** - *Division of Medicinal Chemistry, Amsterdam Institute of Molecular and Life Sciences, Faculty of Science, Vrije Universiteit Amsterdam, De Boelelaan 1108, 1081 HZ Amsterdam, The Netherlands*
- **Christie B. Palmer** - *Immuno-Pharmacology and Interactomics, Department of Infection and Immunity, Luxembourg Institute of Health, 29, rue Henri Koch, L-4354, Esch-sur-Alzette, Luxembourg*.
- **Martyna Szpakowska** - *Immuno-Pharmacology and Interactomics, Department of Infection and Immunity, Luxembourg Institute of Health, 29, rue Henri Koch, L-4354, Esch-sur-Alzette, Luxembourg*.
- **Andy Chevigné** - *Immuno-Pharmacology and Interactomics, Department of Infection and Immunity, Luxembourg Institute of Health, 29, rue Henri Koch, L-4354, Esch-sur-Alzette, Luxembourg*.

### FUNDING SOURCES

This research was funded by European Union’s Horizon2020 Marie Skłodowska-Curie Actions (MSCA) Program under Grant Agreement 860229 (ONCORNET 2.0). This study was supported by the Luxembourg Institute of Health (LIH) (NanoLux Platform), Luxembourg National Research Fund (INTER/FNRS grants 20/15084569 and AFR Opiokine 20/14616593), F.R.S.-FNRS-Télévie (grants 7.4593.19, 7.4529.19,

7.8504.20, 7.4502.21 and 7.8508.22)

## ACKNOWLEDGMENT

We thank Hans Custers for recording HRMS spectra, Elwin Janssen for assistance with NMR analysis and Icaro Simon, Rick Riemens and Kobus Versteegh for helpful discussions.

## ABBREVIATIONS

Α: intrinsic activity
Å: Ångström
ACKR3: atypical chemokine receptor 3
cryoEM: cryogenic electron microscopy
DCM: dichloromethane
DIBAL-H: diisobutylaluminium hydride
DMF: dimethylformamide
DMSO: dimethyl sulfoxide
EDG: electron donating group
EWG: electron withdrawing group
ERK: extracellular signal regulated kinase
GPCR: G protein-coupled receptor
HBSS: Hanks’ Balanced Salt solution
HRMS: high-resolution mass spectrometry
LC-MS: liquid chromatography–mass spectrometry
MDCK: Madin-Darby Canine Kidney
NanoBRET: Nano Bioluminescence Resonance Energy Transfer
NMR: nuclear magnetic resonance
On: overnight
PCC: pyridinium chlorochromate
Pd/C: palladium on carbon
PSS: photostationary state
IFP: interaction fingerprint
MD: molecular dynamics
Rt: room temperature
rVSMCS: rat vascular smooth muscle cell
SAR: structure–activity relationship
satd Aq: saturated aqueous
SD: standard deviation
SEM: standard error of mean
TEA: triethylamine
TFA: trifluoroacetic acid
TM: transmembrane
WT: wild type

## Notes

### Competing Interest Statement

The authors have declared no competing interest.

