## Supplementary Material for "Design, synthesis and pharmacological characterization of the first photoswitchable small-molecule agonist for the Atypical Chemokine Receptor 3"

### Table of contents


| | |
|---|---|
| Page S3 | UV-Vis spectra of **3b-c, 3e-l** |
| Page S4 | Extended $^{1}$H-NMR spectra for illuminations of **3e** |
| Page S5 | Extended LC-MS chromatograms for illuminations of **3e** |
| Page S6 | Thermal relaxation of **3e** |
| Page S7 | Nephelometry of **2**, *trans*-**3e**, and **3e** at $PSS_{cis}$ |
| Page S8 | β-arrestin2 recruitment and ACKR3 internalization in response to CXCL12 and photoswitchable key compound **3e** |
| Page S9 | Supplementary Table 1. Positive controls used in Figure 4 |
| Page S10 | Overlap of ACKR3-CCX662 and ACKR3-$CXCL12_{LRHQ}$ binding pocket with *trans*-**3e** predicted pose |
| Page S11 | 2D interaction plot of *trans*-**3e** |
| Page S12 | SAR validation of *trans*-**3e** predicted pose |
| Page S13 | MD simulation of *cis*-**3e** |
| Page S14 | Photochemistry procedures |
| Page S15 | Molecular modelling procedures |
| Page S16 | Pharmacology procedures |
| Page S19 | Synthesis procedures |
| Page S34 | Chemical analyses |
| Page S58 | References |

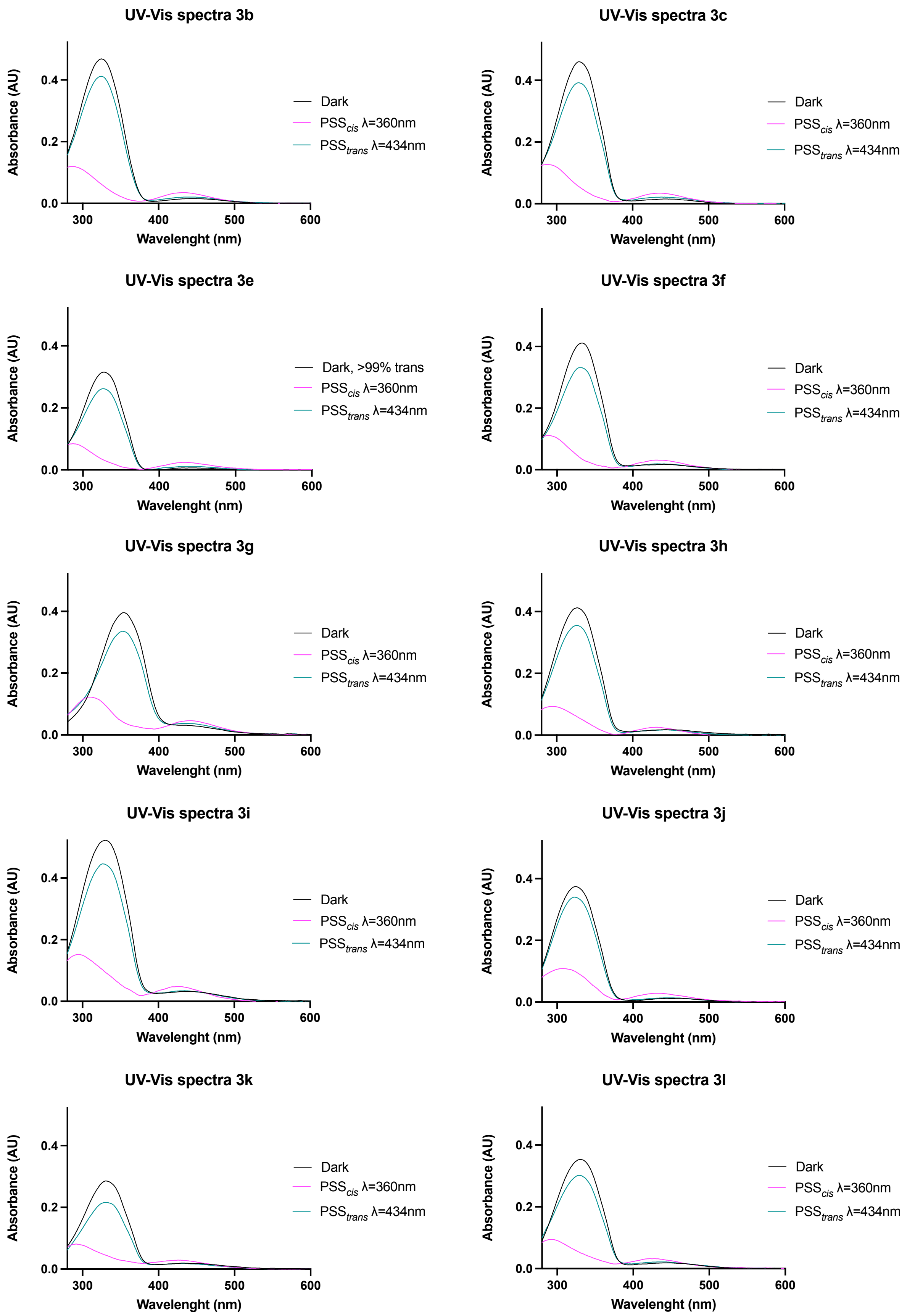


**Figure S1: Absorbance spectra for 3b-c,e-l**. Spectra were recorded for 25 µM of **3b-c,e-l** in DMSO in the dark, after 360 ± 20 nm illumination for 10 min and after subsequent 434 ± 9 nm illumination for 10 min at room temperature. Compounds **3a** and **3d** are omitted because of photodecomposition.

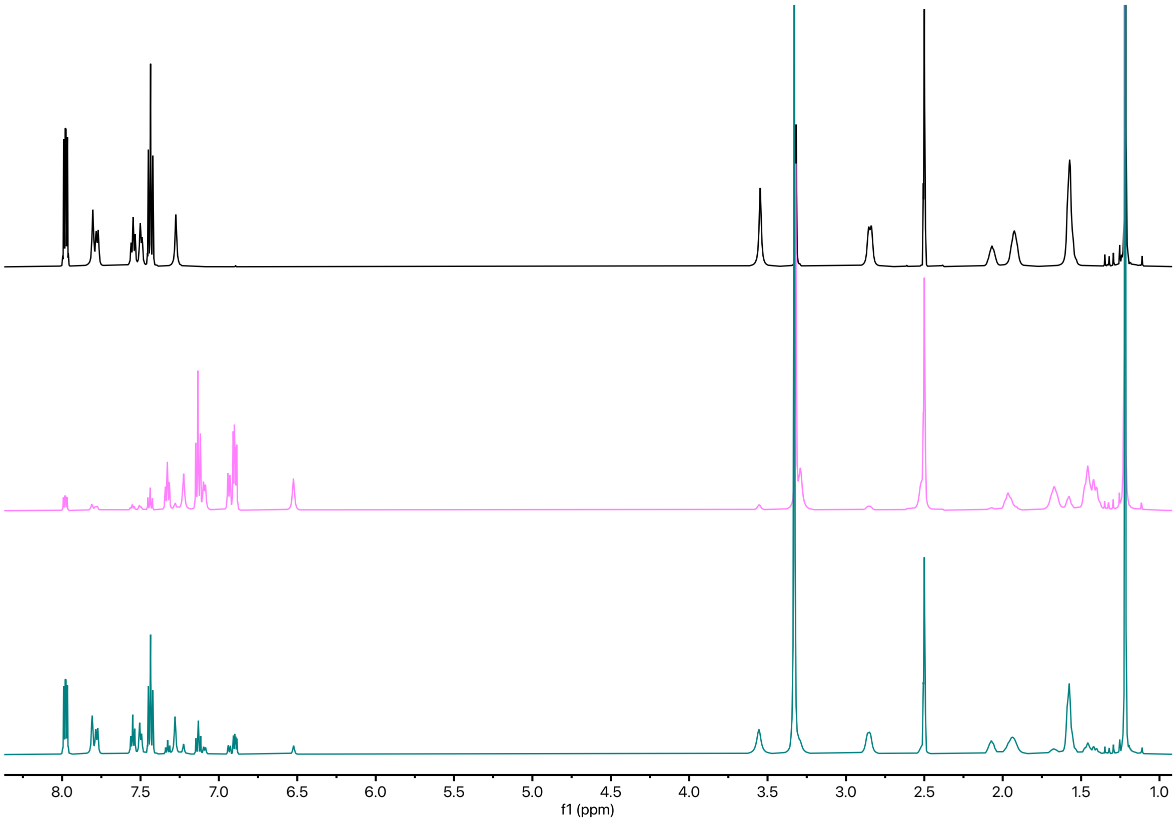


**Figure S2: Full $^{1}$H NMR spectra for the irradiations shown in Fig. 2D.** The irradiations were performed with a solution of 50 mM of **3e** in DMSO-d6. The black spectrum represents the compound as 99% *trans*. The magenta spectrum represents the mixture after 90 min illumination at 360 nm (the sample was vortexed every 10 min). The green spectrum represents the mixture after 90 min illumination at 360 nm (the sample was vortexed every 10 min).

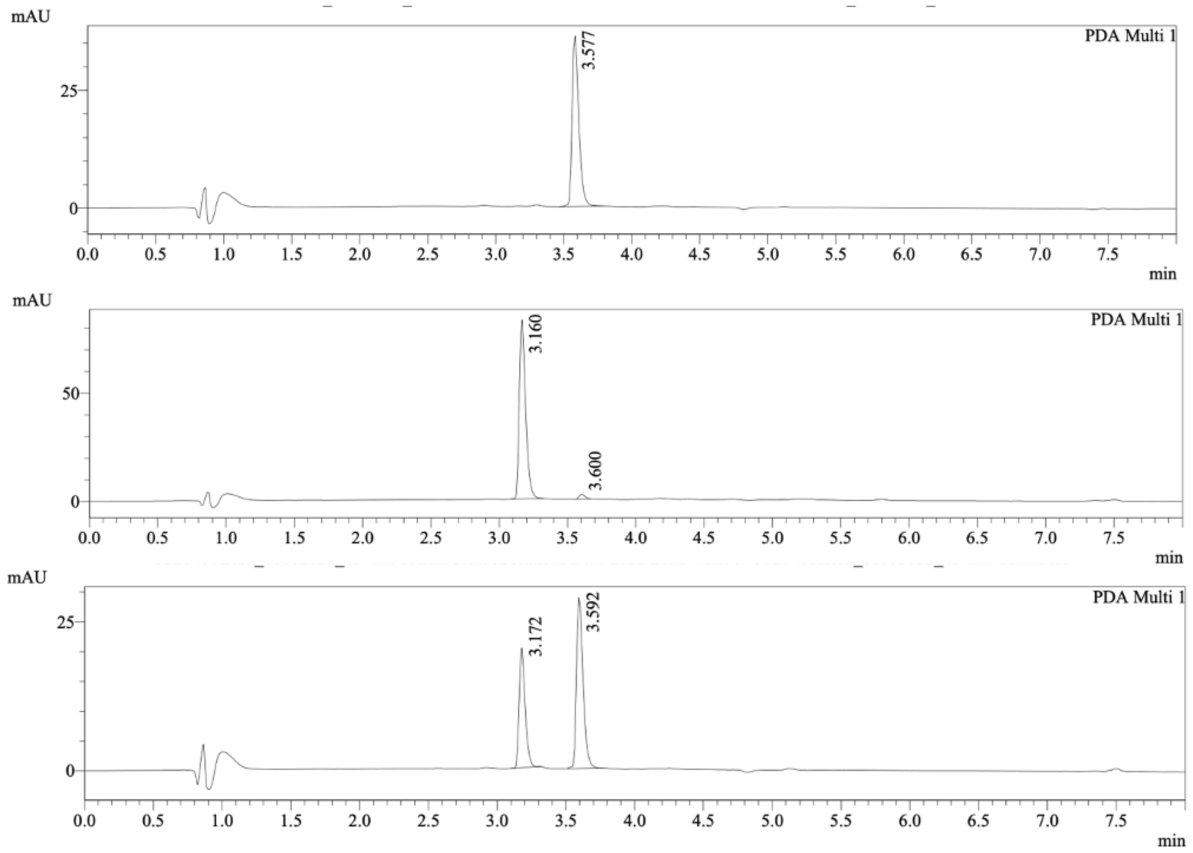


**Figure S3: Full chromatograms for the irradiations shown in Fig. 2E, but with detection at 254 nm.** The first chromatogram is the dark sample (99% *trans*), followed by that for the $PSS_{cis}$ and the $PSS_{trans}$ sample, respectively The irradiations were performed with a solution of 2.5 mM of **3e** in DMSO for 10 min. The LCMS samples measured were diluted to 2.5 μM with can prior to LC analysis

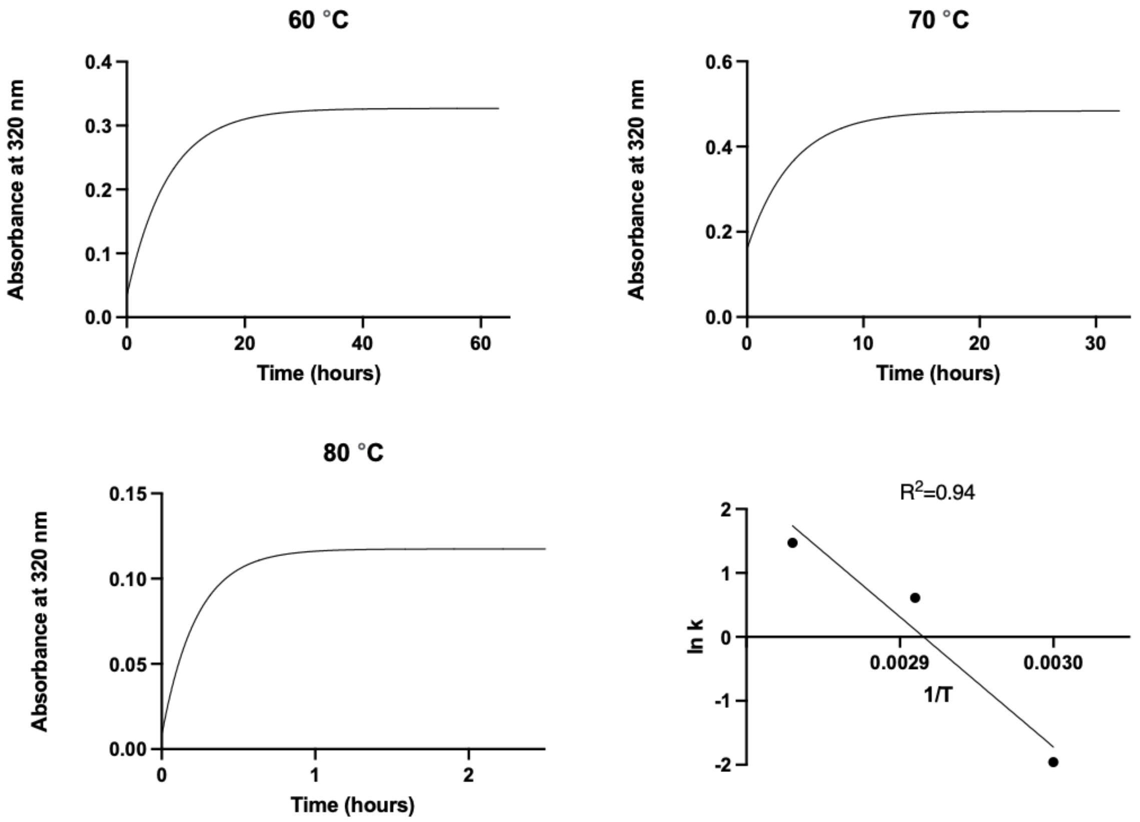


**Figure S4**. **Arrhenius plot for the relaxation of the PSS$_{cis}$ state of 3e**. Relaxation was measured at a concentration of 25 µM in HBSS buffer with 1% DMSO. The value provided in the main text is an extrapolation of the linear fit presented in this plot. The $R^2$ value for linear fit is 0.94.

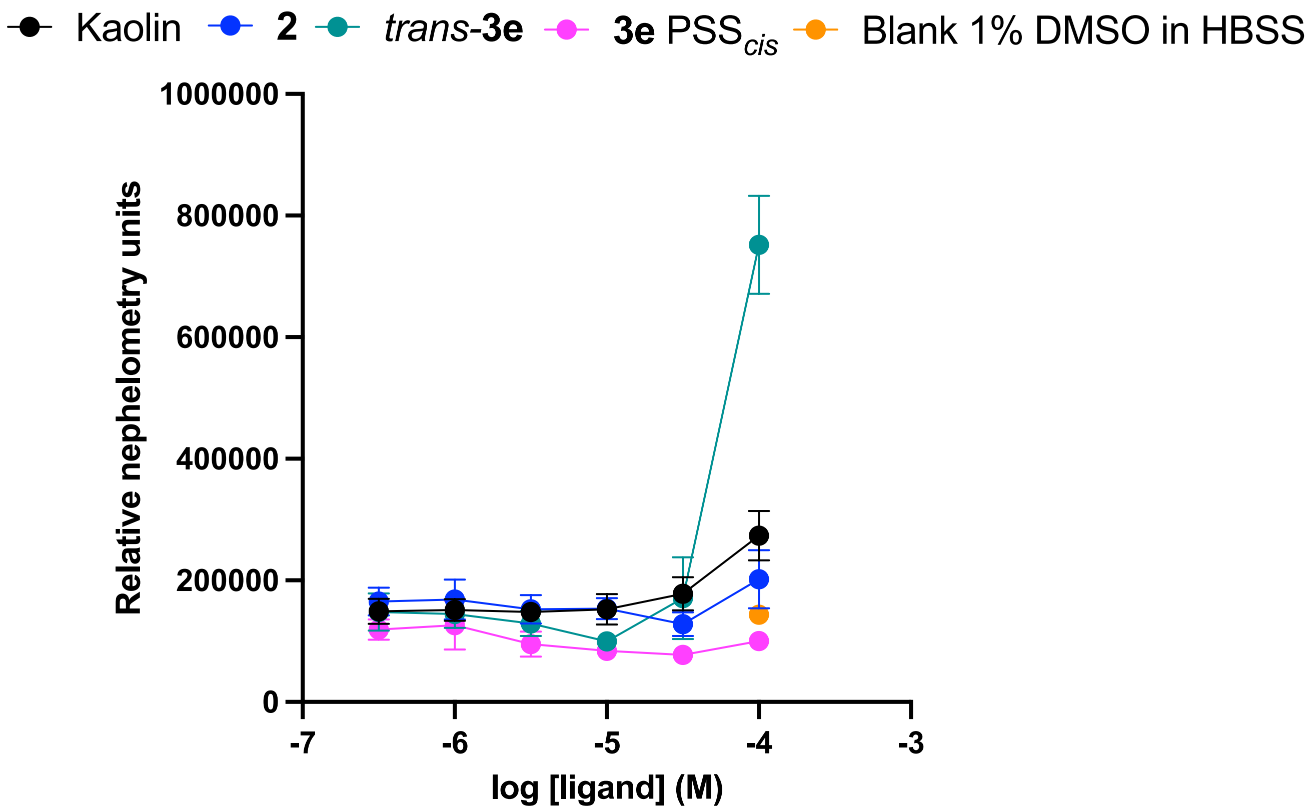


**Figure S5. Nephelometry measurements of compounds 2, *trans*-3e and 3e at *PSS$_{cis}$* in HBSS buffer + 1 % DMSO**. Kaolin was used as a positive control. Three experiments were performed in triplicate and data are shown as mean ± SD.

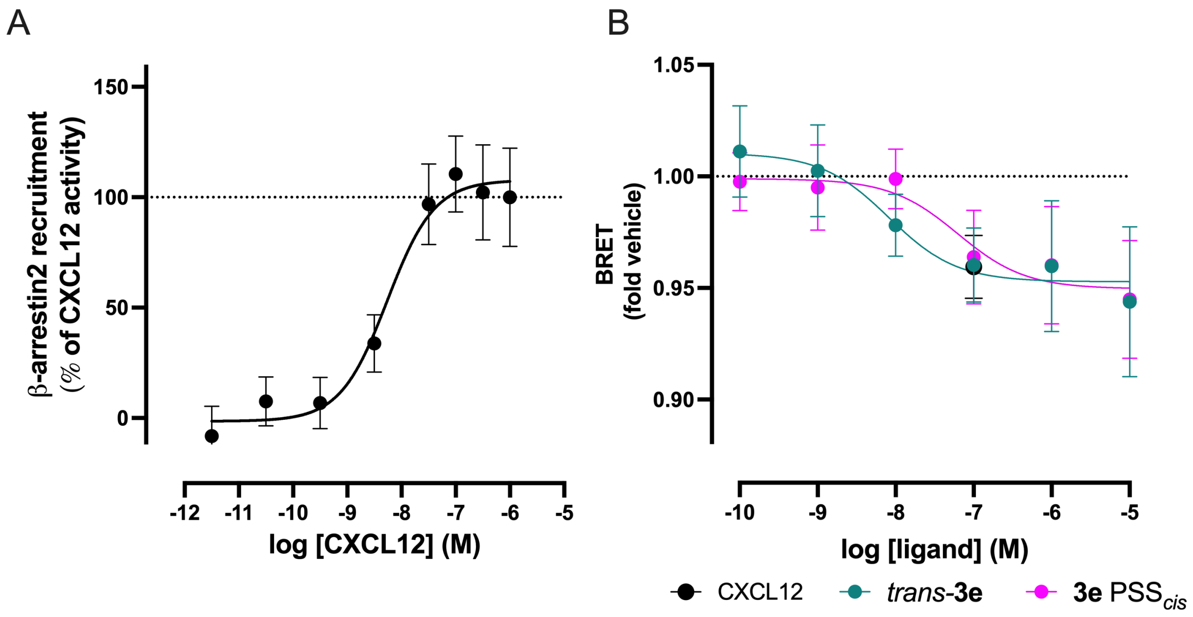


**Figure S6. β-arrestin2 recruitment and ACKR3 internalization in response to CXCL12 and photoswitchable key compound 3e**.(A) CXCL12 induced β-arrestin2 recruitment to the ACKR3. Recruitment was measured using a NanoBit complementation assay. Data are normalized to the maximum luminescence induced by CXCL12 at 1 µM, defined as 100%. Data are shown as the mean ± SD of at least 3 independent experiments. (B) ACKR3 internalization upon stimulation with CXCL12 and photoswitchable key compound **3e.** The colocalization between ACKR3-NLuc and Lyn-NeonGreen by increasing concentrations of **3e** (*trans* and $PSS_{cis}$) and 100 nM of CXCL12 was determined by BRET in HEK293T cells. Data are shown as fold increase in BRET ratio over the vehicle. Data are shown as the mean ± SD of at least 3 independent experiments.

**Supplementary Table 1. Positive controls used in Figure 4**

| Receptor | Positive control |
|---|---|
| CCR1 | CCL3 |
| CCR2B | CCL2 |
| CCR3 | CCL13 |
| CCR4 | CCL22 |
| CCR5 | CCL5 |
| CCR6 | CCL20 |
| CCR7 | CCL19 |
| CCR8 | CCL1 |
| CCR9 | CCL25 |
| CCR10 | CCL27 |
| XCR1 | XCL2 |
| CXCR1 | CXCL8 |
| CXCR2 | CXCL8 |
| CXCR3A | CXCL11 |
| CXCR3B | CXCL11 |
| CXCR4 | CXCL12 |
| CXCR5 | CXCL13 |
| CXCR6 | CXCL16 |
| ACKR1 | CXCL5 |
| ACKR2 | CCL5 |
| ACKR3 | CXCL12 |
| ACKR4 | CCL19 |
| CX3CR1 | CX3CL1 |

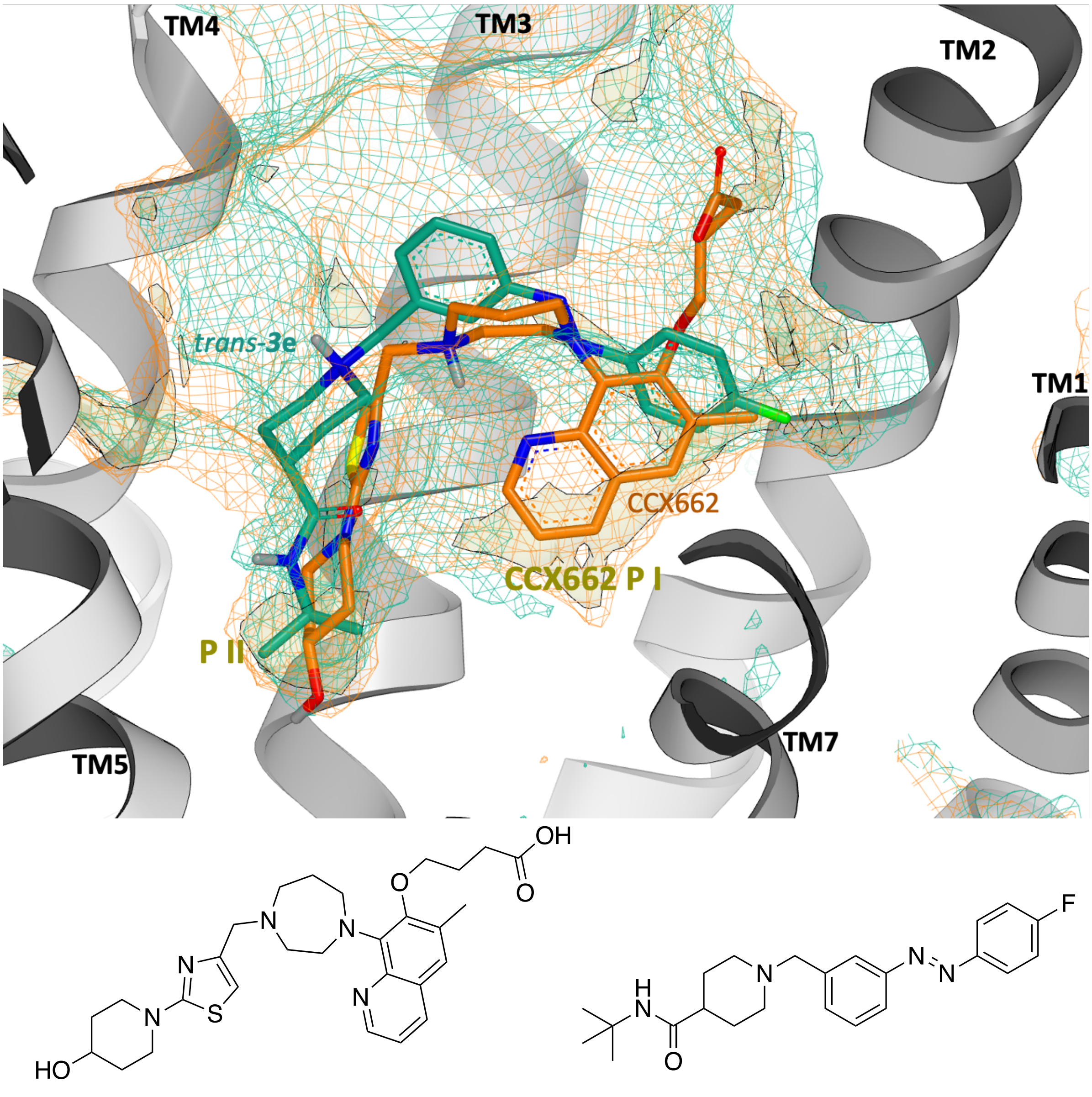


CCX662 *trans*-**3e**

**Figure S7. Overlap of ACKR3-CCX662 and ACKR3-CXCL12$_{LRHQ}$ binding poses with the predicted binding pose of *trans*-3e**. Comparison between ACKR3-CCX662 induced-fit subpocket I (labelled CCX662 PI, GRID C3 mesh colored in orange) and ACKR3-CXCL12$_{LRHQ}$ binding site (GRID C3 mesh colored in cyan). For clarity, only the ACKR3-CCX662 structure (PDB ID 7SK9) is shown (depicted in grey cartoon representation). Docked pose of *trans*-**3e** and CCX662 from cryo-EM structure 7SK9 are shown in the binding site (depicted in stick representation with carbon atoms colored in green and orange, respectively). Physicochemical properties of the binding sites characterized with GRID: GRID C3 surface to define the pocket surface in terms of how close a ligand carbon atom can reside (1.0 kcal mol$^{-1}$; mesh) and C1= lipophilic hotspots (-2.8 kcal mol$^{-1}$; yellow transparent solid) are shown. Key lipophilic hotspots, subpocket I and II, are labeled PI and PII, respectively. The 2D structures of CCX662 and *trans*-**3e** are also shown.

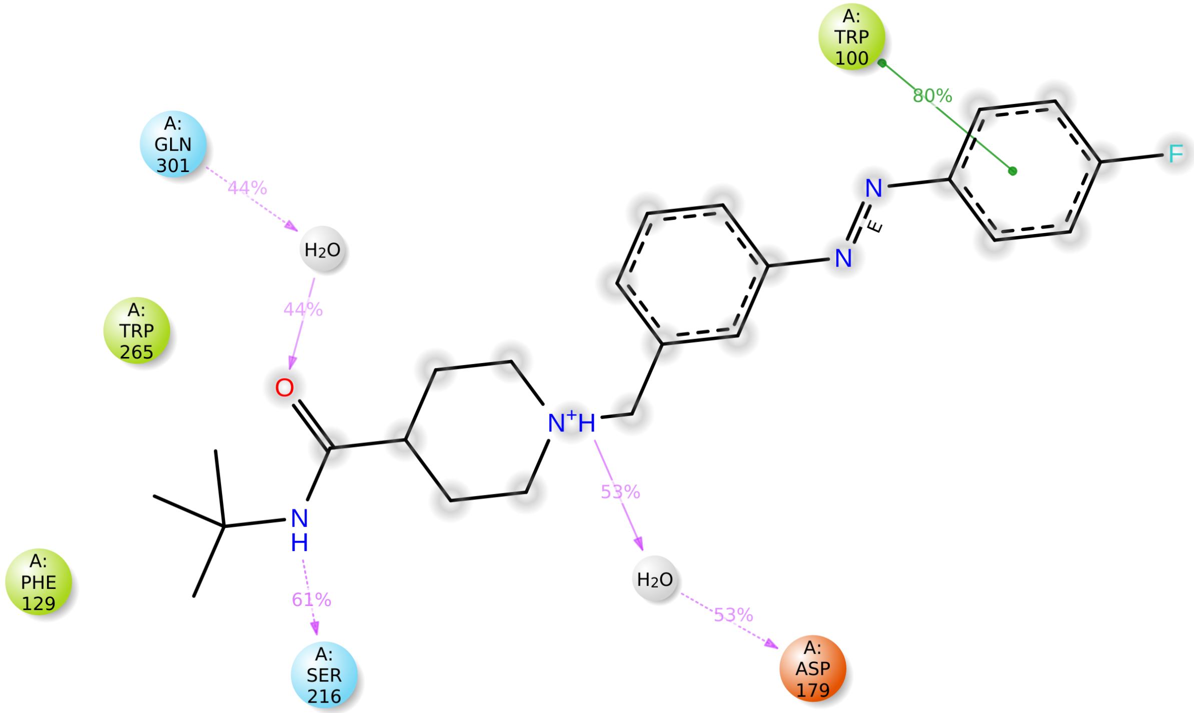


**Figure S8. 2D interaction plot of *trans*-3e.** Ligand-protein contacts during MD simulations: interactions of atoms of *trans*-**3e** with the protein residues with occurrence shown as percentage of the simulation time in the trajectory. Residues involved in hydrophobic contacts, polar interactions and ionic interactions are shown in green, blue and orange, respectively. Green lines represent π−π stacking, and pink lines represent hydrogen bonds. Dashed lines correspond to side-chain interactions. The percentage reported reflects the contact strength along the simulation. The grey spheres correspond to the water network involved in the interactions. The figure was made with Maestro (Schrodinger).[1]

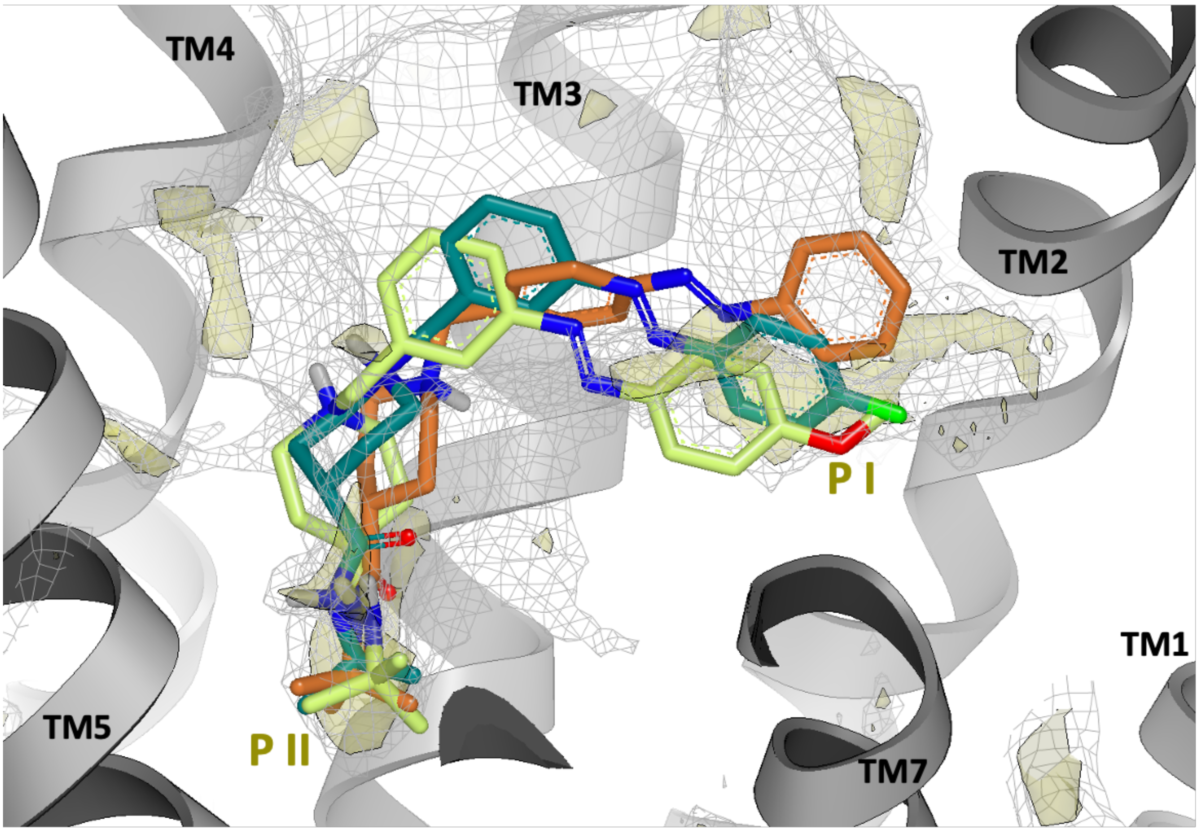


**Figure S9**. **SAR validation of the predicted pose for *trans*-3e.** Overlap of the docking poses of *trans*-**3e** (green carbon atoms), *trans*-**3f** (orange carbon atoms) and *trans*-**3g** (yellow carbon atoms) to ACKR3 binding site of 7SK4 structure (grey cartoon). GRID C3 surface to define the pocket surface in terms of how close a ligand carbon atom can reside (1.0 kcal mol-1; grey mesh) and C1= lipophilic hotspots (-2.8 kcal mol-1; yellow transparent solid) are shown. Key lipophilic hotspots, subpocket I and II, are labeled P I and P II, respectively. The figure was made with Vida (OpenEye).[2]

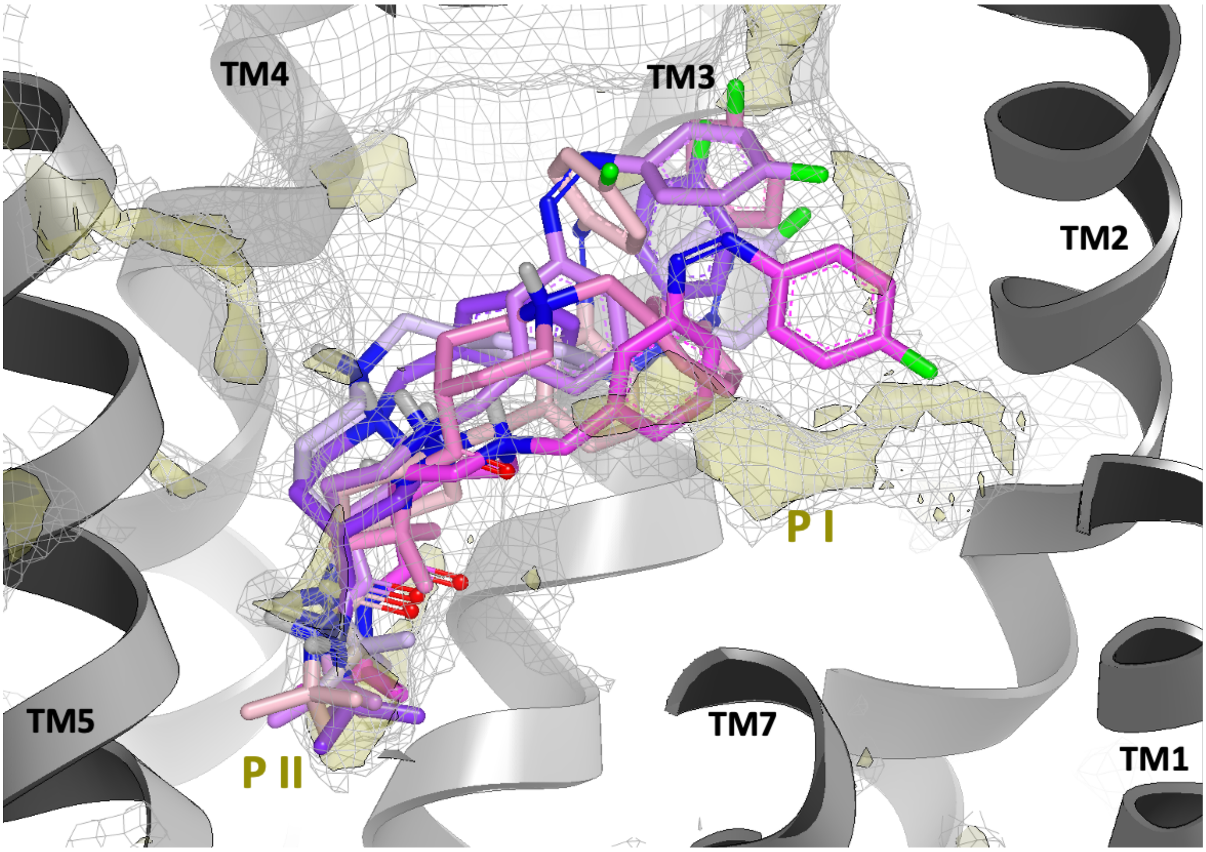


**Figure S10**. **MD simulation of *cis*-3e.** Selected frames from the 100 ns molecular dynamics simulation trajectories of the complex of ACKR3 and *cis*-**3e** (colored from magenta (initial frame) to light pink (last frame). The receptor depiction from the last frame is shown in grey cartoon representation. Key lipophilic hotspots, subpocket I and II, are labeled P I and P II, respectively. The figure was made with Vida (OpenEye).[2]

### Photochemistry

UV−Vis spectra were recorded using a Thermo-scientific Evolution 201 PC spectrophotometer equipped with a thermostated cell holder set at 20 °C. Fits of UV−vis spectroscopy data were generated using Prism 9.4.1. Illumination was executed using a Sutter instruments Lambda LS with a 300 W full-spectrum lamp connected to a Sutter instruments Lambda 10-3 optical filter changer equipped with 434 ± 9 nm and 360 ± 20 nm filters. The light intensity used in all illuminations is 0.77 $mW/mm^2$ using the 360 ± 20 nm filter and 0.57 $mW/mm^2$ for the 434 ± 9 nm filter as measured using a Thorlabs PM16−401 power meter. For photochemical analyses, illuminations were performed in Hellma Suprasil quartz 114QS cuvettes. Thermal relaxation experiments and Arrhenius extrapolations were performed according to Priimagi et al[3], using a compound concentration of 25 µM in HBSS +1% DMSO and temperatures of 60, 70, and 80 °C. Illuminations for pharmacological experiments were performed in cylindrical clear glass vials with a volume of 4.5 mL. The typical distance between light source and vial or cuvette was 2 cm. For Fig. 2C, a different Sutter instrument was used. This Lambda 721 lamp is connected to a Sutter instruments Lambda 10-3 optical filter changer equipped with 434 ± 28 nm and 365 ± 12 nm filters. The light intensity used for the experiments with this lamp is 200 mW using the 365 ± 12 nm filter and 560 mW for the 440 ± 28 nm filter as indicated by the supplier. This lamp allows for continuous measurement with an integration time of 1.000000 $\cdot 10^{-1}$ second.

### Molecular modelling

Cryo-EM structures of ACKR3 in complex with CXCL12 (wild-type and variant) and the small-molecule agonist CCX662 were downloaded from the Protein Data Bank (PDB IDs: 7SK3, 7SK4, 7SK5, 7SK6, 7SK7, 7SK8, 7SK9).[4] Proteins were prepared with Protein Preparation Wizard in Maestro (Schrödinger release 2022-1) with default settings. Missing side chains and hydrogen atoms were added, alternative main-chain conformations were kept, residues protonation states were assigned with PROPKA at pH 7.0, water orientation and hydrogen bond network were optimised, and a brief minimisation was performed to a root-mean-square deviation (RMSD) of up to 0.5 Å using the OPLS4 force field. WaterMap[5] was used to calculate the locations and thermodynamic properties of water molecules in the pseudo-apo binding sites for druggability assessment. Physicochemical properties of the binding site in the different structures were analysed using GRID.[6,7] Ligands were prepared using MoKa (Molecular Discovery), [8] for the generation of tautomeric and ionic states and Corina (MN-AM)[9] for low-energy 3D conformations. Molecular docking was carried out with Glide SP (Schrodinger release 2022-1) [10,11] starting from the Corina-generated conformations. Ligand conformational sampling was carried out using MacroModel Monte Carlo plus Minimization (MCMM) mode, using default parameters with the OPLS4 force field (Schrödinger Release 2022-1).[12] Dihedral angle analysis of compounds from the Cambridge Structural Database (CSD)[13] was performed with ConQuest and Mogul [14,15,16] to validate the geometry of the docking poses of **3e**.

ACKR3 in complex with the docking poses of *trans*-**3e** and *cis*-**3e** were subjected to 100 ns molecular dynamics simulations using Desmond (Schrödinger release 2022-1).[1,17] Both intracellular and extracellular Fabs bound to opposite sides of the receptor were discarded. WaterMap was used to hydrate the receptor, and water molecules fitting into receptor cavities were retained. The system was then embedded in an equilibrated POPC (1-palmitoyl-2-oleoyl-sn-glycero-3-phosphocholine) bilayer and parameterised using the OPLS4 force field using the System Builder in Maestro.[18] Desmond default relaxation membrane model system protocol was carried out before the simulation. 100 ns simulations were performed at 300K/1atm in the NPT ensemble using a Nose-Hoover thermostat and a Martyna–Tobias–Klein barostat with a 2.0 ps relaxation time. Coulomb interactions were evaluated using a 9 Å short-range cut. The resulting molecular dynamics trajectories were analysed with the simulation interactions diagram method in Maestro.

Conformers of each derivative were generated in MacroModel (Schrödinger Release 2022-1: MacroModel, Schrödinger, LLC, New York, NY, 2021) using the OPLS4 force field, GB/SA water. Molecular energy minimizations were performed using the PRCG method with 5000 maximum iterations and 0.001 gradient convergence threshold. The conformational searches were carried out by application of the MCMM torsional sampling method, performing automatic setup with 20 kJ/mol in the energy window for saving structure and a 0.5 Å cutoff distance for redundant conformers. Minimum energy conformations of *trans*-**3e** and *cis*-**3e** were superposed to minimum energy conformation of **2** to generate Fig. 1C.

### Pharmacology

#### Materials

HBSS (with $Ca^{2+}$ and $Mg^{2+}$), Dulbecco's Modified Eagle's Medium (DMEM; high glucose), 0.05% trypsin solution and penicillin/streptomycin solution were purchased from Thermo Scientific (Waltham, United States). Fetal bovine serum (FBS) was obtained from Bodinco (Alkmaar, the Netherlands). Linear 25 kDa polyethylenimine (PEI) was purchased from Polysciences (Warrington, United States). Bovine serum albumin (BSA) was obtained from Melford (Ipswich, United Kingdom). NanoLuc luciferase substrate, furimazine (N1130) was purchased from Promega (Madison, United States). 96-well white plates were obtained from Greiner Bio-one (Kremsmünster, Austria). White low volume 384 well plates were bought from Corning (Corning, United States). Human recombinant CXCL12 (#CN-11) and fluorescently labelled CXCL12-A647 (#CAF-11) were purchased from Almac (Craigavon, United Kingdom).

#### Constructs

The HA-tagged ACKR3 was C-terminally fused to SmBit by a TSSGSSGGGGSGGGGSS-linker and subcloned in the expression plasmid pcDEF3, as previously described [19] NanoLuc-ACKR3 and LgBit-b-arrestin2 constructs were kindly provided by Dr. Hill (Nottingham University, Nottingham, United Kingdom) and Dr. Seong (Korea University, Seoul, Republic of Korea), respectively. [20,21]pNBe vectors encoding a human chemokine receptors- SmBiT and LgBiT-human β-arrestin2 and SmBit-human β- arrestin2 and CXCR4-LgBit were reported previously. [22]

#### Cell culture

Human embryonic kidney 293T cells were cultured in DMEM supplemented with 10% FBS and penicillin (100 mg/ml) and streptomycin (50 mg/ml) at 37 °C with 5% $CO_2$.

#### Nluc-ACKR3 HEK293T membrane preparation

HEK293T cells ($2{\cdot}10^6$) were seeded per 10 $cm^2$ culture dish. After 24 hours, cells were transfected with 0.25 µg plasmid DNA encoding human NanoLuc-ACKR3 and 4.75 µg pcDEF3 plasmid DNA using 30 µg linear PEI. After two days, cells were collected in PBS and centrifuged at 2700 rpm at 4°C. Next, cell pellet was resuspended in ice-cold membrane buffer (15 mM Tris, 0.3 mM EDTA, 2 mM $MgCl_2$, pH 7.4 at 4°C) and homogenized by plunging the pestle using 10 strokes at 1100 rpm. Cell homogenates were exposed to two freeze- and thaw cycles using liquid nitrogen and were centrifuged at 25.000 rpm. Pellet was resuspended in Tris-sucrose buffer (20 mM Tris, 250 mM Sucrose, pH 7.4 at 4°C). Finally, cell membranes were homogenized using a 23-gauge needle followed by snap-freezing with liquid nitrogen and then stored at -80 °C.

#### NanoBRET binding

Measurement of NanoBRET was performed in triplicate on white low-volume 384-well plates. Reactions were started by combining 0.3 nM fluorescently labelled CXCL12 (CXCL12-A647 purchased from Almac, Craigavon, United Kingdom), increasing concentrations of photoswitchable compounds ($10^{-5}$ M – $10^{-10}$ M) with 30 ng (protein) HEK293T membranes expressing NanoLuc-ACKR3. Dilutions of all the above were prepared in HBSS supplemented with 0.2% BSA. The plate was pulse-centrifuged at 2000 rpm, after addition of all components of binding reaction. After 1 hour incubation at room temperature, NanoGlo substrate was added (310-fold dilution from stock) to a final volume of 13.5 µL. The total light intensity was measured for 460 nm wavelength with 80 nm bandwidth and separately for wavelengths ≥610 nm using the PHERAstar-FSX (BMG Labtech, Ortenberg, Germany) with a dual emission filter. The ratio of light intensities (>610 nm over 460 nm) is a measure for the relative binding of CXCL12-A647 to the NanoLuc-ACKR3.

#### β-arrestin2 recruitment (NanoBit) for screening photoswitchable compounds

A suspension of HEK293T cells ($1{\cdot}10^6$) was transfected in suspension using 2 µg DNA (0.4 µg ACKR3-SmBit, 0.6 µg human LgBit-β-arrestin2 ,1 µg pcDEF3 ) and 12 µg of PEI, and seeded ($3{\cdot}10^4$/well) in 96-well plates. After 48 hrs, the culture medium was replaced with HBSS with 0.05% BSA and cells were incubated at 37 °C for 15 mins with various concentration of photoswitchable compounds ($10^{-5}$ M – $10^{-10}$ M). Lastly, NanoGlo substrate was added (310-fold dilution from stock). After 5 min incubation at 37 °C, luminescence was measured using the PHERAstar-FSX.

**β-arrestin2 recruitment (NanoBit) for testing specificity of 3e**

HEK293T cells ($1{,}5{\cdot}10^6$/well) were plated in a 6-well dish and after 24 hours co-transfected with pNBe vectors encoding a human chemokine receptor, C-terminally tagged to SmBit (150 ng) and human β-arrestin2, N- terminally fused to LgBit (15 ng). Except for CXCR4, β-arrestin2 was N-terminally fused to the SmBit and CXCR4 was C-terminally fused to the LgBit. After 24 hrs transfection, cells were harvested and incubated 20 min at 37 °C with coelenterazine -h substrate diluted 500-fold and distributed into white 96-well plates (1,5 $\cdot 10^5$ cells/well). Then, cells were stimulated with 1 µM of **3e** (*trans* or $PSS_{cis}$). As a positive control 200 nM of chemokine for each chemokine receptors was used (details in Meyrath et al., 2020 [22]). Ligand-induced, β-arrestin2 recruitment to chemokine receptors was monitored with a GloMax plate reader for 20 min.

To assess antagonist properties of compound **3e** (*trans* or $PSS_{cis}$), chemokines, activating each chemokine receptors (20 nM, except for CCR1, CCR9, CCR10 100 nM was used and for CXCR3B 50 nM was used) were added after 20 min incubation with **3e** (*trans* or $PSS_{cis}$). Signal obtained from wells treated with chemokine only was defined as 100% and signals obtained from wells without presence of agonist was defined as 0% activity.

**BRET-based localization assay**

HEK293T cells ($5{\cdot}10^6$) were seeded per 10 $cm^2$ culture dish and 24 hours later transfected with 0,4 µg ACKR3-Nluc and 3,6 µg Lyn-NeonGreen using PEI method. Next day, cells were harvested and distributed into black 96-well plates ($1{,}5{\cdot}10^6$ cells per well). Thereafter, cells were stimulated 30 min at 37°C with various concentrations of compound **3e** (*trans* or $PSS_{cis}$) or 100 nM CXCL12 and followed by addition of coelenterazine -h substrate diluted 500-fold. The total light intensity was measured at 450 nm and separately for wavelengths ≥530 nm using the GloMax plate reader.

**Nephelometry**

Increasing concentrations ($10^{-4.0}$ M – $10^{-6.5}$ M) of compound **3e** (*trans* and $PSS_{cis}$) and template compound **2** were placed in transparent flat-bottom 96-well plate. Dilutions were prepared in HBSS buffer with 1% DMSO as a final concentration. Kaolin was used as a positive control and serial dilutions were prepared as described above. After at least 1 hour, nephelometry was measured using the NEPHELO star Plus plate reader (BMG Labtech, Germany) with the following settings: 1 cycle, measurement start time 0.1 s, measurement interval time 0.1 s, laser intensity 80%, beam focus 2.0 mm, Orbital shaking mode at 200 rpm with an additional shaking time of 10 s before each cycle.

**Data analysis**

All experiments were analyzed using Graphpad Prism 9.0. For binding experiments, $pIC_{50}$ values were obtained using one-site $pIC_{50}$ model using global fitting with shared non-specific binding values.

### Synthesis procedures

All chemicals and solvents were obtained from commercial suppliers e.g. Sigma-Aldrich, Fluorochem, and Combi-Blocks, and used without purification. Compounds **7a-c** were obtained from Combi-Blocks. Anhydrous DCM, toluene, and DMF were obtained after passing through PureSolv columns. IUPAC names were generated using ChemBioDraw 19.1 (Perkin-Elmer). Monitoring of reactions was performed using thin-layer chromatography (Merck Silicagel 60 F254) by visualization under natural light for colored compounds or under a 254-nm lamp. Flash column chromatography was executed by means of Biotage Isolera equipment using Biotage SNAP or Grace GraceResolve columns. Nuclear magnetic resonance (NMR) spectra were determined with a Bruker Avance II 500 MHz or a Bruker Avance III HD 600 MHz spectrometer. Chemical shifts are reported in parts per million (ppm) against the reference compound using the signal of the residual non-deuterated solvent ($CDCl_3$ δ = 7.26 ppm ($^1$H), δ = 77.16 ppm ($^{13}$C); DMSO-d6 δ = 2.50 ppm ($^1$H), δ = 39.52 ppm ($^{13}$C)). NMR spectra were processed using MestReNova 14.0 software. The peak multiplicities are defined as follows: s, singlet; d, doublet; t, triplet; q, quartet; dd, doublet of doublets; ddd, doublet of doublets of doublets; dt, doublet of triplets; dq, doublet of quartets; td, triplet of doublets; tt, triplet of triplets; qd, quartet of doublets; p, pentet; dp, doublet of pentets; br, broad signal; m, multiplet. For NMR listings, in addition to specific instructions that are given by the journal in the guidelines for authors the following additional procedures were used: 1) Multiplicity is not solely reported based on peak shapes, but also distinguishes the coupling to all non-equivalent protons that have similar J values; 2) If additional smaller couplings are observed but are too small for accurate quantitation because the precision is smaller than the digital resolution, a symbol $^{\Delta}$ will be used; 3) The notation 'm' is used in case of obscured accurate interpretation as a result of (i) overlapping signals for different protons, or (ii) a result of overlapping signal lines within the same proton signal; 4) For any rotamers or diastereomers, signals will be listed separately; 5) NMR signals that could only be detected with HSQC analysis are denoted with a $^{\#}$ symbol; 6) NMR signals that could only be detected with HMBC analysis are denoted with a * symbol; 7) If one or more signals remain undetected after extensive 1D and 2D NMR analyses, this will be mentioned. 8) Signals for exchangeable proton atoms (such as NH and OH groups) are only listed if clearly visible (excluding e.g. the use of $D_2O$ or $CD_3OD$) and if confirmed by a $D_2O$ shake and/or HSQC analysis. Purity determination was performed with Liquid Chromatography using a Shimadzu LC-20AD liquid chromatography pump system with a Shimadzu SPDM20A photodiode array detector and MS detection with a Shimadzu LCMS-2010EV mass spectrometer operating in both positive and negative ionization mode. A Waters XBridge C18 column 5 µm 4.6x50 mm was used at 40 ºC. The mobile phase used was a mixture of A = $H_2O$ + 0.1% HCOOH and B = acetonitrile (MeCN) + 0.1% HCOOH. The eluent program used is as follows: flow rate: 1.0 mL/min, start 95% A in a linear gradient to 10% A over 4.5 min, hold 1.5 min at 10% A, in 0.5 min in a linear gradient to 95% A, hold 1.5 min at 95% A, total runtime: 8.0 min. For measuring in **acidic mode**, the mobile phase was a mixture of A = $H_2O$ + 0.1% HCOOH and B = MeCN + 0.1% HCOOH. The eluent program used is as follows: flow rate of 1.0 mL/min, start 5% B, linear gradient to 90% B in 4.5 min, then 5.5 min at 90% B, then linear gradient to 5% B in 0.5 min, then 1.5 min at 5% B, total run time of 12 min. For measuring in **basic mode**, the mobile phase was a mixture of A = $H_2O$ + 10% buffer and B = MeCN + 10% buffer. The buffer mentioned is a 0.4% (w/v) $NH_4HCO_3$ aq. soln., adjusted to pH 8.0 with aq. $NH_4OH$. The eluent program used is as follows: flow rate of 1.0 mL/min, start 5% B, linear gradient to 90% B in 4.5 min, then 5.5 min at 90% B, then linear gradient to 5% B in 0.5 min, then 1.5 min at 5% B, total run time of 12 min. High-resolution mass spectra (HRMS) were recorded on a Bruker micrOTOF mass spectrometer using ESI in positive ion mode (HRMS).

Compound purities were calculated as the percentage peak area of the analyzed compound by UV detection at 254 nm. Unless mentioned otherwise, all compounds have a purity > 95 %. All reactions with photosensitive compounds were carried out in glassware covered with aluminum foil. All analyses of photosensitive compounds were carried out under dimmed or red light.

### Chemistry procedures

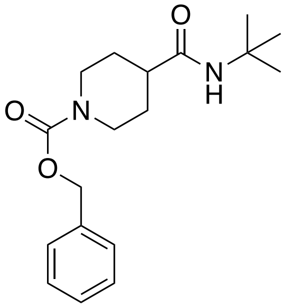

**Benzyl 4-(*tert*-butylcarbamoyl)piperidine-1-carboxylate (5).**
Carboxylic acid **5** (6.32 g, 24.0 mmol) was dissolved in DCM (60 mL) at rt. $(COCl)_2$ (4.50 mL, 52.8 mmol) and DMF (2 drops) were added. After 3 h at rt, the reaction mixture was concentrated *in vacuo*. This afforded the acid chloride (1.68 g) as a white solid, which was used without further purification. 2-Methylpropan-2-amine (308 mg, 4.22 mmol) was dissolved in DCM (4 mL) and TEA (1.60 mL, 11.5 mmol) was added. This reaction mixture was cooled to 0 °C and the acid chloride (1.08 g, 3.83 mmol) was added dropwise. The mixture was allowed to reach rt and stirred overnight. The mixture was washed with 1 M HCl and brine. The combined organic layers were dried with $MgSO_4$, filtered and concentrated *in vacuo*. The crude material was purified by flash chromatography (*c*Hex/EtOAc, 85/45 to 15/55%) to afford the product (1.06 g, 87%) as a white solid. **LC/MS (acid mode)** $R_t$ = 4.29 min, purity = 89%, $[M+H]^+$ = 319. **$^1$H NMR** (500 MHz, DMSO-d6) δ 7.40 – 7.27 (m, 5H), 5.06 (s, 2H), 4.04 – 3.96 (m, 2H), 2.87 – 2.66 (m, 2H), 2.26 (tt, *J* = 11.5, 3.8 Hz, 1H), 1.64 – 1.57 (m, 2H), 1.44 – 1.32 (m, 2H), 1.22 (s, 9H). **$^{13}$C NMR** (126 MHz, DMSO-d6) δ 174.14, 154.89, 137.49, 128.91, 128.30, 127.96, 66.57, 50.22, 43.56, 42.44, 28.99, 28.76.

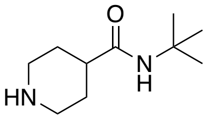

***N*-(*tert*-butyl)piperidine-4-carboxamide (6).**
CBz-protected compound **6** (7.65 g, 24.0 mmol) was dissolved in MeOH (75 mL) to which Pd/C (1.30 g, 5%) was added. The mixture was hydrogenated overnight at rt using a $H_2$ balloon. The catalyst was removed by filtration. The filtrate was concentrated *in vacuo* to afford the product (4.12 g, 93%) as a white solid. **LC/MS (acid mode)** $R_t$ = 2.51 min, purity= 99% at 200 nm, $[M+H]^+$ = 185. **$^1$H NMR** Rotamers were observed in an approximate ratio of 1:4 (500 MHz, DMSO-d6) δ Rotamer 1: δ 3.20 (s, 1H), 2.48 – 2.40 (m, 2H), 2.03 – 1.93 (m, 2H), 1.68 – 1.50 (m, 1H), 1.03 – 0.96 (m, 2H), 0.97 – 0.85 (m, 2H), 0.50 (s, 9H). Rotamer 2: 3.20 (s, 1H), 2.18 – 2.09 (m, 2H), 1.73 – 1.63 (m, 2H), 1.35 – 1.25 (m, 1H), 1.03 – 0.96 (m, 2H), 0.97 – 0.85 (m, 2H), 0.51 (s, 9H). **$^{13}$C NMR** (126 MHz, $CDCl_3$) Rotamer 1 : δ 174.80#, 40.99, 35.87, 33.55, 18.40, 18.98. Rotamer 2 : δ 174.80#, 40.99, 35.87, 35.05, 20.47, 19.46.

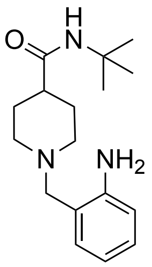

**1-(2-aminobenzyl)-*N*-(*tert*-butyl)piperidine-4-carboxamide (8a).**
Amine **7** (1.05 g, 5.69 mmol) was dissolved in DCM (25 mL). Aldehyde **7a** (1.7 g, 7.68 mmol) was added followed by $NaBH(OAc)_3$ (1.93 g, 9.11 mmol). The mixture was stirred overnight at rt. The mixture was quenched with 1 M aq. NaOH. The organic layer was separated and the aqueous layer was extracted twice with DCM. The combined organic layers were dried over $Na_2SO_4$, filtered, and concentrated *in vacuo*. The crude material was purified using flash chromatography (*c*Hex/EtOAc = 50/50 to 0/100). The resulting intermediate was dissolved in DCM (20 mL). TFA (5 mL) was added slowly and the mixture was stirred overnight at rt. The crude mixture was co-evaporated with MeOH three times. The resulting powder was dissolved in DCM and washed with 2 M aq. NaOH. The combined organic layers were dried over $Na_2SO_4$, filtered, and concentrated *in vacuo*. This afforded the product (0.69 g, 42%) as a white solid. **LC/MS (acid mode)** $R_t$ = 2.83 min, purity= 98%, $[M+H]^+$ = 290. $^1$**H NMR** (500 MHz, $CDCl_3$) δ 8.14 (s, 1H), 7.06 (ddd, *J* = 7.8, 7.8, 1.5 Hz, 1H), 6.94 (dd, *J* = 7.8, 1.7 Hz, 1H), 6.66 – 6.58 (m, 2H), 5.48 (s, 1H), 3.58 (s, 2H), 3.10 – 2.90 (m, 2H), 2.21 – 2.10 (m, 2H), 3.10 – 2.90 (m, 2H), 1.87 – 1.79 (m, 2H), 1.80 – 1.68 (m, 2H), 1.29 (s, 9H).$^{13}$**C NMR** (126 MHz, $CDCl_3$) δ 174.26, 147.27, 131.16, 129.04, 119.05*, 117.62, 116.07, 61.18, 52.23, 51.09, 40.98, 28.84, 28.37.

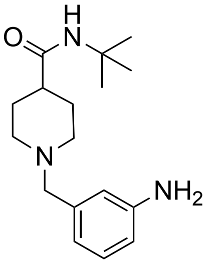

**1-(3-aminobenzyl)-*N*-(*tert*-butyl)piperidine-4-carboxamide (8b).**
Amine **7** (100 mg, 0.54 mmol) was dissolved in DCM (5 mL). Aldehyde **7b** (180 mg, 0.81 mmol) was added followed by $NaBH(OAc)_3$ (184 mg, 0.86 mmol). The mixture was stirred overnight at rt. The mixture was quenched with 1 M aq. NaOH. The organic layer was separated and the aqueous layer was extracted twice with DCM. The combined organic layers were dried over $Na_2SO_4$, filtered, and concentrated *in vacuo*. The crude material was purified using flash chromatography (*c*Hex/EtOAc = 50/50 to 0/100). The resulting intermediate was dissolved in DCM (5 mL). TFA (1 mL) was added slowly and the mixture was stirred overnight at rt. The crude mixture was co-evaporated with MeOH three times. The resulting powder was dissolved in DCM and washed with 2 M aq. NaOH. The combined organic layers were dried over $Na_2SO_4$, filtered, and concentrated *in vacuo*. This afforded the product (101 mg, 64%) as a white solid. **LC/MS (acid mode)** $R_t$ = 2.27 min, purity= 97%, $[M+H]^+$ = 290. $^1$**H NMR** (500 MHz, $CDCl_3$) δ 7.08 (dd, *J* = 7.9, 7.9 Hz, 1H), 6.70 – 6.66 (m, 2H), 6.57 (ddd, *J* = 7.9, 1.7, 1.5 Hz 1H), 5.24 (s, 1H), 3.63 (s, 1H), 3.39 (s, 2H), 3.00 – 2.88 (m, 2H), 2.00 – 1.90 (m, 3H), 1.81 – 1.75 (m, 2H), 1.76 –

1.65 (m, 2H), 1.33 (s, 9H). **$^{13}$C NMR** (151 MHz, $CDCl_3$) δ 174.61, 146.51, 139.83*, 129.19, 119.61, 115.80, 113.99, 63.35, 53.37, 51.06, 44.44, 29.24, 28.99.

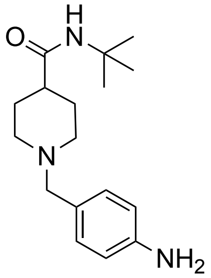


**1-(4-aminobenzyl)-*N*-(*tert*-butyl)piperidine-4-carboxamide (8c).**
Amine **7** (0.80 g, 4.34 mmol) was dissolved in DCM (20 mL). Aldehyde **7c** (1.01 g, 4.56 mmol) was added followed by $NaBH(OAc)_3$ (1.47 g, 6.94 mmol). The mixture was stirred overnight at rt. The mixture was quenched with 1 M aq. NaOH. The organic layer was separated and the aqueous layer was extracted twice with DCM. The combined organic layers were dried over $Na_2SO_4$, filtered, and concentrated *in vacuo*. The crude material was purified using flash chromatography (*c*Hex/EtOAc = 50/50 to 0/100). The resulting intermediate was dissolved in DCM (20 mL). TFA (5 mL) was added slowly and the mixture was stirred overnight at rt. The crude mixture was co-evaporated with MeOH three times. The resulting powder was dissolved in DCM and washed with 2 M aq. NaOH. The combined organic layers were dried over $Na_2SO_4$, filtered, and concentrated *in vacuo*. This afforded the product (2.07 g, 48%) as a white solid. **LC/MS (acid mode)** $R_t$ = 2.25 min, purity= 97 %, $[M+H]^+$ = 290. **$^1$H NMR** (500 MHz, $CDCl_3$) δ 7.10 ($d^{\Delta}$, *J* = 7.9 Hz, 2H), 6.64 (ddd, *J* = 7.9, 2.5, 1.8 Hz, 2H), 5.29 (s, 1H), 3.64 (s, 2H), 3.44 (s, 2H), 3.12 – 2.73 (m, 2H), 2.00 (m, 3H), 1.81 (m, 2H), 1.78 – 1.63 (m, 2H), 1.32 (s, 9H). **$^{13}$C NMR** (126 MHz, $CDCl_3$) δ 174.47, 145.69*, 130.65, 115.04, 62.63, 51.08, 49.48, 49.39*, 42.62*, 28.96, 28.87.

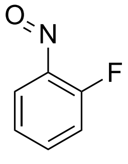


**1-fluoro-2-nitrosobenzene (10a).**
To a solution of aniline **9a** (1.00 g, 8.99 mmol) in DCM (20 mL) was added a solution of Oxone® (11.1 g, 17.9 mmol) in water (70 mL). The biphasic mixture was vigorously stirred until the consumption of the starting aniline (TLC analysis). The aqueous layer was discarded and the green organic layer was washed with 1 M aq. HCl solution, aq. satd. $NaHCO_3$, and brine. The organic layer was dried over $MgSO_4$ and filtered. This afforded a product solution that was used as such in the next step.

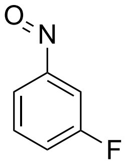


**1-fluoro-3-nitrosobenzene (10b).**
To a solution of aniline **9b** (0.80 g, 7.19 mmol) in DCM (16 mL) was added a solution of Oxone® (5.85 g, 13.4 mmol) in water (55 mL). The biphasic mixture was vigorously stirred until the consumption of the starting aniline (TLC analysis). The aqueous layer was discarded and the green organic layer was washed with 1 M aq. HCl solution, aq. satd. $NaHCO_3$, and brine. The

organic layer was dried over $MgSO_4$ and filtered. This afforded a product solution that was used as such in the next step.

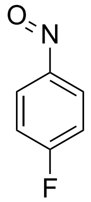

**1-fluoro-3-nitrosobenzene (10c).**

To a solution of aniline **9c** (0.50 g, 4.50 mmol) in DCM (10 mL) was added a solution of Oxone® (5.53 g, 8.99 mmol) in water (35 mL). The biphasic mixture was vigorously stirred until the consumption of the starting aniline (TLC analysis). The aqueous layer was discarded and the green organic layer was washed with 1 M aq. HCl solution, aq. satd. $NaHCO_3$, and brine. The organic layer was dried over $MgSO_4$ and filtered. This afforded a product solution that was used as such in the next step.

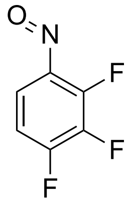

**1,2,3-difluoro-4-nitrosobenzene (10d).**

To a solution of aniline **9d** (0.50 g, 3.39 mmol) in DCM (8 mL) was added a solution of Oxone® (3.00 g, 4.88 mmol) in water (20 mL). The biphasic mixture was vigorously stirred until the consumption of the starting aniline (TLC analysis). The aqueous layer was discarded and the green organic layer was washed with 1 M aq. HCl solution, aq. satd. $NaHCO_3$, and brine. The organic layer was dried over $MgSO_4$ and filtered. This afforded a product solution that was used as such in the next step.

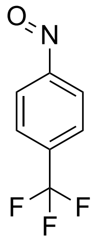

**1-nitroso-4-(trifluoromethyl)benzene (10e).**

To a solution of aniline **9e** (0.50 g, 3.10 mmol) in DCM (8 mL) was added a solution of Oxone® (3.81 g, 6.20 mmol) in water (25 mL). The biphasic mixture was vigorously stirred until the consumption of the starting aniline (TLC analysis). The aqueous layer was discarded and the green organic layer was washed with 1 M aq. HCl solution, aq. satd. $NaHCO_3$, and brine. The organic layer was dried over $MgSO_4$ and filtered. This afforded a product solution that was used as such in the next step.

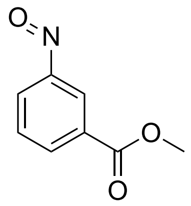

**Methyl 3-nitrosobenzoate (12).**

To a solution of aniline **11** (2.00 g, 13.2 mmol) in DCM (30 mL) was added a solution of Oxone® (8.13 g, 13.2 mmol) in water (70 mL). The biphasic mixture was vigorously stirred until the consumption of the starting aniline (TLC analysis). The aqueous layer was discarded and the green organic layer was washed with 1 M aq. HCl solution, aq. satd. $NaHCO_3$, and brine, then dried over $MgSO_4$. This afforded a product solution used as such in the next step.

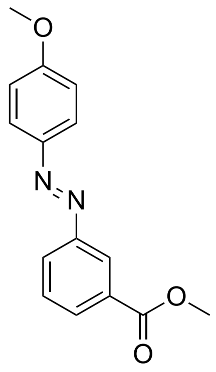

**Methyl (*E*)-3-((4-methoxyphenyl)diazenyl)benzoate (13a).**

Nitroso compound **12** (174 mg, 1.05 mmol) and 4-methoxyaniline **9f** (100 mg, 0.81 mmol) were dissolved in DCM/AcOH (1:1). The mixture was stirred for 16 h at rt. The solvents were removed *in vacuo* after which EtOAc was added. The organic phase was washed with aq. satd. $NaHCO_3$ and brine, dried over $Na_2SO_4$, filtered, and concentrated *in vacuo.* The resulting residue was purified with flash chromatography (*c*Hex/EtOAc, 80/20 to 20/80%) to afford the product (150 mg, 68%) as an orange solid. **LC/MS (acid mode)** $R_t$ = 5.37 min, purity= 96%, $[M+H]^+$ = 271, $\lambda_{max}$= 347 nm. **$^1$H NMR** (600 MHz, $CDCl_3$) 8.53 (dd, *J* = 2.0, 2.0 Hz, 1H), 8.11 (ddd, *J* = 7.6, 1.8, 0.8 Hz, 1H), 8.06 (ddd, *J* = 7.9, 2.0, 1.1 Hz, 1H), 7.95 (ddd, *J* = 8.8, 3.0, 1.9 Hz, 2H), 7.58 (dd, *J* = 7.8, 7.8 Hz, 1H), 7.03 (ddd, *J* = 8.9, 3.1, 2.0 Hz, 2H), 3.97 (s, 3H), 3.90 (s, 3H). **$^{13}$C NMR** (151 MHz, $CDCl_3$) δ 166.83, 162.56, 152.89, 147.01, 131.37, 131.19, 129.26, 126.88, 125.15, 123.85, 114.43, 55.76, 52.46.

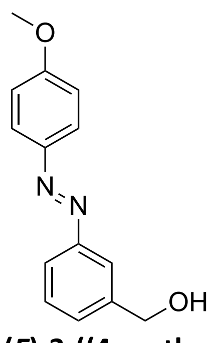

**(*E*)-3-((4-methoxyphenyl)diazenyl)**benzaldehyde **(14a).**

Ester **13a** (150 mg, 0.55 mmol) was dissolved in anhydrous *c*Hex. DIBAL-H (1.0 M in hexanes, 2.20 mL, 2.20 mmol) was added slowly at 0-5°C. The reaction mixture was warmed slowly to rt. After 4 h of stirring, the solution was quenched with satd. aq. $NH_4Cl$ (15 mL). Aq. Rochelle salt solution (10%) (8 mL) and EtOAc (10 mL) were added and the resulting mixture was stirred at rt for 1 h. The layers were separated and the organic layer was washed with brine, dried over $MgSO_4$, and filtered. The filtrate was concentrated *in vacuo* and purified by flash chromatography (*c*Hex/EtOAc, 85/45 to 15/55) to afford the product (70 mg, 52%) as an orange solid. **LC/MS (acid mode)** $R_t$ = 4.23 min, purity= 99%, $[M+H]^+$ = 243, $\lambda_{max}$ = 348 nm. **$^1$H NMR** (600 MHz, $CDCl_3$) 7.93 (ddd, *J* = 9.0, 3.1, 2.0 Hz, 2H), 7.88 (dd, *J* = 2.1 ,2.1 Hz, 1H), 7.82 (ddd, *J* = 7.8, 1.7, 1.7 Hz, 1H), 7.50 (dd, *J* = 7.7, 7.6 Hz, 1H), 7.46 (ddd, *J* = 7.6, 2.1, 1.4 Hz, 1H), 7.02 (ddd, *J* = 9.0, 3.0, 2.1 Hz, 2H), 4.80 (s, 2H), 3.90 (s, 3H). **$^{13}$C NMR** (151 MHz, $CDCl_3$) δ 162.34, 153.08, 147.08, 142.12, 129.42, 129.42, 124.90, 122.55, 120.46, 114.40, 65.20, 55.75.

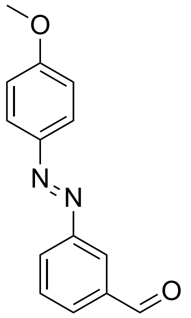

**(*E*)-(3-((4-methoxyphenyl)diazenyl)phenyl)methanol (15a).**

Alcohol **14a** (72.0 mg 0.29 mmol) and Dess-Martin periodinane (252 mg, 0.59 mmol) were dissolved in DCM (5 mL). The resulting red solution was stirred for 2 h at rt. Satd. aq. $NaHCO_3$ (8 mL) and DCM (15 mL) were added and the layers were separated. The organic layer was washed with satd. aq. $NaHCO_3$ (2x), water, and brine. The organic layer was dried over $MgSO_4$, filtered, and concentrated *in vacuo*. The residue was purified by flash chromatography (*c*Hex/EtOAc) to give the product (57 mg, 80%) as an orange solid. **LC/MS (acid mode)** $R_t$ = 5.05 min, purity= 99%, $[M+H]^+$ = 241, $\lambda_{max}$ = 348 nm. **$^1$H NMR** (600 MHz, $CDCl_3$) δ 10.13 (s, 1H), 8.36 (dd, *J* = 1.8, 1.8 Hz, 1H), 8.14 (ddd, *J* = 7.8, 2.1, 1.1 Hz, 1H), 8.01 – 7.91 (m, 3H), 7.67 (dd, *J* = 7.8, 7.8 Hz, 1H), 7.04 (ddd, *J* = 8.8, 3.1, 2.1 Hz, 2H), 3.91 (s, 3H). **$^{13}$C NMR** (151 MHz, $CDCl_3$) δ 192.04, 162.76, 153.31, 146.94, 137.46, 130.63, 129.93, 128.73, 125.27, 123.73, 114.49, 55.78.

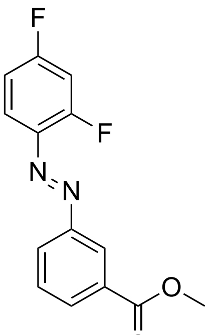

**Methyl (*E*)-3-((2,4-difluorophenyl)diazenyl)benzoate (13b).**

Nitroso compound **12** (332 mg, 2.01 mmol) and 2,4-difluoroaniline **9g** (200 mg, 1.55 mmol) were dissolved in 12 mL of DCM/AcOH (1:1). The mixture was stirred for 16 h at rt. The solvents were removed *in vacuo* after which EtOAc was added. The organic phase was washed with aq. satd. $NaHCO_3$ and brine, dried over $Na_2SO_4$, filtered, and concentrated *in vacuo.* The resulting residue was purified with flash chromatography (*c*Hex/EtOAc, 80/20 to 20/80%) to afford the product (312 mg, 73%) as an orange solid. **LC/MS (acid mode)** $R_t$ = 5.41 min, purity = 99%, $[M+H]^+$ = 277, $\lambda_{max}$ = 323 nm. **$^1$H NMR** (600 MHz, $CDCl_3$) δ 8.57 (dd, *J* = 1.9, 1.9 Hz, 1H), 8.17 (ddd, *J* = 7.7, 7.7, 1.4 Hz, 1H), 8.10 (ddd, *J* = 7.9, 2.0, 1.2 Hz, 1H), 7.84 (ddd, *J*

= 8.7, 8.7, 6.3 Hz, 1H), 7.60 (dd, *J* = 7.8, 7.8 Hz, 1H), 7.03 (ddd, *J* = 10.4, 8.6, 2.7 Hz, 1H), 6.97 (dddd, *J* = 9.0, 7.7, 2.7, 1.4 Hz, 1H), 3.97 (s, 3H). **$^{13}$C NMR** (151 MHz, $CDCl_3$) δ 166.78, 165.25 (dd, *J* = 255.2, 11.5 Hz), 161.14 (dd, *J* = 261.6, 12.4 Hz), 152.93, 137.75 (dd, *J* = 6.8, 4.0 Hz), 132.49, 131.78, 129.62, 126.90, 125.18, 119.47 (dd, *J* = 10.0, 1.5 Hz), 112.28 (dd, *J* = 22.7, 3.8 Hz), 105.58 (dd, *J* = 26.0, 23.4 Hz), 52.75.

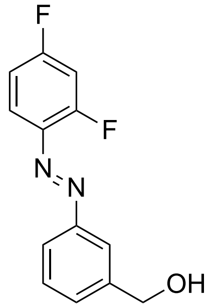


**(*E*)-(3-((2,4-difluorophenyl)diazenyl)phenyl)methanol (14b).**
Ester **13b** (158 mg, 0.57 mmol) was dissolved in anhydrous *c*Hex. DIBAL-H (1.0 M in hexanes, 2.20 mL, 2.20 mmol) was added slowly at 0-5°C. The reaction mixture was warmed slowly to rt. After 4 h stirring, the solution was quenched with satd. aq. $NH_4Cl$ (15mL). Aq. Rochelle salt solution (10%) (8 mL) and EtOAc (10 mL) were added and the resulting mixture was stirred at rt for 1h. The layers were separated and the organic layer was washed with brine, dried over $MgSO_4$, and filtered. The filtrate was concentrated *in vacuo* and purified by flash chromatography (*c*Hex/EtOAc, 85/45 to 15/55) to afford the product (70 mg, 88%) as an orange solid. **LC/MS (acid mode)** $R_t$ = 4.54 min, purity= 93%, $[M+H]^+$ = 249, $\lambda_{max}$ = 325 nm. **$^1$H NMR** (600 MHz, $CDCl_3$) δ 7.93 – 7.90 (m, 1H), 7.89 – 7.84 (m, 1H), 7.82 (ddd, *J* = 8.7, 8.7, 6.2 Hz, 1H), 7.55 – 7.49 (m, 2H), 7.02 (ddd, *J* = 10.8, 8.5, 2.7 Hz, 1H), 6.96 (dddd, *J* = 9.1, 7.8, 2.7, 1.4 Hz, 1H), 4.81 (s, 2H), 1.79 (s, 1H). **$^{13}$C NMR** (151 MHz, $CDCl_3$) 165.00 (dd, *J* = 254.5, 11.5 Hz), 160.97 (dd, *J* = 260.9, 12.5 Hz), 153.19, 142.49, 137.82, 130.19, 129.74, 123.23, 121.21, 119.40 (dd, *J* = 10.1, 1.6 Hz), 112.20 (dd, *J* = 22.6, 3.6 Hz), 105.50 (dd, *J* = 26.0, 23.7 Hz), 65.22.

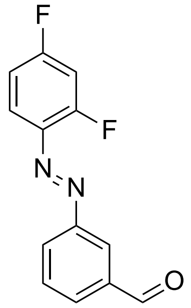


**(*E*)-3-((2,4-difluorophenyl)diazenyl)benzaldehyde (15b).**
Alcohol **14b** (100 mg, 0.40 mmol) and PCC (182 mg, 0.85 mmol) were mixed with DCM (7 mL). The resulting red solution was stirred for 2 h at rt. The reaction mixture was filtered through a short silica plug and the plug was rinsed with $Et_2O$. The filtrate was concentrated *in vacuo* and purified by flash chromatography (*c*Hex/EtOAc, 100/0 to 75/25) to afford the product (95 mg, 96%) as an orange solid. **LC/MS (acid mode)** $R_t$ = 5.09 min, purity= 99%, $[M+H]^+$ = 247, $\lambda_{max}$ = 324 nm.**$^1$H NMR** (600 MHz, $CDCl_3$) δ 10.14 (s, 1H), 8.41 (dd, *J* = 1.9, 1.9 Hz, 1H), 8.19 (ddd, *J* = 7.9, 2.0, 1.2 Hz, 1H), 8.03 (ddd, *J* = 7.6, 7.6, 1.4 Hz, 1H), 7.85 (ddd, *J* = 8.7, 8.7, 6.2 Hz, 1H), 7.70 (dd, *J* = 7.7, 7.7 Hz, 1H), 7.04 (ddd, *J* = 10.8, 8.6, 2.6 Hz, 1H), 6.98 (dddd, *J* = 9.0, 7.6, 2.7, 1.4 Hz, 1H).**$^{13}$C NMR** (151 MHz, $CDCl_3$) 191.93, 165.43 (dd, *J* = 255.6, 11.6 Hz), 161.27 (dd, *J* = 261.7, 12.5 Hz), 153.32, 137.76, 137.68 (dd, *J* = 6.8, 4.0 Hz), 131.87, 130.30, 128.97, 124.76, 119.45 (d, *J* = 10.5 Hz), 112.36 (dd, *J* = 22.7, 3.8 Hz), 105.66 (dd, *J* = 26.0, 23.6 Hz).

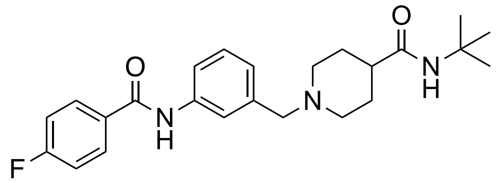

***N*-(*tert*-butyl)-1-(3-(4-fluorobenzamido)benzyl)piperidine-4-carboxamide (2)**

Aniline **8b** (60.5 mg, 0.21 mmol) was dissolved in DCM (2 mL) followed by the addition of 4-fluorobenzoyl chloride (25 μL, 0.21 mmol) and TEA (38 μL, 0.27 mmol). The reaction mixture was stirred for 1 h. Water was added to the mixture, and the organic layer was washed with 1 M NaOH. The resulting water layer was extracted with DCM. The combined organic layers were dried with $MgSO_4$, filtered and concentrated *in vacuo*. The crude material was purified by flash chromatography (EtOAc/MeOH, 100/0 to 70/30%). An extraction with aq. satd. $NaHCO_3$ /DCM afforded the product (66 mg, 77%) as a white solid. **LC/MS (acid mode)** $R_t$ = 3.43 min, purity = 98%, $[M+H]^+$ = 412. **$^1$H NMR** (600 MHz, $CDCl_3$) δ 7.92 – 7.87 (m, 2H), 7.81 (s, 1H), 7.64 (ddd, *J* = 7.7, 1.4, 1.4 Hz, 1H), 7.51 (dd, *J* = 1.9, 1.9 Hz, 1H), 7.31 (dd, *J* = 7.8, 7.8 Hz, 1H), 7.21 – 7.14 (m, 2H), 7.09 ($d^D$, *J* = 7.5 Hz, 1H), 5.24 (s, 1H), 3.49 (s, 2H), 2.96 – 2.87 (m, 2H), 2.03 – 1.93 (m, 3H), 1.82 – 1.67 (m, 4H), 1.33 (s, 9H). **$^{13}$C NMR** (151 MHz, $CDCl_3$) δ 174.61, 165.89, 164.21 (d, *J* = 260.5 Hz), 139.78, 138.04, 131.30 (d, *J* = 3.2 Hz), 129.63 (d, *J* = 9.0 Hz), 129.10, 125.44, 120.75, 119.21, 115.97 (d, *J* = 22.3 Hz), 63.01, 53.29, 51.10, 44.30, 29.15, 28.97. **HRMS** m/z (ESI+) $[M+H]^+$ calculated for $C_{24}H_{31}FN_3O_2$ 412.2395, found 412.2406.

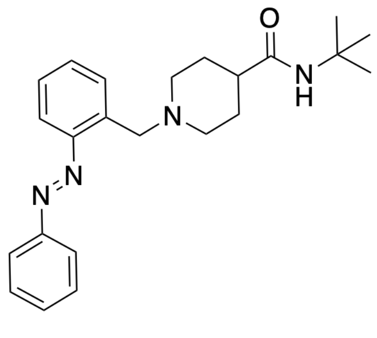

**(*E*)-*N*-(*tert*-butyl)-1-(2-(phenyldiazenyl)benzyl)piperidine-4-carboxamide (3a).**

To a stirred solution of aniline **9a** (685 mg, 2.37 mmol) in AcOH (10 mL) was added PhNO (304 mg, 2.84 mmol). The solution was heated at 90 °C for 48 h. The solvent was removed *in vacuo* after which EtOAc was added. The organic phase was washed with aq. satd. $NaHCO_3$ and brine, dried over $Na_2SO_4$, filtered and concentrated *in vacuo*. The resulting residue was purified by flash chromatography (DCM/MeOH-2%TEA, 95/5 to 85/15), affording a mixture of co-eluting **9a** and product. EtOAc was added. The solution was washed with aq. satd. $NaHCO_3$ and brine, dried over $Na_2SO_4$, filtered and concentrated *in vacuo*. This procedure was repeated two more times. The crude product was purified by flash chromatography (DCM/$NH_3$ 7N in MeOH, 95/5 to 85/15) to afford the product (35 mg, 4%) as a dark orange solid. **LC/MS (acid mode)** $R_t$ =

3.62 min, purity= 96%, $[M+H]^+$ = 290, $\lambda_{max}$ = 324 nm. **$^1$H NMR** (600 MHz, $CDCl_3$) δ 7.93 – 7.89 (m, 2H), 7.67 – 7.64 (m, 1H), 7.62 (dd, *J* = 8.1, 1.3 Hz, 1H), 7.55 – 7.50 (m, 2H), 7.49 – 7.46 (m, 1H), 7.44 (dd, *J* = 7.6, 1.4 Hz, 1H), 7.34 (dddd, *J* = 8.4, 7.4, 2.0, 1.3 Hz, 1H), 5.24 (s, 1H), 4.09 (s, 2H), 3.01 (ddd, *J* = 11.9, 3.6, 3.6 Hz, 2H), 2.11 (dd, *J* = 12.4, 10.2 Hz, 2H), 2.00 – 1.92 (m, 1H), 1.81 – 1.75 (m, 2H), 1.76 – 1.68 (m, 2H), 1.31 (s, 9H). **$^{13}$C NMR** (151 MHz, $CDCl_3$) δ 174.54, 153.02, 151.01, 137.56, 131.06, 131.01, 130.78, 129.23, 127.74, 123.12, 115.35, 57.01, 53.30, 51.01, 44.25, 29.30, 28.95. **HRMS** m/z (ESI+) $[M+H]^+$ calculated for $C_{23}H_{30}N_4O$ 379.2492, found 379.2493.

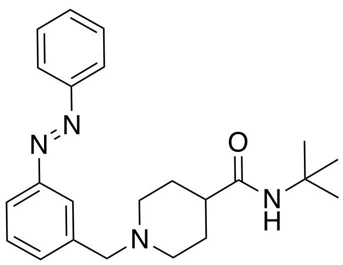


**(*E*)-*N*-(*tert*-butyl)-1-(3-(phenyldiazenyl)benzyl)piperidine-4-carboxamide (3b).**
To a stirred solution of aniline **8b** (100 mg, 0.35 mmol) in DCM (5 mL) and AcOH (5 mL) was added PhNO (37 mg, 0.35 mmol). The solution was stirred overnight at rt. The solvents were removed *in vacuo* after which DCM was added. The organic phase was washed with aq. satd. $NaHCO_3$ and brine, dried over $Na_2SO_4$, filtered, and concentrated *in vacuo.* The resulting residue was purified by flash chromatography (*c*Hex/EtOAc-2%TEA, 100/0 to 80/20) to afford the product (66 mg, 50%) as a dark orange solid. **LC/MS (basic mode)** $R_t$ = 5.68 min, purity= 99%, $[M+H]^+$ = 379, $\lambda_{max}$ = 320 nm. **$^1$H NMR** (600 MHz, $CDCl_3$) δ 7.97 – 7.89 (m, 2H), 7.89 – 7.85 (m, 1H), 7.83 ($d^{\Delta}$, *J* = 7.8 Hz, 1H), 7.54 – 7.51 (m, 2H), 7.52 – 7.50 (m, 1H), 7.50 – 7.45 (m, 2H), 5.33 (s, 1H), 3.69 (s, 2H), 3.16 – 2.96 (m, 2H), 2.19 – 2.04 (m, 2H), 1.98 – 1.85 (m, 2H), 1.84 – 1.71 (m, 3H), 1.33 (s, 9H). **$^{13}$C NMR** (151 MHz, $CDCl_3$) δ 174.48, 152.87, 152.82, 131.82, 131.12, 129.23, 129.16, 123.51, 122.98, 121.84, 62.89, 53.24, 51.10, 44.27, 29.13, 28.99. **HRMS** m/z (ESI+) $[M+H]^+$ calculated for $C_{23}H_{30}N_4O$ 379.2492, found 379.2498

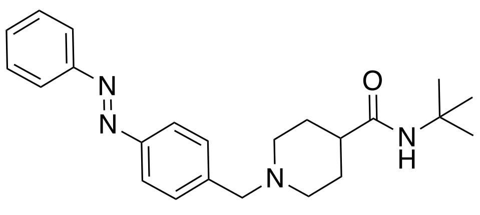


**(*E*)-*N*-(*tert*-butyl)-1-(4-(phenyldiazenyl)benzyl)piperidine-4-carboxamide (3c).**
To a stirred solution of aniline **8c** (276 mg, 1.80 mmol) in DCM (10 mL) and AcOH (10 mL) was added PhNO (193 mg, 1.80 mmol). The solution was stirred overnight at rt. The solvents were removed *in vacuo* after which DCM was added. The organic phase was washed with aq. satd. $NaHCO_3$ and brine, dried over $Na_2SO_4$, filtered, and concentrated *in vacuo.* The resulting residue was purified by flash chromatography (*c*Hex/EtOAc-2%TEA, 100/0 to 50/50 to afford the product (151 mg, 42%) as a dark orange solid. **LC/MS (basic mode)** $R_t$ = 5.67 min, purity= 96%, $[M+H]^+$ = 379, $\lambda_{max}$ = 323 nm. **$^1$H NMR** (600 MHz, $CDCl_3$) δ 7.93 – 7.89 (m, 2H), 7.87 (d, *J* = 8.3 Hz, 2H), 7.54 – 7.51 (m, 2H), 7.51 – 7.44 (m, 3H), 5.27 (s, 1H), 3.60 (s, 2H), 3.06 – 2.91 (m, 2H), 2.16 – 1.96 (m, 2H), 1.88 – 1.81 (m, 1H), 1.82 – 1.72 (m, 4H), 1.34 (s, 9H). **$^{13}$C NMR** (151 MHz, $CDCl_3$) δ 174.29, 152.72, 151.99, 141.44*, 130.92, 129.76, 129.09, 122.82, 62.63, 53.06,

51.00, 43.88#, 28.94, 28.86. **HRMS** m/z (ESI+) $[M+H]^+$ calculated for $C_{23}H_{30}N_4O$ 379.2492, found 379.2498.

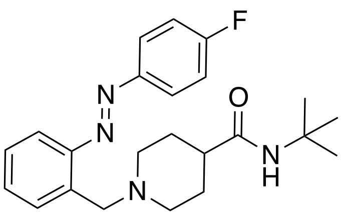

**(*E*)-*N*-(*tert*-butyl)-1-(2-((4-fluorophenyl)diazenyl)benzyl)piperidine-4-carboxamide (3d).**

To a stirred solution of aniline **8a** (85 mg, 0.30 mmol) in DCM (1 mL), AcOH (4 mL) and a solution of crude **10c** (3 mL, *vide supra*) were added. The solution was stirred for 48 h at rt. The solvents were removed *in vacuo* after which DCM was added. The organic phase was washed with aq. satd. $NaHCO_3$ and brine, dried over $Na_2SO_4$, filtered, and concentrated *in vacuo.* The resulting residue was purified by reverse flash chromatography ($H_2O$/ACN 0.1% TFA, 100/0 to 0/100). The free base was liberated from the resulting TFA salt by extraction with 1 M aq. NaOH. This afforded the product (8 mg, 7%) as a dark orange solid. **LC/MS (acid mode)** $R_t$ = 5.73 min, purity= 95 %, $[M+H]^+$ = 397, $\lambda_{max}$ =323 nm. **$^1$H NMR** (600 MHz, $CDCl_3$) δ 7.96 – 7.88 (m, 2H), 7.65 ($d^{\Delta}$ *J* = 7.8 Hz, 1H), 7.62 ($d^{\Delta}$, *J* = 8.2 Hz, 1H), 7.45 (dd, *J* = 7.4, 7.4 Hz, 1H), 7.34 (dd, *J* = 7.7, 7.7 Hz, 1H), 7.20 (dd, *J* =8.4 ,8.4 Hz, 2H), 5.24 (s, 1H), 4.08 (s, 2H), 3.07 – 2.93 (m, 2H), 2.18 – 2.03 (m, 2H), 2.01 – 1.91 (m, 1H), 1.82 – 1.76 (m, 2H), 1.76 – 1.68 (m, 2H), 1.31 (s, 9H). **$^{13}$C NMR** (151 MHz, $CDCl_3$) δ 174.48, 165.45 (d, *J* = 252.0 Hz), 150.82, 149.58 (d, *J* = 2.9 Hz), 137.39*, 131.08, 130.86, 127.81, 125.09 (d, *J* = 8.9 Hz), 116.20 (d, *J* = 22.8 Hz), 115.34, 56.99, 53.27, 51.04, 44.20, 29.83, 28.96. **HRMS** m/z (ESI+) $[M+H]^+$ calculated for $C_{23}H_{29}FN_4O$ 397.2398, found 397.2407

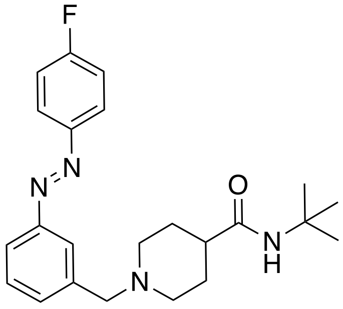

**(*E*)-*N*-(*tert*-butyl)-1-(3-((4-fluorophenyl)diazenyl)benzyl)piperidine-4-carboxamide (VUF25471, 3e).**

To a stirred solution of aniline **8b** (115 mg, 0.40 mmol) in DCM (1 mL and AcOH (3 mL) was added **10c** (2 mL, *vide supra*). The solution was stirred overnight at rt. The solvents were removed *in vacuo* after which DCM was added. The organic phase was washed with aq. satd. $NaHCO_3$ and brine, dried over $Na_2SO_4$, filtered, and concentrated *in vacuo.* The resulting residue was purified by flash chromatography (*c*Hex/EtOAc-2% TEA, 100/0 to 80/20) to afford the product (64 mg, 40%) as a dark orange solid. **LC/MS (acid mode)** $R_t$ = 3.85 min, purity= 99%, $[M+H]^+$ = 397, $\lambda_{max}$ = 324 nm. **$^1$H NMR** (600 MHz, $CDCl_3$) δ 7.94 (ddd, *J* = 5.2, 3.6, 2.0 Hz, 2H), 7.86 – 7.82 (m, 1H), 7.80 – 7.75 (m, 1H), 7.48 – 7.42 (m, 2H), 7.19 (ddd, *J* = 8.5, 8.5, 3.1 Hz, 2H), 5.27 (s, 1H), 3.59 (s, 2H), 2.99 – 2.91 (m, 2H), 2.08 – 1.95 (m, 3H), 1.83 – 1.76 (m, 2H), 1.78 – 1.70 (m, 2H), 1.33 (s, 9H). **$^{13}$C NMR** (151 MHz, $CDCl_3$) δ 174.48, 164.46 (d, *J* = 251.9 Hz), 152.66, 149.30 (d, *J* = 2.9 Hz), 139.87, 131.78, 129.11, 124.96 (d, *J* = 8.9 Hz), 123.37, 121.73, 116.14 (d, *J* = 23.0 Hz), 62.87, 53.26, 51.05, 44.21, 29.14, 28.96. **HRMS** m/z (ESI+) $[M+H]^+$ calculated for $C_{23}H_{29}FN_4O$ 397.2398, found 397.2391.

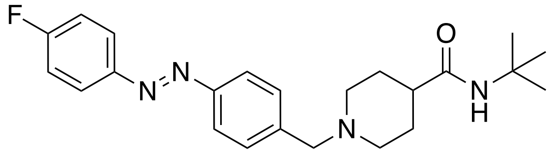

**(*E*)-*N*-(*tert*-butyl)-1-(4-((4-fluorophenyl)diazenyl)benzyl)piperidine-4-carboxamide (3f).**
To a stirred solution of aniline **8c** (110 mg, 0.38 mmol) in DCM (1 mL) and AcOH (3 mL) was added **10c** (2 mL, *vide supra*). The solution was stirred overnight at rt. The solvents were removed *in vacuo* after which DCM was added. The organic phase was washed with aq. satd. $NaHCO_3$ and brine, dried over $Na_2SO_4$, filtered, and concentrated *in vacuo.* The resulting residue was purified by flash chromatography (*c*Hex/EtOAc-2% TEA, 100/0 to 50/50) to afford the product (45 mg, 26%) as a dark orange solid. **LC/MS (basic mode)** $R_t$ = 4.4 min, purity= 98 %, $[M+H]^+$ = 397, $\lambda_{max}$ = 321 nm. **$^1$H NMR** (600 MHz, $CDCl_3$) δ 7.93 (ddd, *J* = 5.4, 3.6, 1.9 Hz, 2H), 7.85 ($d^{\Delta}$, *J* = 8.0 Hz, 2H), 7.55 – 7.43 (m, 2H), 7.19 (ddd, *J* = 8.6, 8.6, 3.3 Hz, 2H), 5.27 (s, 1H), 3.74 – 3.43 (m, 2H), 3.11 – 2.81 (m, 2H), 2.17 – 1.90 (m, 3H), 1.87 – 1.68 (m, 4H), 1.34 (s, 9H). **$^{13}$C NMR** (151 MHz, $CDCl_3$) δ 174.38, 164.47 (d, *J* = 251.6 Hz), 151.94, 149.35 (d, *J* = 3.1 Hz), 141.99*, 129.88, 124.95 (d, J =8.8 Hz), 122.92, 116.17 (d, J =23.0 Hz), 62.77, 53.27, 51.14, 43.86$^{\#}$, 29.84, 28.98. **HRMS** m/z (ESI+) $[M+H]^+$ calculated for $C_{23}H_{29}FN_4O$ 397.2398, found 397.2401.

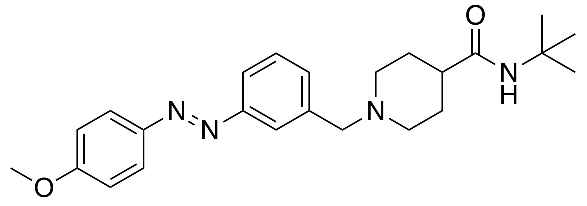

**(*E*)-*N*-(*tert*-butyl)-1-(3-((4-methoxyphenyl)diazenyl)benzyl)piperidine-4-carboxamide (3g).**
Amine **6** (55 mg, 0.30 mmol) was dissolved in DCM (3 mL). Aldehyde **15a** (79 mg, 0.33 mmol) was added followed by $NaBH(OAc)_3$ (101 g, 0.48 mmol). The mixture was stirred overnight at rt. The mixture was quenched with 1 M aq. NaOH. The organic layer was separated, and the aqueous layer was extracted twice with DCM. The combined organic layers were dried over $Na_2SO_4$, filtered, and concentrated *in vacuo*. The crude material was purified using flash chromatography (*c*Hex/EtOAc = 50/50 to 0/100) to afford the product (22 mg, 18%) as a dark orange solid. **LC/MS (basic mode)** $R_t$ = 5.53 min, purity= 96%, $[M+H]^+$ = 409, $\lambda_{max}$= 347 nm. **$^1$H NMR (**600 MHz, $CDCl_3$) δ 7.92 (dd, *J* = 9.0, 3.4 Hz, 2H), 7.84 – 7.81 (m, 1H), 7.80 – 7.77 (m, 1H), 7.53 – 7.43 (m, 2H), 7.02 (dd, *J* = 8.6, 3.1 Hz, 2H), 5.27 (s, 1H), 3.90 (s, 3H), 3.63 (s, 2H), 3.09 – 2.78 (m, 2H), 2.11 – 1.93 (m, 3H), 1.85 – 1.72 (m, 4H), 1.33 (s, 9H). **$^{13}$C NMR** (151 MHz, $CDCl_3$) δ 174.32, 162.26, 152.99, 147.16, 129.22, 124.94, 123.37, 121.60, 114.37, 62.84, 55.74, 51.16, 46.02, 43.33, 29.85, 28.97. **HRMS** m/z (ESI+) $[M+H]^+$ calculated for $C_{24}H_{32}N_4O_2$ 409.2598, found 409.2607.

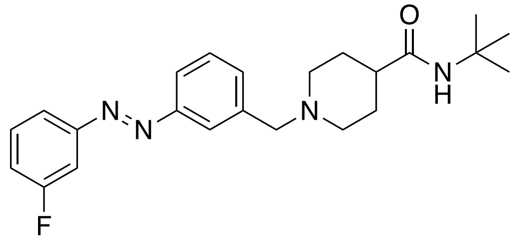

**(*E*)-*N*-(*tert*-butyl)-1-(3-((3-fluorophenyl)diazenyl)benzyl)piperidine-4-carboxamide (3h).**

To a stirred solution of aniline **8c** (100 mg, 0.35 mmol) in DCM (1 mL) and AcOH (3 mL) was added **10b** (2 mL, *vide supra*). The solution was stirred overnight at rt. The solvents were removed *in vacuo* after which DCM was added. The organic phase was washed with aq. satd. $NaHCO_3$ and brine, dried over $Na_2SO_4$, filtered, and concentrated *in vacuo.* The resulting residue was purified by flash chromatography (EtOAc-2% TEA/MeOH, 100/0 to 80/20) to afford the product (84 mg, 61%) as a dark orange solid. **LC/MS (basic mode)** $R_t$ = 5.80 min, purity= 99 %, $[M+H]^+$ = 397, $\lambda_{max}$ = 320 nm. **$^1$H NMR** (600 MHz, $CDCl_3$) δ 7.90 – 7.83 (m, 1H), 7.80 (ddd, *J* = 7.5, 1.8, 1.4 Hz, 1H), 7.75 (ddd, *J* = 7.9, 1.3, 1.3 Hz, 1H), 7.59 (ddd, *J* = 9.8, 2.2, 2.2 Hz, 1H), 7.54 – 7.40 (m, 3H), 7.16 (ddd, *J* = 8.2, 8.2, 2.6 Hz, 1H), 5.32 (s, 1H), 3.59 (s, 2H), 3.06 – 2.85 (m, 2H), 2.10 – 1.94 (m, 3H), 1.84 – 1.76 (m, 2H), 1.78 – 1.71 (m, 2H), 1.32 (s, 9H). **$^{13}$C NMR** (151 MHz, $CDCl_3$) δ 174.43, 163.35 (d, *J* = 247.5 Hz), 154.21 (d, *J* = 7.0 Hz), 152.48, 139.78*, 132.22, 130.34 (d, *J* = 8.6 Hz), 129.15, 123.56, 121.99, 120.60 (d, *J* = 2.8 Hz), 117.77 (d, *J* = 22.1 Hz), 108.10 (d, *J* = 22.7 Hz), 62.76, 53.16, 51.02, 44.04, 29.03, 28.93. **HRMS** m/z (ESI+) $[M+H]^+$ calculated for $C_{23}H_{29}FN_4O$ 397.2398, found 397.2398.

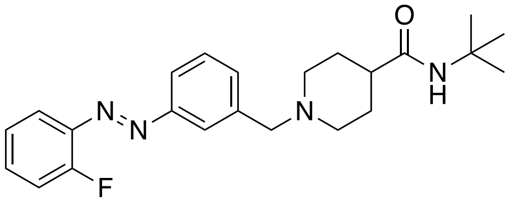

**(*E*)-*N*-(*tert*-butyl)-1-(3-((2-fluorophenyl)diazenyl)benzyl)piperidine-4-carboxamide (3i).**

To a stirred solution of aniline **8c** (100 mg, 0.35 mmol) in DCM (1 mL) and AcOH (3 mL) was added **10a** (2 mL, *vide supra*). The solution was stirred overnight at rt. The solvents were removed *in vacuo* after which DCM was added. The organic phase washed with aq. satd. $NaHCO_3$ and brine, dried over $Na_2SO_4$, filtered, and concentrated *in vacuo.* The resulting residue was purified by flash chromatography (EtOAc-2% TEA/MeOH, 100/0 to 80/20) to afford the product (98 mg, 71%) as a dark orange solid. **LC/MS (basic mode)** $R_t$ = 5.63 min, purity= 99%, $[M+H]^+$ = 397, $\lambda_{max}$ = 324 nm. **$^1$H NMR** (600 MHz, $CDCl_3$) δ 7.90 – 7.85 (m, 1H), 7.82 (dd$^{\Delta}$, *J* = 7.7, 7.7 Hz, 1H), 7.74 (ddd, *J* = 8.2, 8.2, 1.7 Hz, 1H), 7.51 – 7.47 (m, 1H), 7.48 – 7.37 (m, 2H), 7.28 – 7.22 (m, 1H), 7.21 (dd$^{\Delta}$, *J* = 7.7, 7.7 Hz, 1H), 5.33 (s, 1H), 3.60 (s, 2H), 3.22 – 2.75 (m, 2H), 2.20 – 2.02 (m, 2H), 2.02 – 1.97 (m, 1H), 1.85 – 1.76 (m, 2H), 1.78 – 1.64 (m, 2H), 1.32 (s, 9H). **$^{13}$C NMR** (151 MHz, $CDCl_3$) δ 174.44, 160.16 (d, *J* = 257.6 Hz), 152.95, 140.77 (d, *J* = 6.9 Hz), 140.1*, 132.56 (d, *J* = 8.2 Hz), 132.25, 129.15, 124.40 (d, *J* = 3.8 Hz), 123.93, 121.94, 117.89, 117.13 (d, *J* = 19.9 Hz), 62.70, 53.10, 51.02, 43.95, 28.98, 29.92. **HRMS** m/z (ESI+) $[M+H]^+$ calculated for $C_{23}H_{29}FN_4O$ 397.2398, found 397.2398.

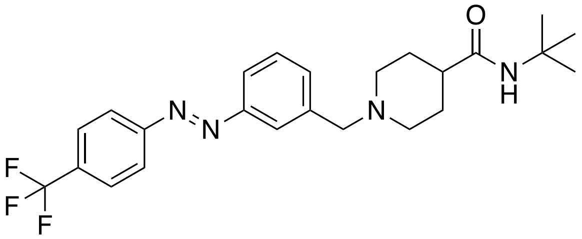

**(*E*)-*N*-(*tert*-butyl)-1-(3-((4-(trifluoromethyl)phenyl)diazenyl)benzyl)piperidine-4-carboxamide (3j).**

To a stirred solution of aniline **8c** (150 mg, 0.52 mmol) in DCM (2 mL) and AcOH (4 mL) was added **10e** (2 mL, *vide supra*). The solution was stirred overnight at rt. The solvents were removed *in vacuo* after which DCM was added. The organic phase was washed with aq. satd.

$NaHCO_3$ and brine, dried over $Na_2SO_4$, filtered, and concentrated *in vacuo.* The resulting residue was purified by flash chromatography (*c*Hex/EtOAc-2% TEA, 70/30 to 40/60) to afford the product (206 mg, 89%) as a dark orange solid. **LC/MS (basic mode)** $R_t$ = 5.85 min, purity= 98 %, $[M+H]^+$ = 447, $\lambda_{max}$ = 317 nm. **$^1$H NMR** (600 MHz, $CDCl_3$) § **$^{13}$C NMR** (151 MHz, $CDCl_3$) δ 174.42, 154.58, 152.65, 140.02*, 132.61, 132.22 k), 129.28, 126.43 (q, *J* = 3.7 Hz), 124.97 (q, *J* = 272.4 Hz), 124.97, 123.72, 122.18, 62.78, 53.30, 51.12, 44.28, 29.09, 28.98. **HRMS** m/z (ESI+) $[M+H]^+$ calculated for $C_{24}H_{29}F_3N_4O$ 447.2366, found 447.2367.
kk

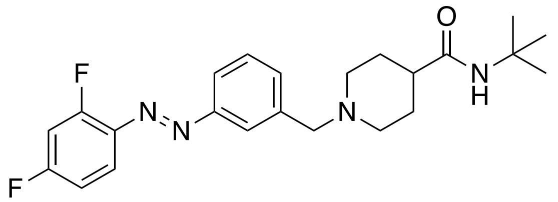

**(*E*)-*N*-(*tert*-butyl)-1-(3-((2,4-difluorophenyl)diazenyl)benzyl)piperidine-4-carboxamide (3k).**
Amine **6** (75 mg, 0.41 mmol) was dissolved in DCM (5 mL). Aldehyde **15b** (120 mg, 0.49 mmol) was added followed by $NaBH(OAc)_3$ (138 g, 0.65 mmol). The mixture was stirred overnight at rt. The mixture was quenched with 1 M aq. NaOH. The organic layer was separated, and the aqueous layer was extracted twice with DCM. The combined organic layers were dried over $Na_2SO_4$, filtered, and concentrated *in vacuo*. The crude material was purified using flash chromatography (*c*Hex/EtOAc = 50/50 to 0/100) to afford the product (15 mg, 9%) as a dark orange solid. **LC/MS (acid mode)** $R_t$ = 5.55 min, purity = 95%, $[M+H]^+$ = 415, $\lambda_{max}$ = 325 nm. **$^1$H NMR** (600 MHz, $CD_3OD$) δ 8.09 – 8.05 (m, 1H), 8.04 – 7.98 (m, 1H), ), 7.88 (ddd, *J* = 8.8, 6.3, 2.5 Hz, 1H), 7.71 – 7.64 (m, 2H), 7.55 (s, 1H), 7.24 (ddd, *J* = 11.2, 8.8, 2.7 Hz, 1H), 7.11 (dddd, *J* = 9.2, 7.9, 2.7, 1.3 Hz, 1H), 4.27 (s, 2H), 3.47 – 3.35 (m, 2H), 2.96 – 2.85 (m, 2H), 2.49 – 2.38 (m, 1H), 2.04 – 1.77 (m, 4H), 1.32 (s, 9H). **$^{13}$C NMR** (151 MHz, $CD_3OD$) δ 174.54, 166.48 (dd, *J* = 254.0, 11.8 Hz), 162.14 (dd, *J* = 260.2, 12.7 Hz), 154.32, 149.11*, 138.72 (dd, *J* = 7.0, 4.0 Hz), 135.00, 131.19, 125.81, 125.57, 120.26 (d, *J* = 10.4 Hz), 113.19 (dd, *J* = 22.9, 3.6 Hz), 106.20 (dd, *J* = 26.7, 24.0 Hz), 61.65, 53.01, 51.88, 41.87, 28.86, 27.74. **HRMS** m/z (ESI+) $[M+H]^+$ calculated for $C_{23}H_{28}F_2N_4O$ 415.2263, found 415.2320.

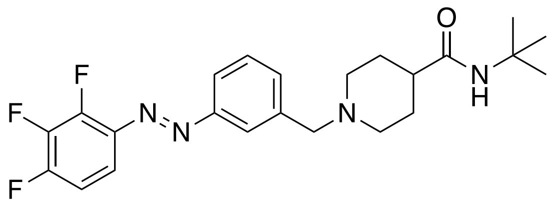

**(*E*)-*N*-(*tert*-butyl)-1-(3-((2,3,4-trifluorophenyl)diazenyl)benzyl)piperidine-4-carboxamide (3l).**
To a stirred solution of aniline **8c** (200 mg, 0.69 mmol) in DCM (3 mL) and AcOH (7 mL) was added **10d** (4 mL, *vide supra*). The solution was stirred overnight at rt. The solvents were removed *in vacuo* after which DCM was added. The organic phase was washed with aq. satd. $NaHCO_3$ and brine, dried over $Na_2SO_4$, filtered, and concentrated *in vacuo.* The resulting residue was purified by flash chromatography (*c*Hex/EtOAc-2% TEA, 100/0 to 70/30) to afford the product (60 mg, 20%) as a dark orange solid. **LC/MS (acid mode)** $R_t$ = 5.74 min, purity= 95%, $[M+H]^+$ = 433, $\lambda_{max}$ = 322 nm. **$^1$H NMR** (600 MHz, $CDCl_3$) δ 7.92 – 7.87 (m, 1H), 7.83 (dd delta, *J* = 7.8, 7.8 Hz, 1H), 7.63 – 7.53 (m, 1H), 7.54 – 7.48 (m, 1H), 7.49 – 7.43 (m, 1H), 7.05 (dddd, *J* = 9.3, 9.3, 7.1, 2.1 Hz, 1H), 5.26 (s, 1H), 3.61 (s, 2H), 3.15 – 2.92 (m, 2H), 2.10 – 1.96 (m, 3H), 1.84 – 1.73 (m, 4H), 1.33 (s, 9H). **$^{13}$C NMR** (151 MHz, $CDCl_3$) δ 174.39, 152.93 (dd, *J* = 252.5, 9.85 Hz), 152.72, 149.73 (dd, *J* = 262.8, 9.0 Hz), 142.76 (ddd, *J* = 252.8, 14.2, 14.2 Hz),

138.31 (d, *J* = 4.2 Hz), 132.76, 129.31, 124.17, 121.99, 112.15 (dd, *J* = 8.2, 4.0 Hz), 111.98 (dd, *J* = 18.6, 3.9 Hz), 62.73, 53.31, 51.13, 44.24, 28.98, 28.87. **HRMS** m/z (ESI+) $[M+H]^+$ calculated for $C_{23}H_{27}F_3N_4O$ 433.2210, found 433.2206.

### Chemical analyses

$^{1}$H NMR spectrum of **4**

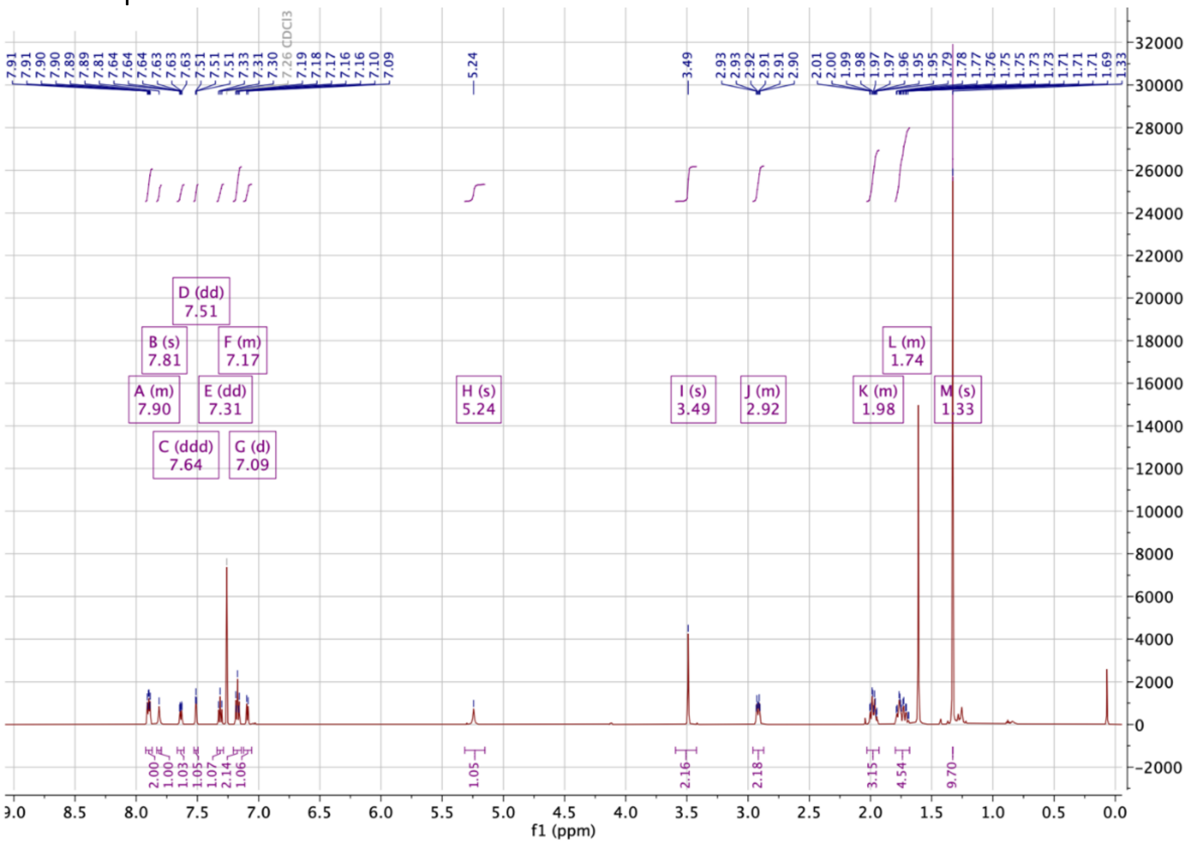


$^{13}$C NMR spectrum of **4**

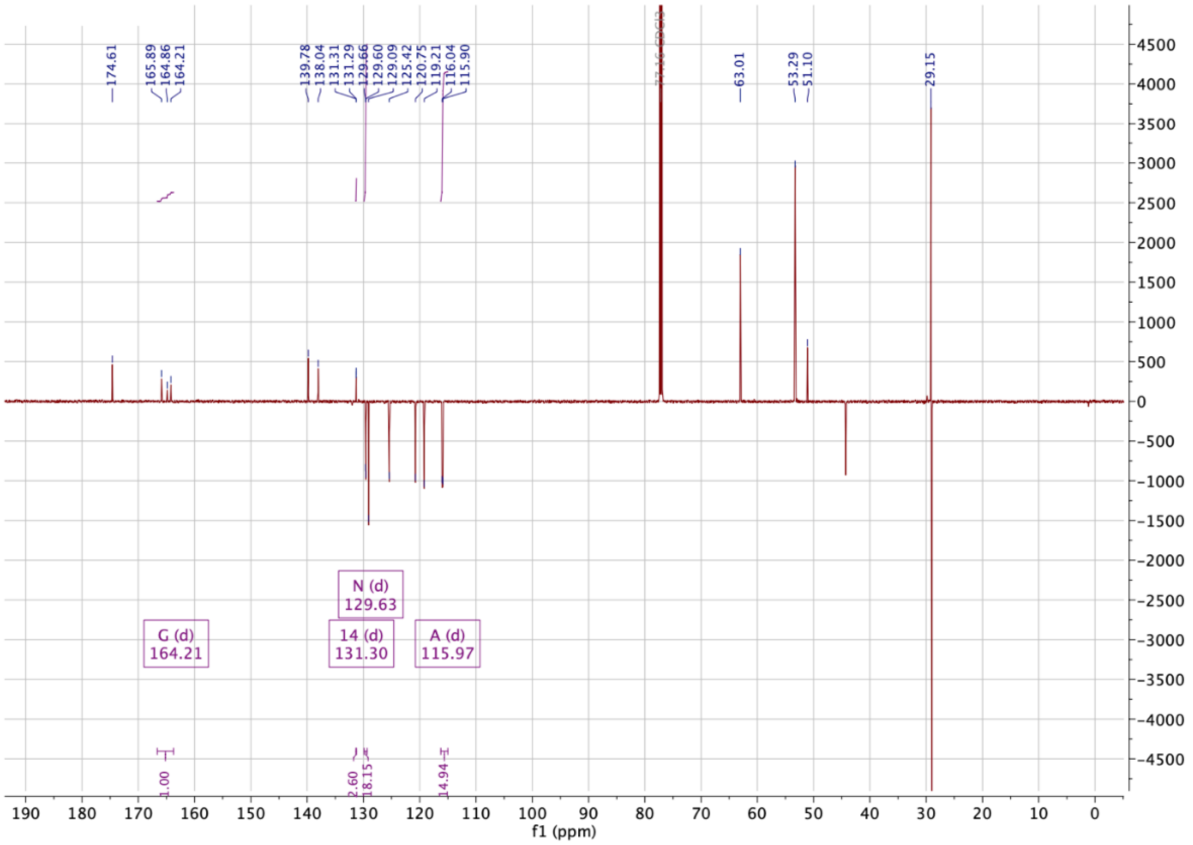

$^{1}H$ NMR spectrum of **5**

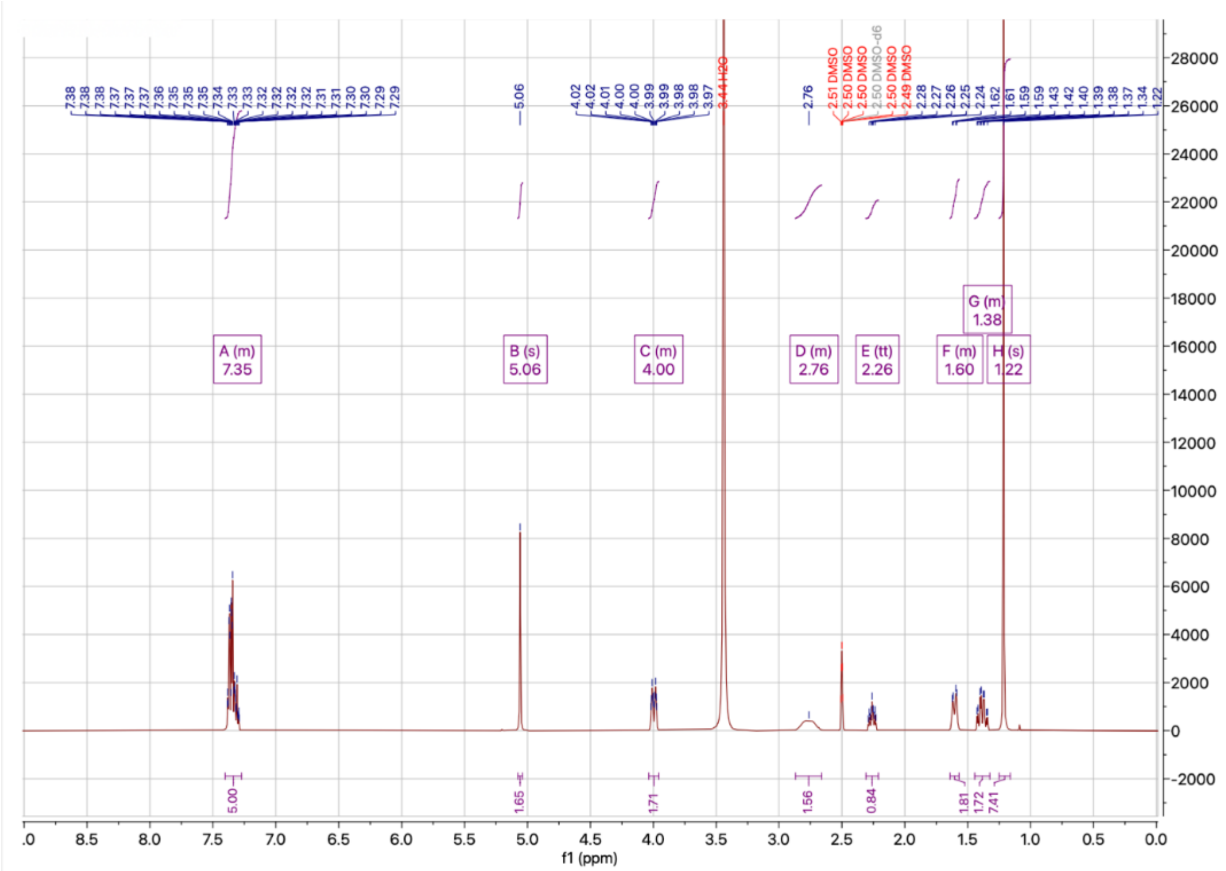


$^{13}C$ NMR spectrum of **5**

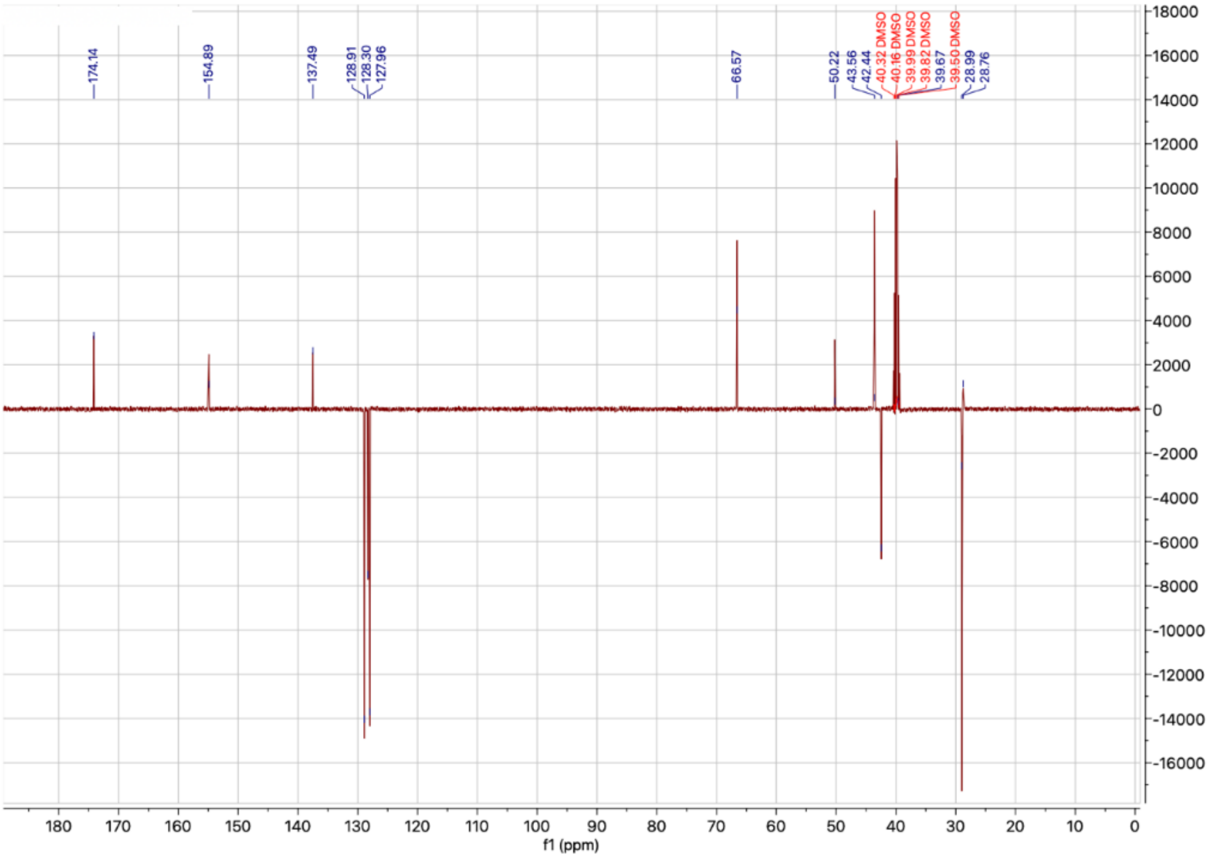

$^{1}$H NMR spectrum of **6**

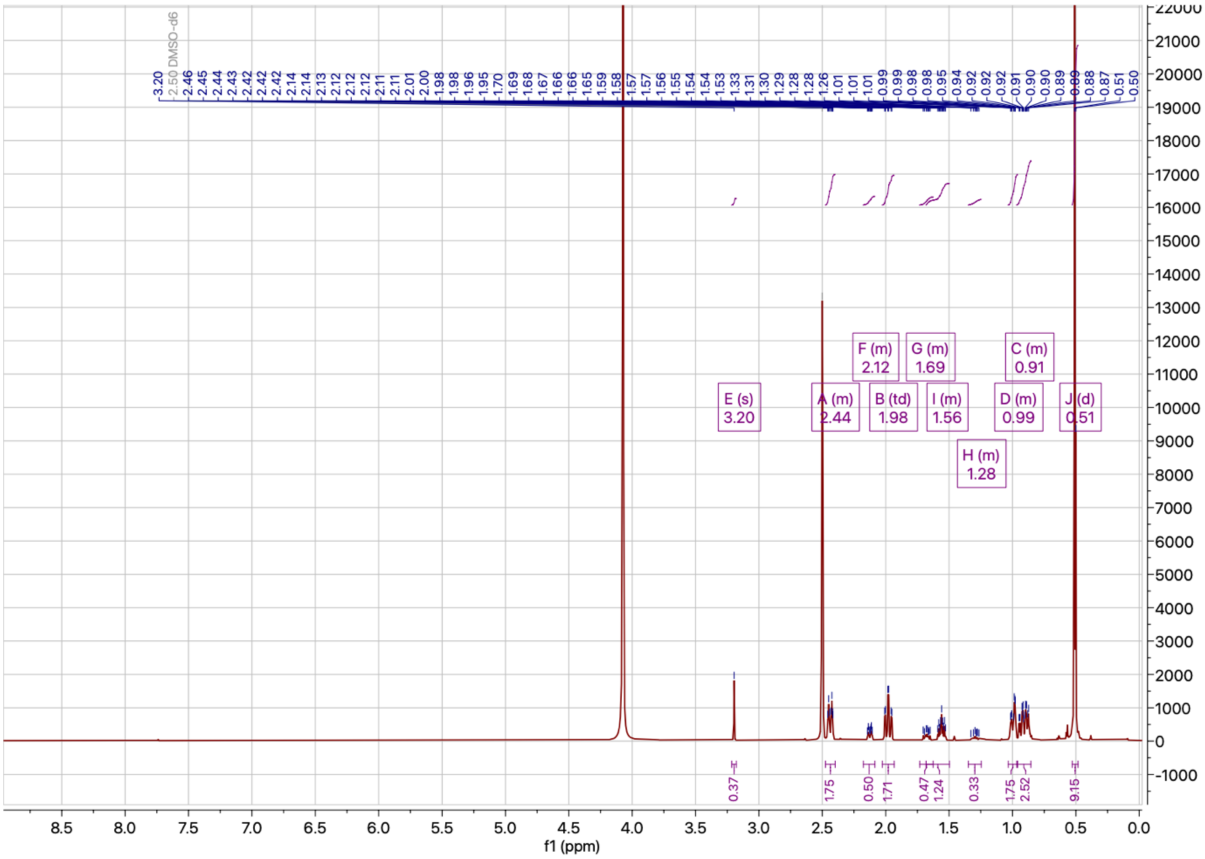

$^{13}$C NMR spectrum of **6**

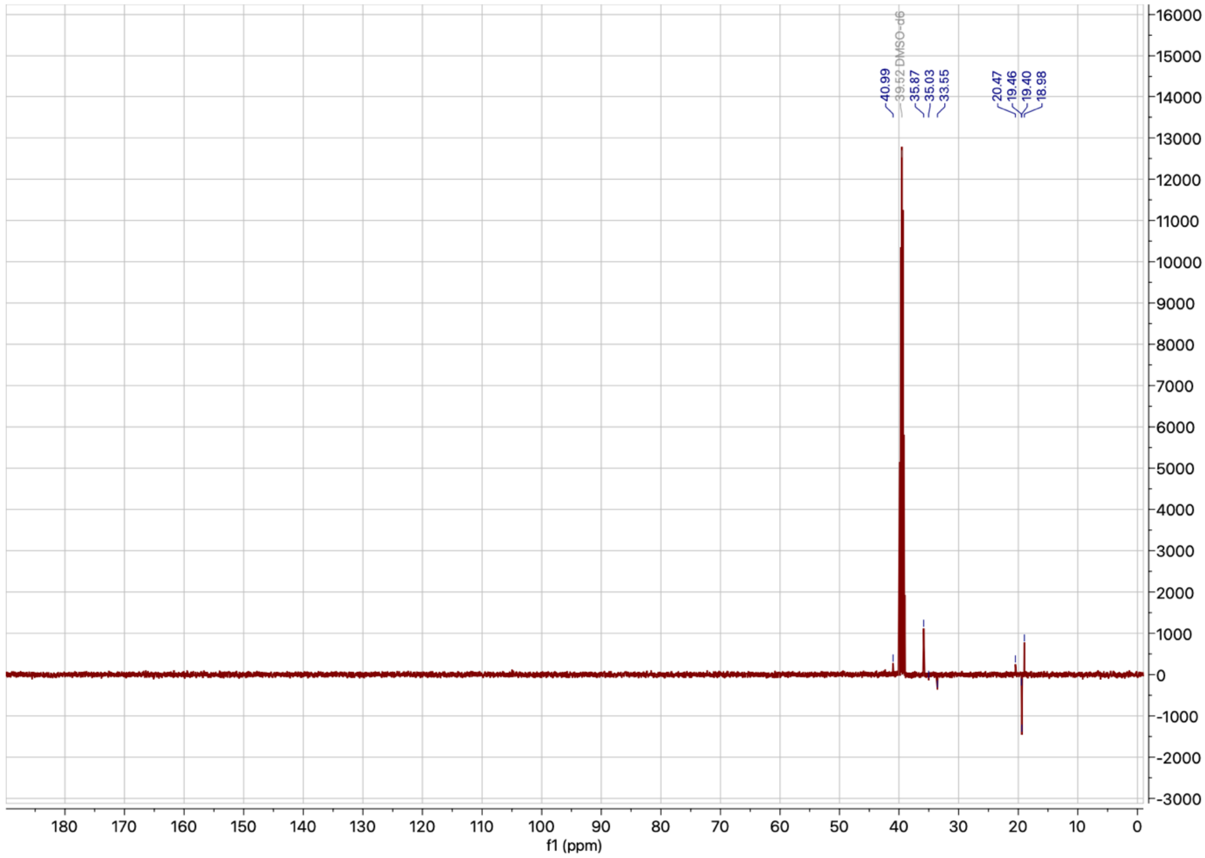

[1]H NMR spectrum of **8a**

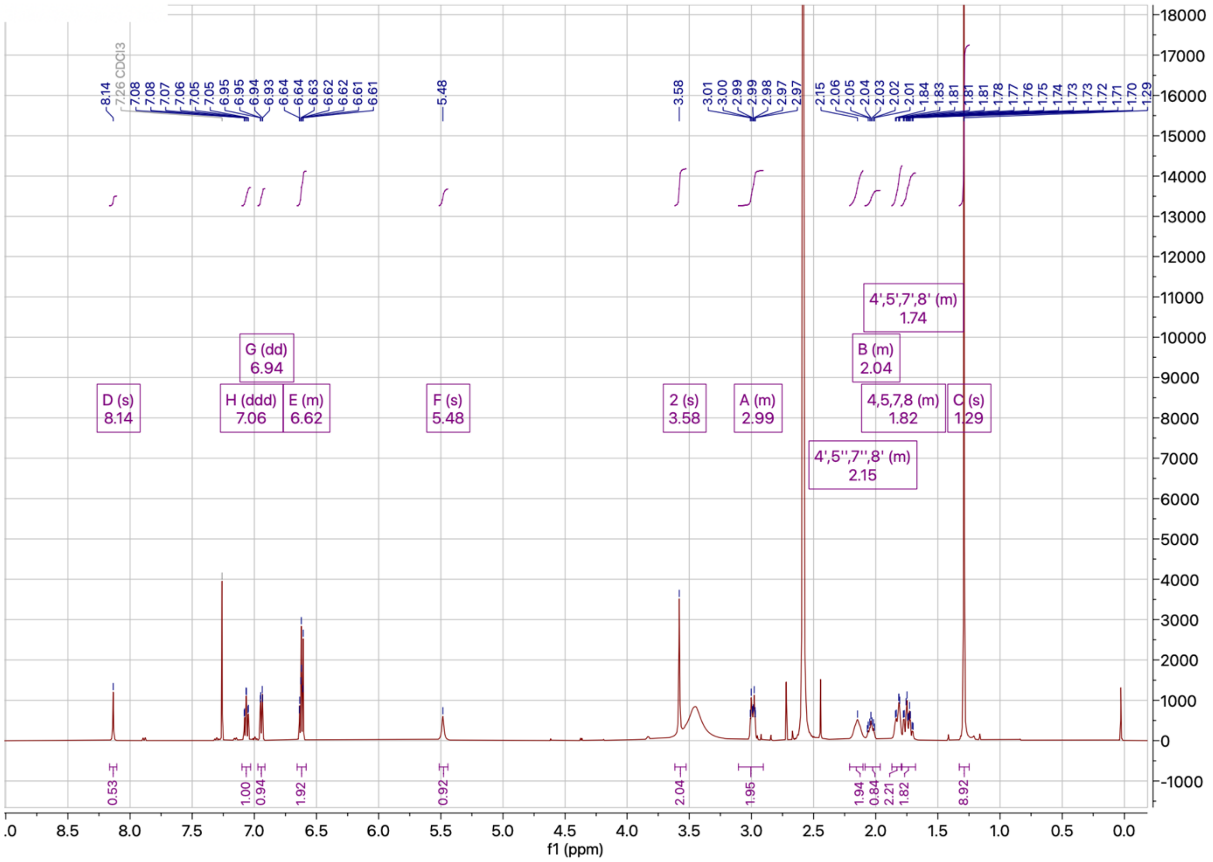


[13]C NMR spectrum of **8a**

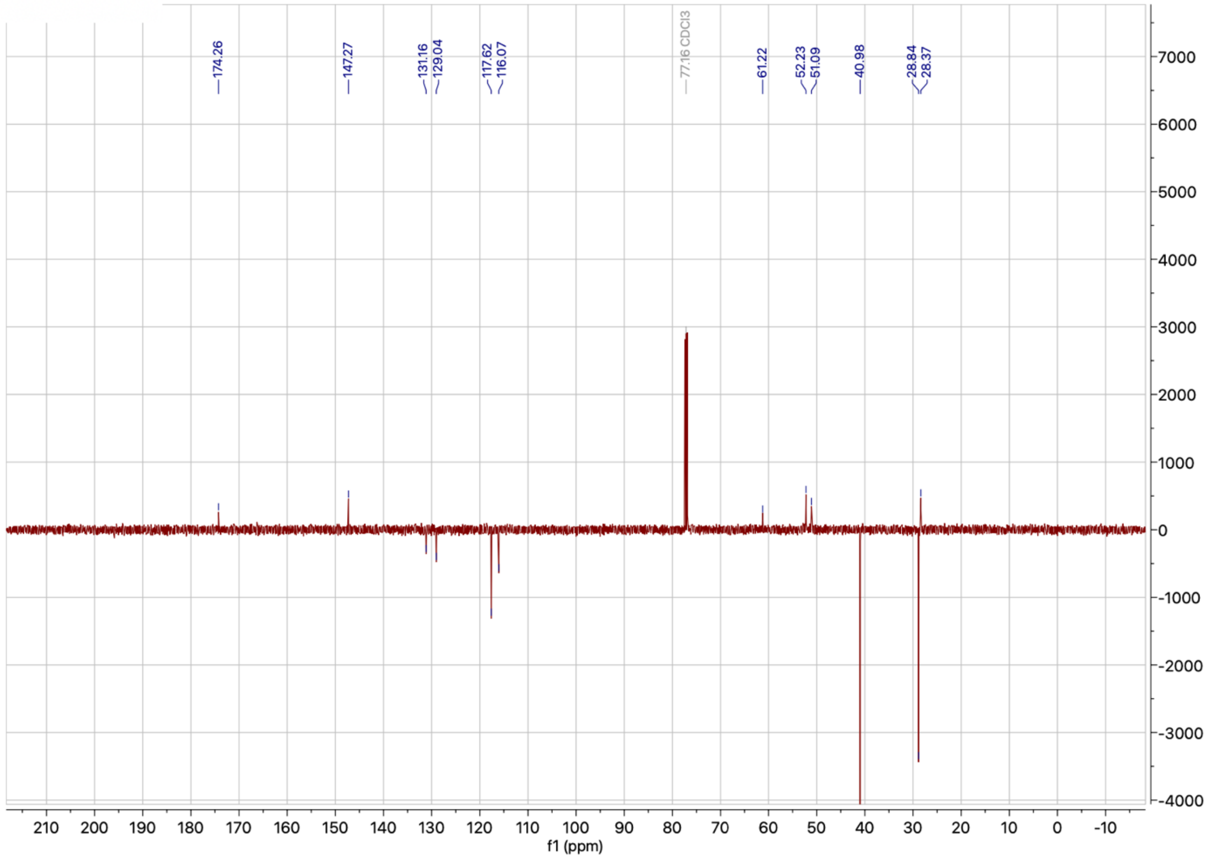

$^{1}H$ NMR spectrum of **8b**

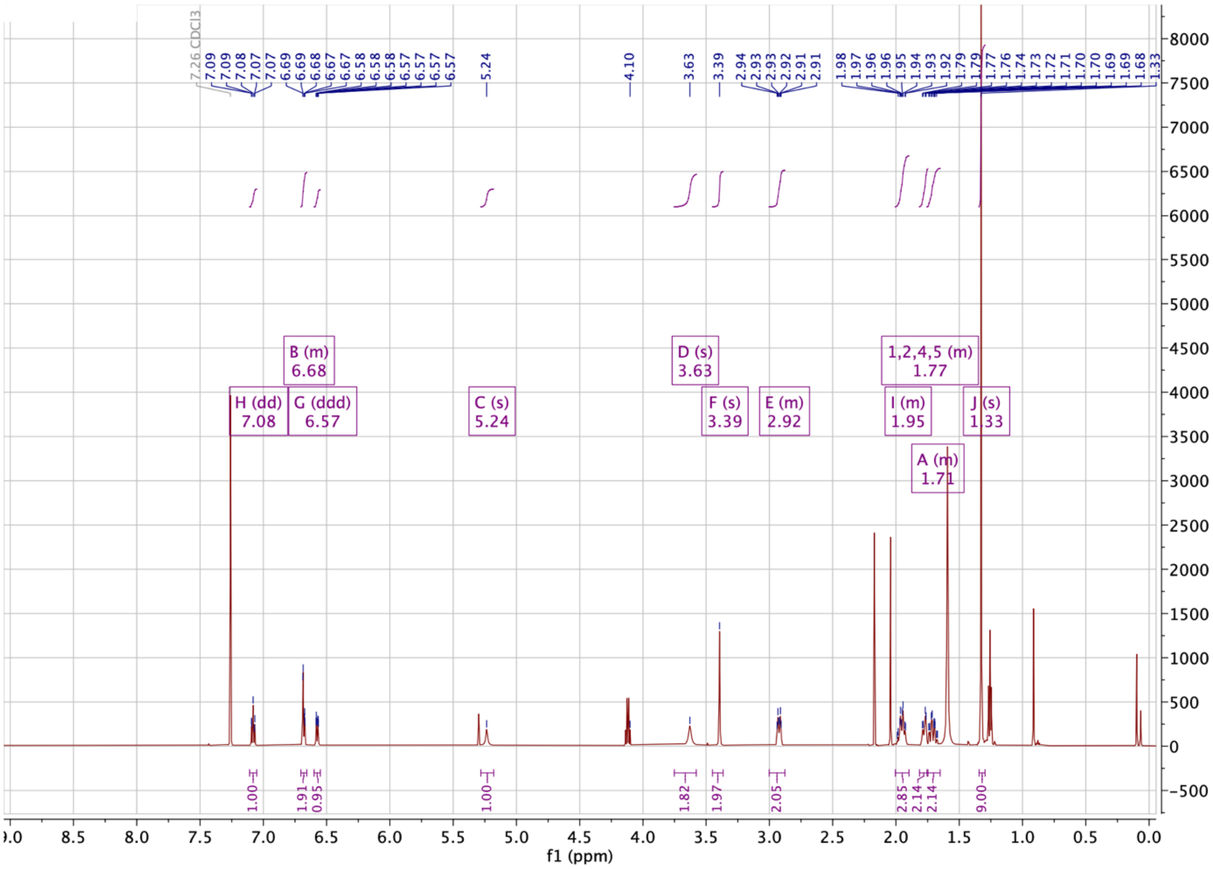


$^{13}C$ NMR spectrum of **8b**

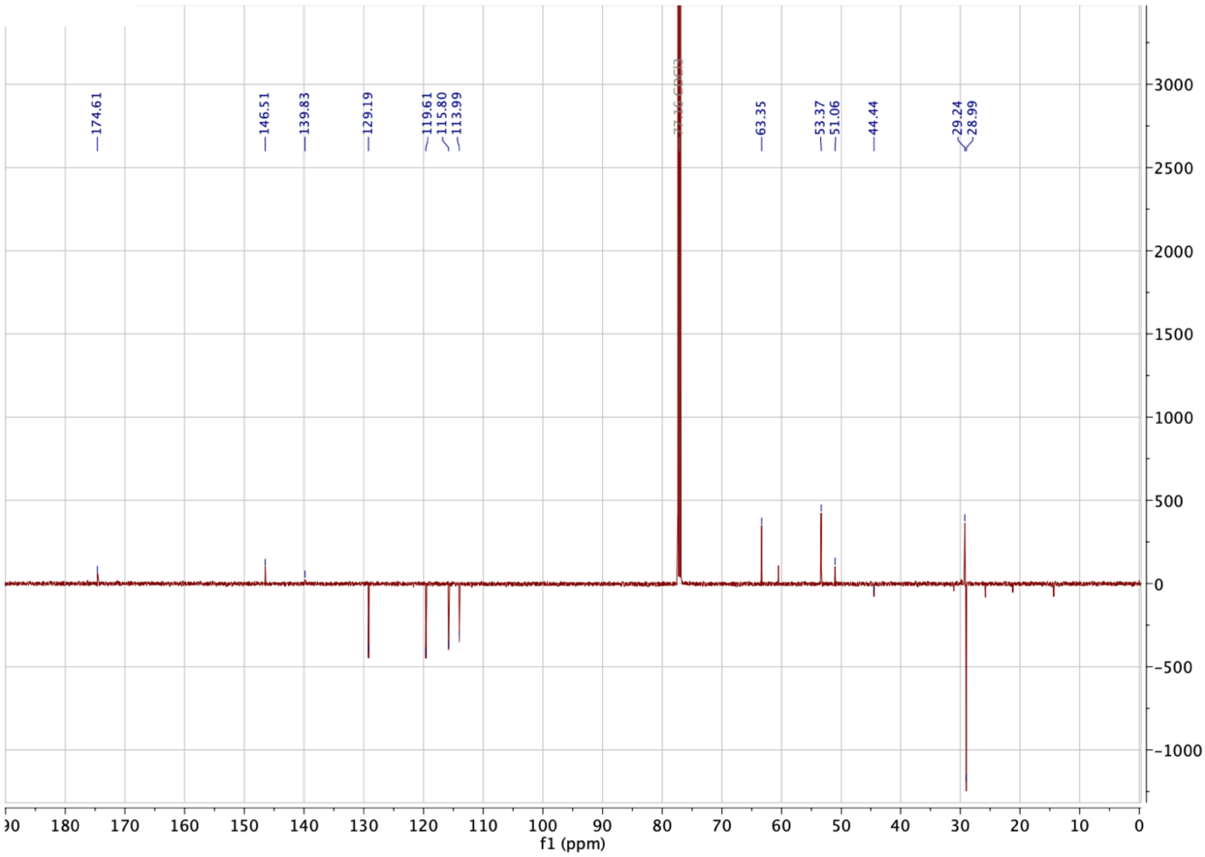

$^1H$ NMR spectrum of **8c**

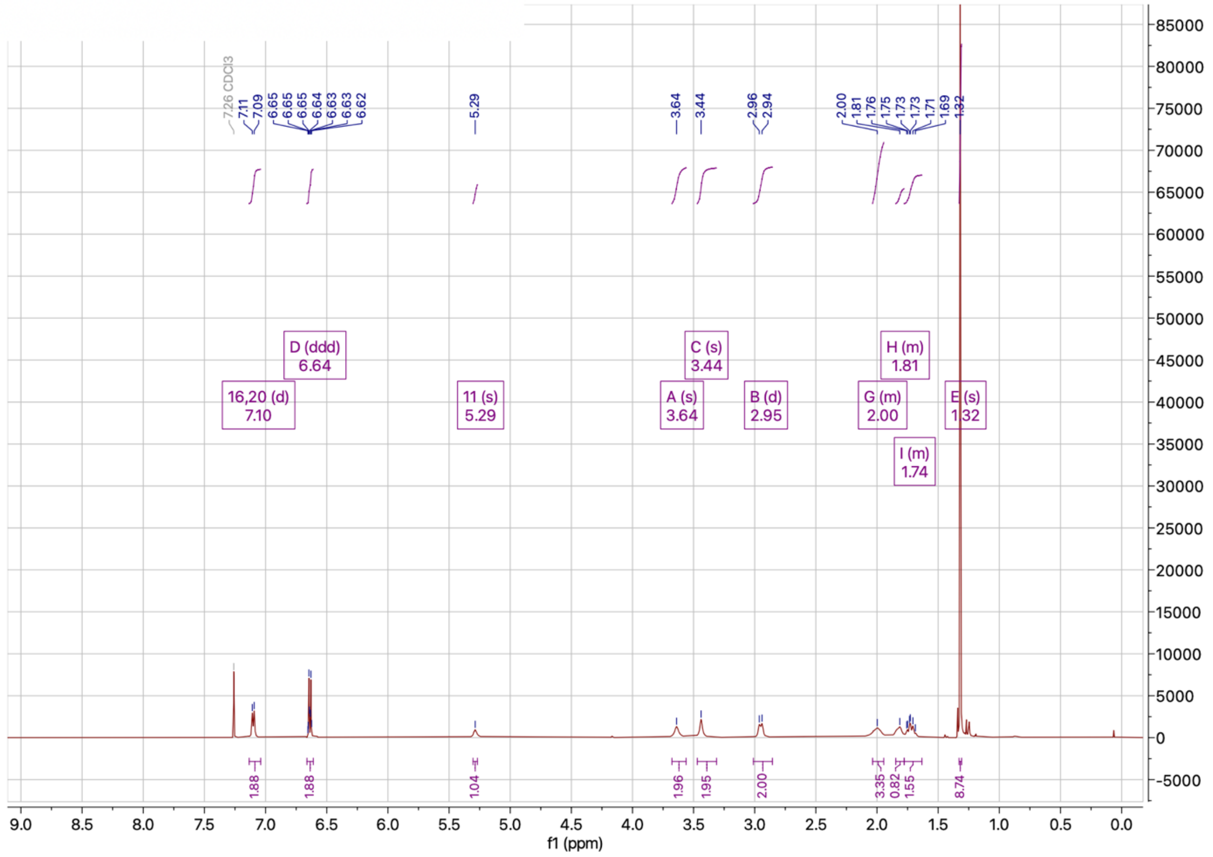


$^{13}C$ NMR spectrum of **8c**

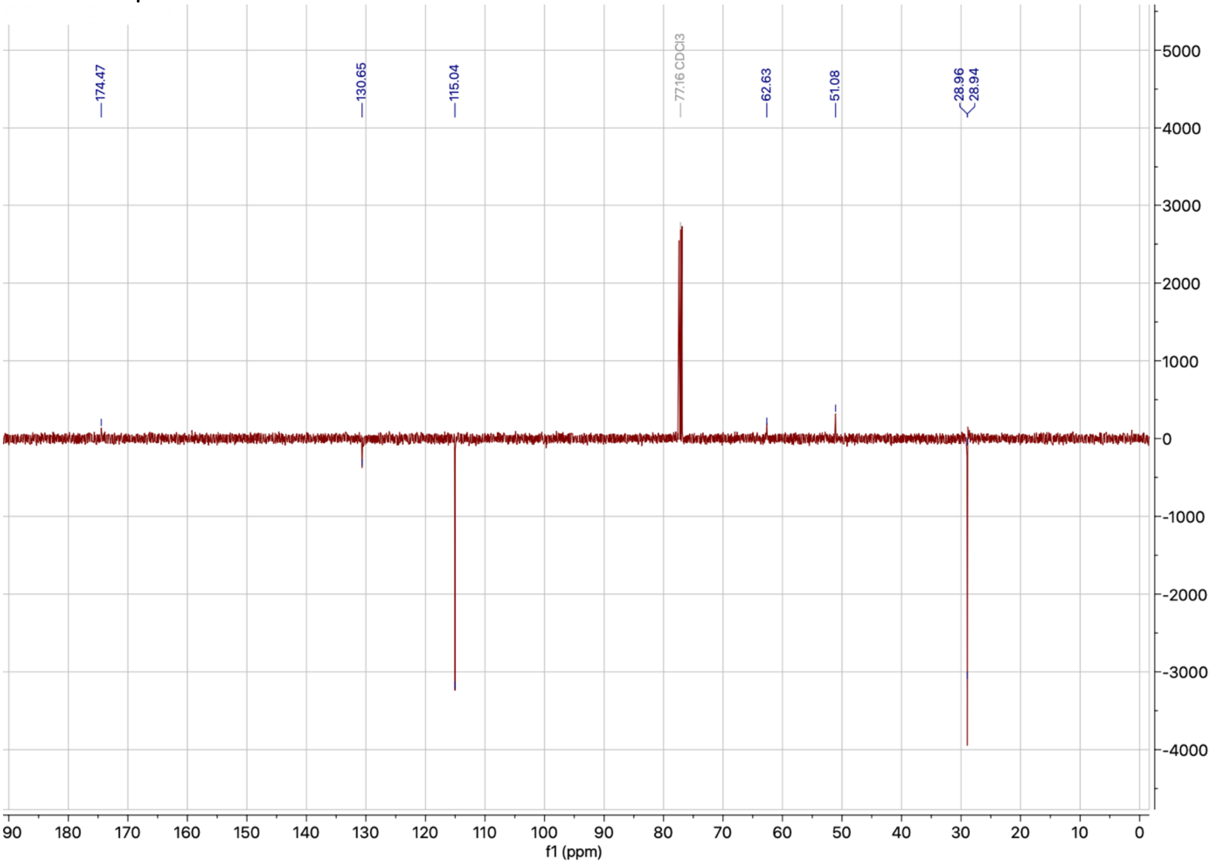

$^{1}$H NMR spectrum of **13a**

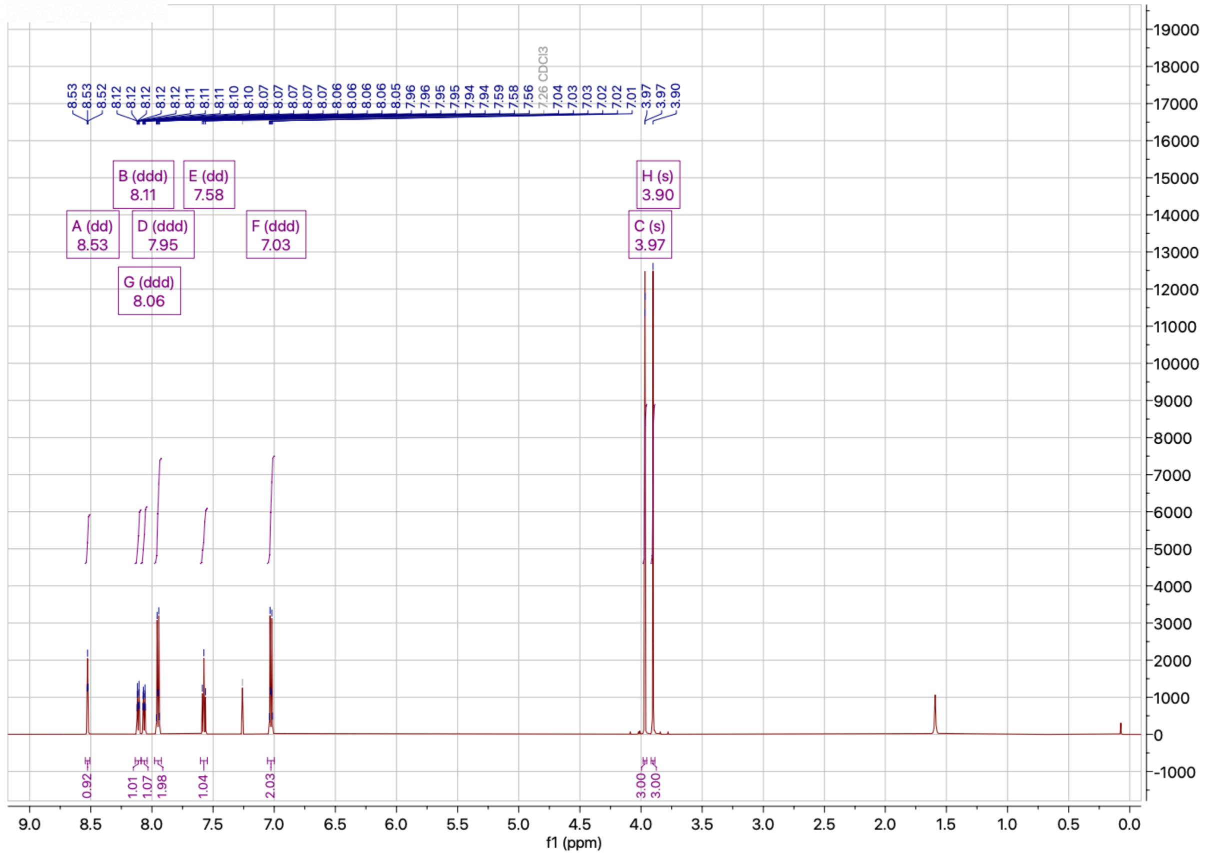


$^{13}$C NMR spectrum of **13a**

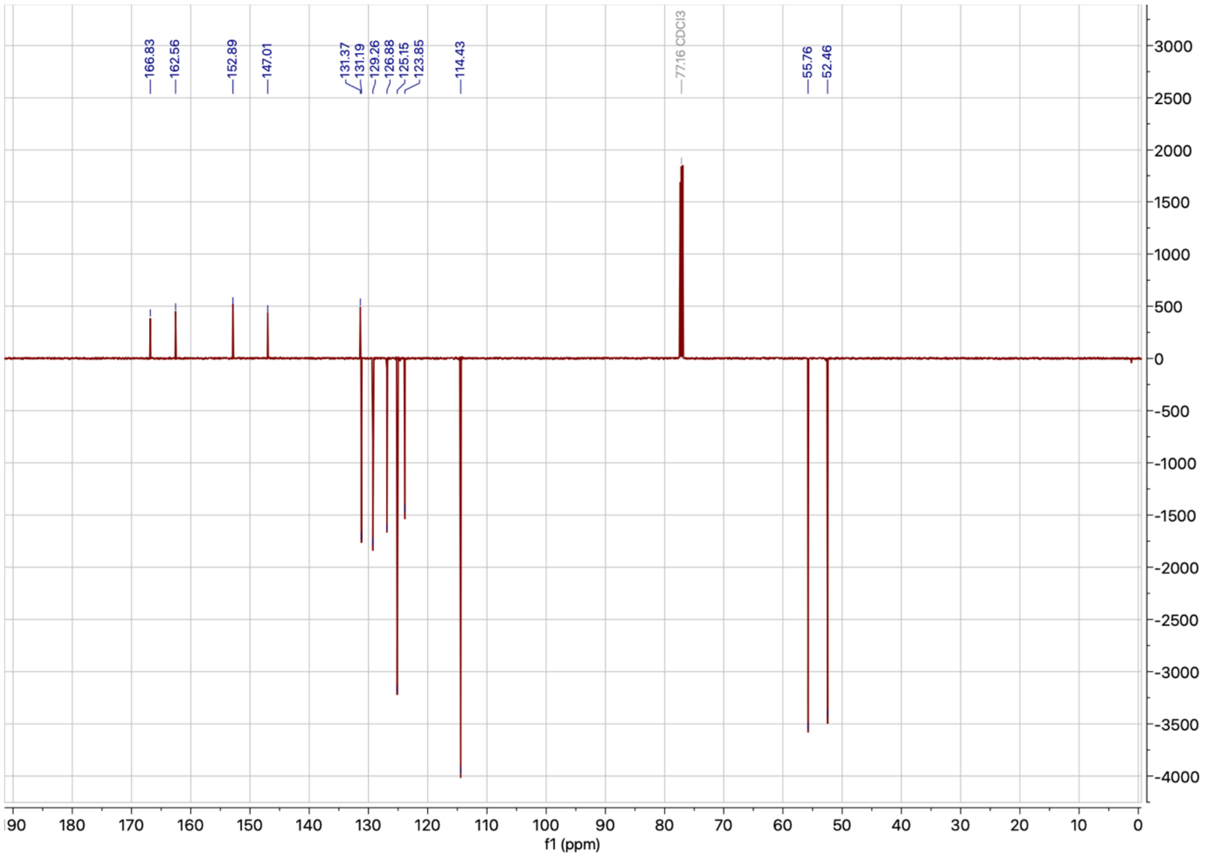

$^1$H NMR spectrum of **14a**

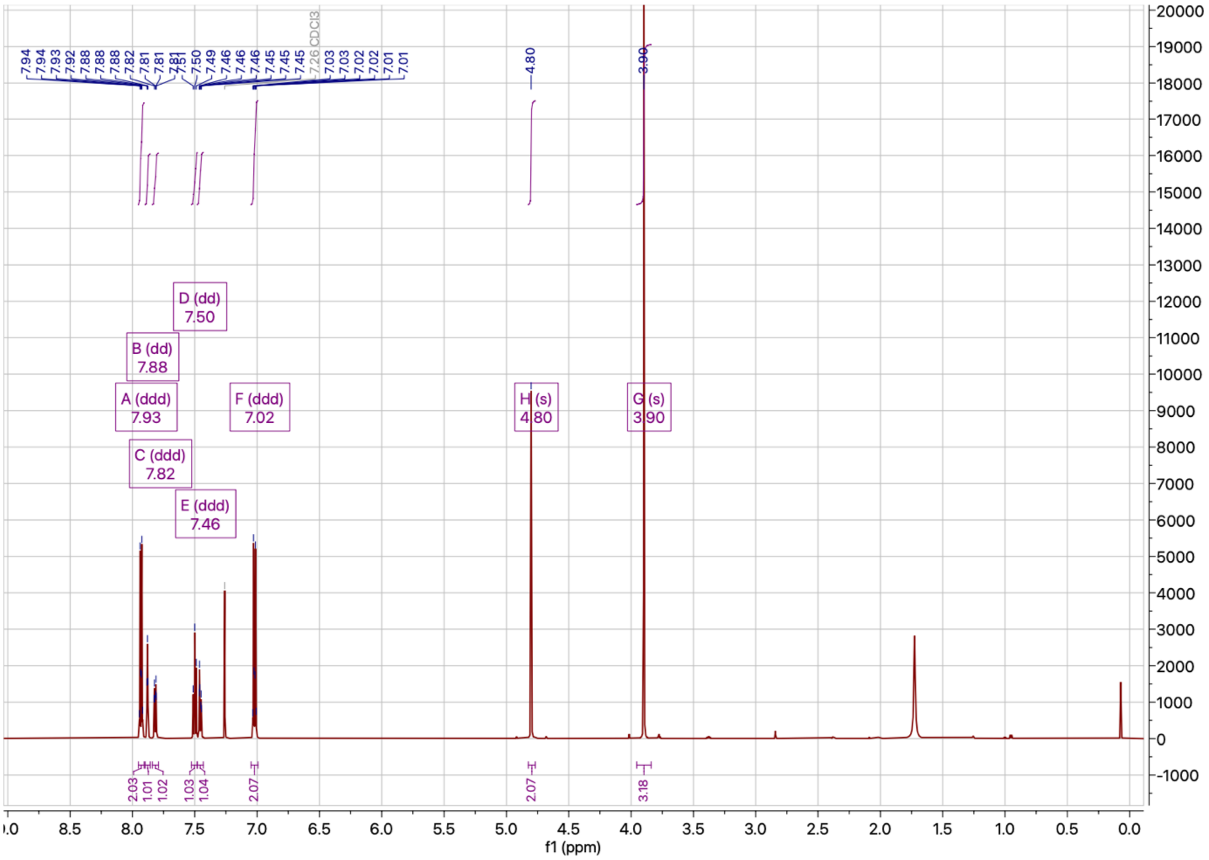


$^{13}$C NMR spectrum of **14a**

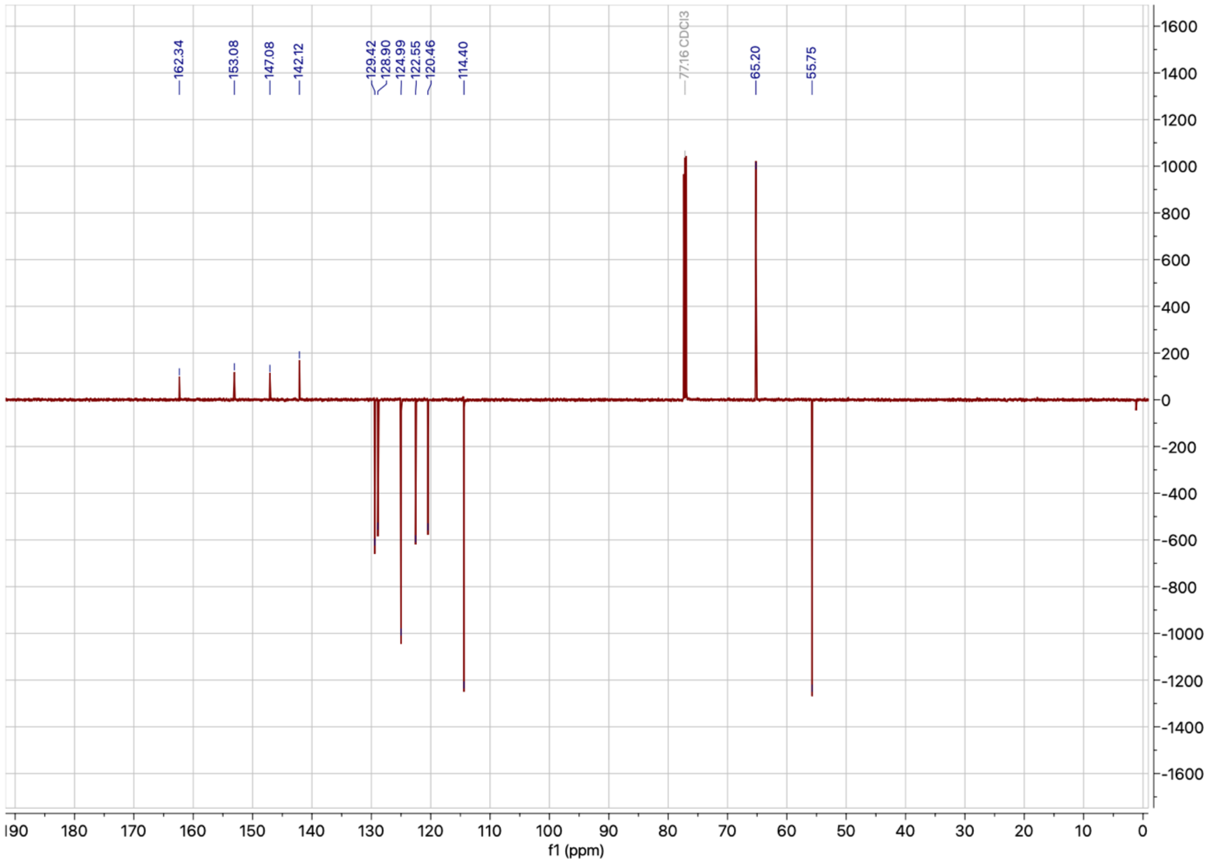

$^{1}$H NMR spectrum of **15a**

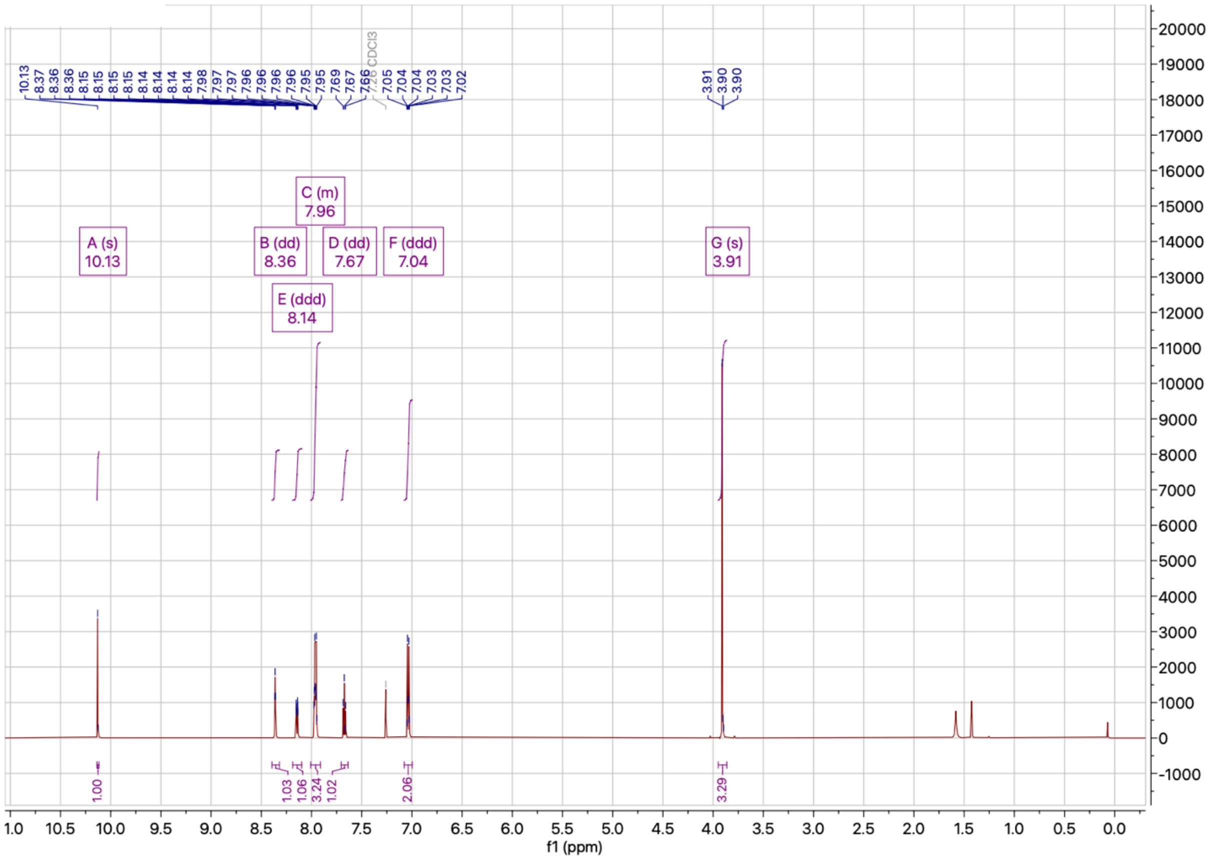


$^{13}$C NMR spectrum of **15a**

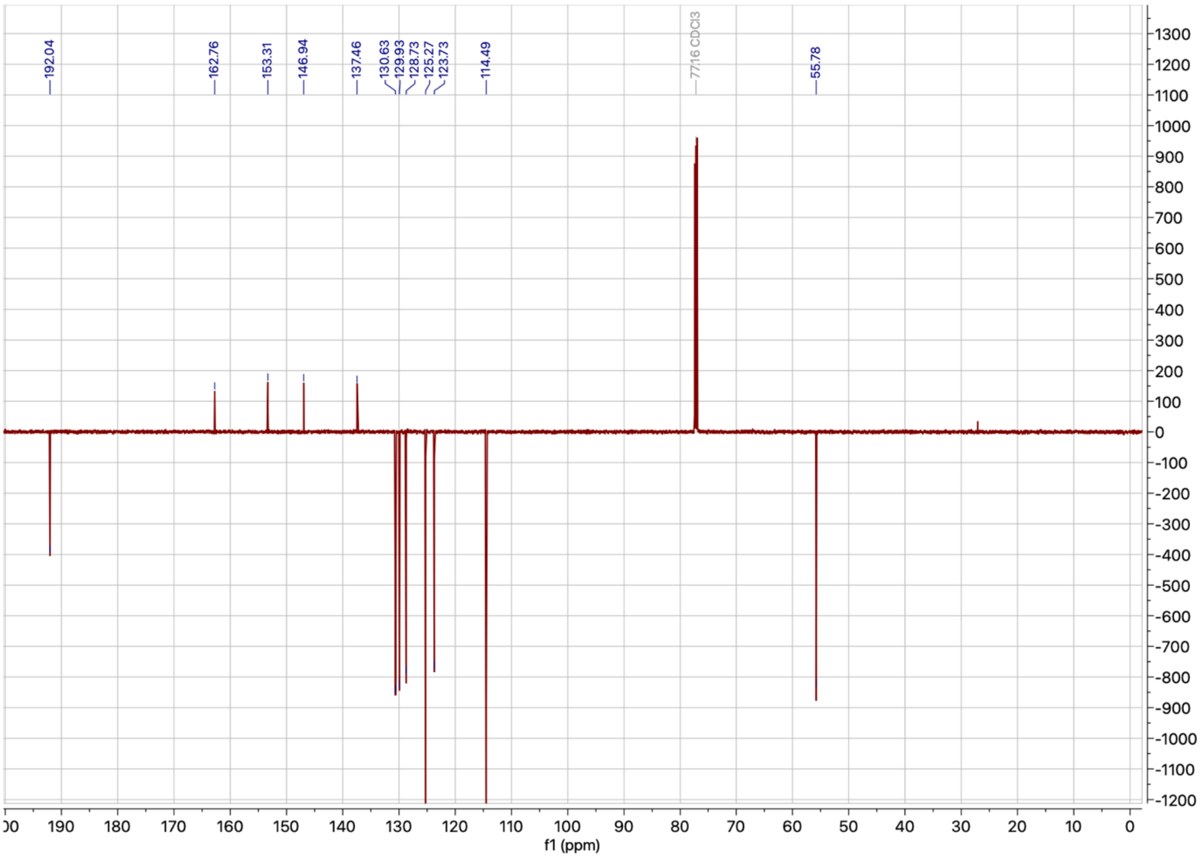

$^{1}$H NMR spectrum of **13b**

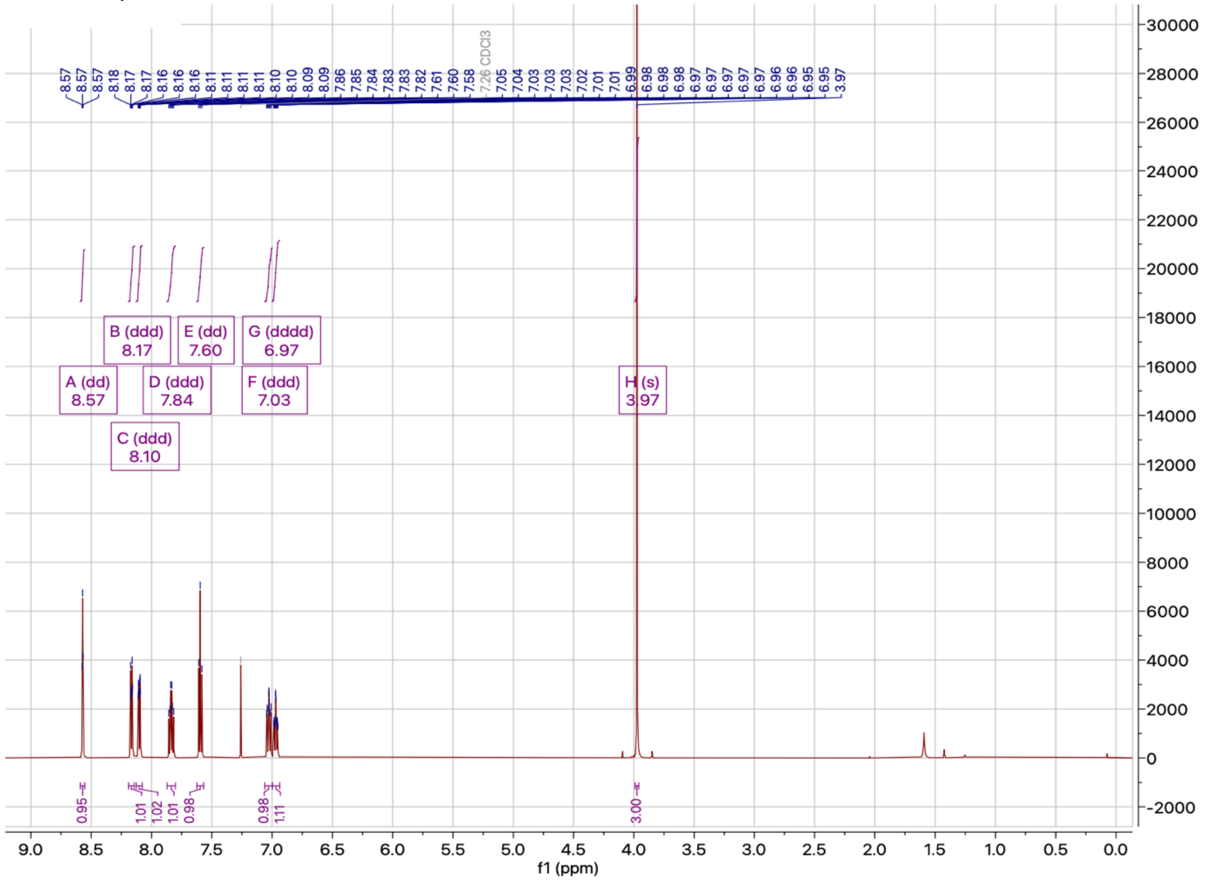


$^{13}$C NMR spectrum of **13b**

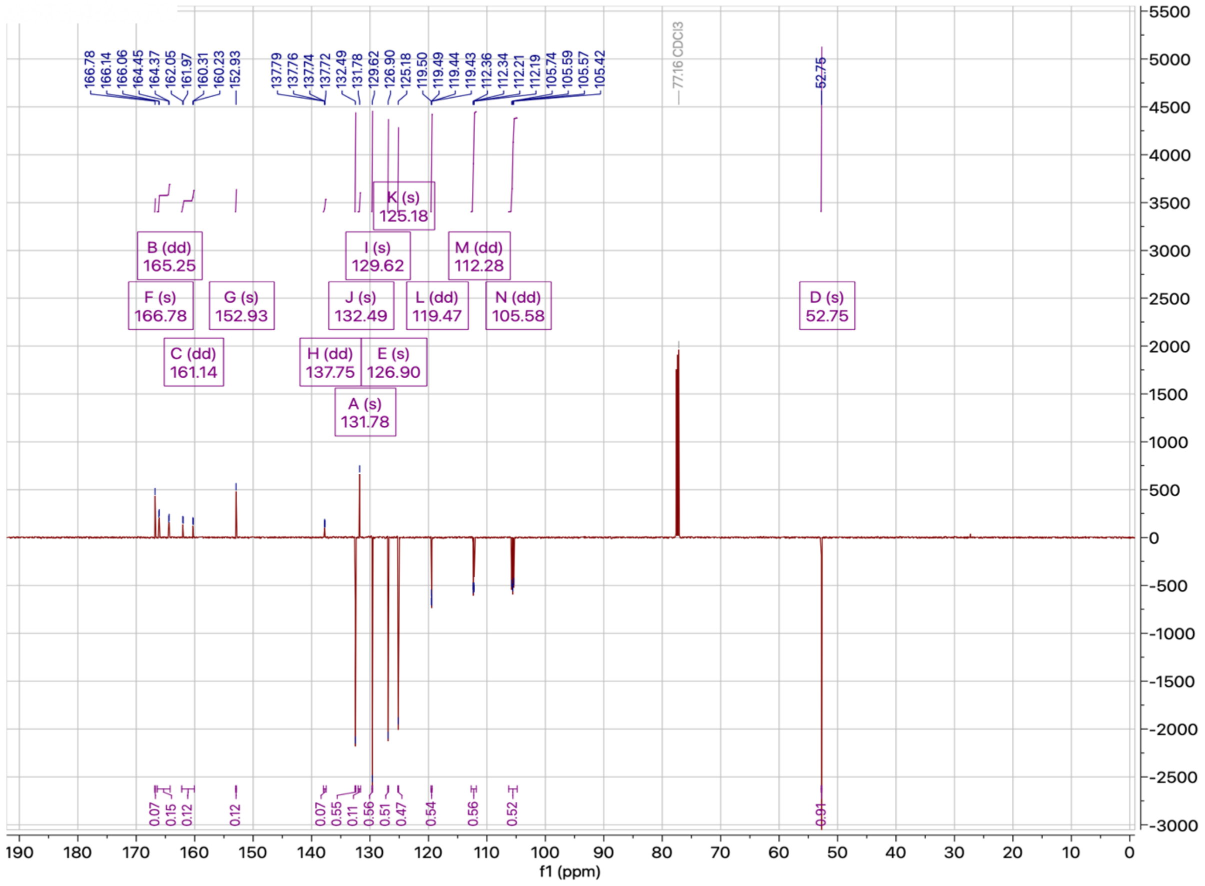

$^{1}H$ NMR spectrum of **14b**

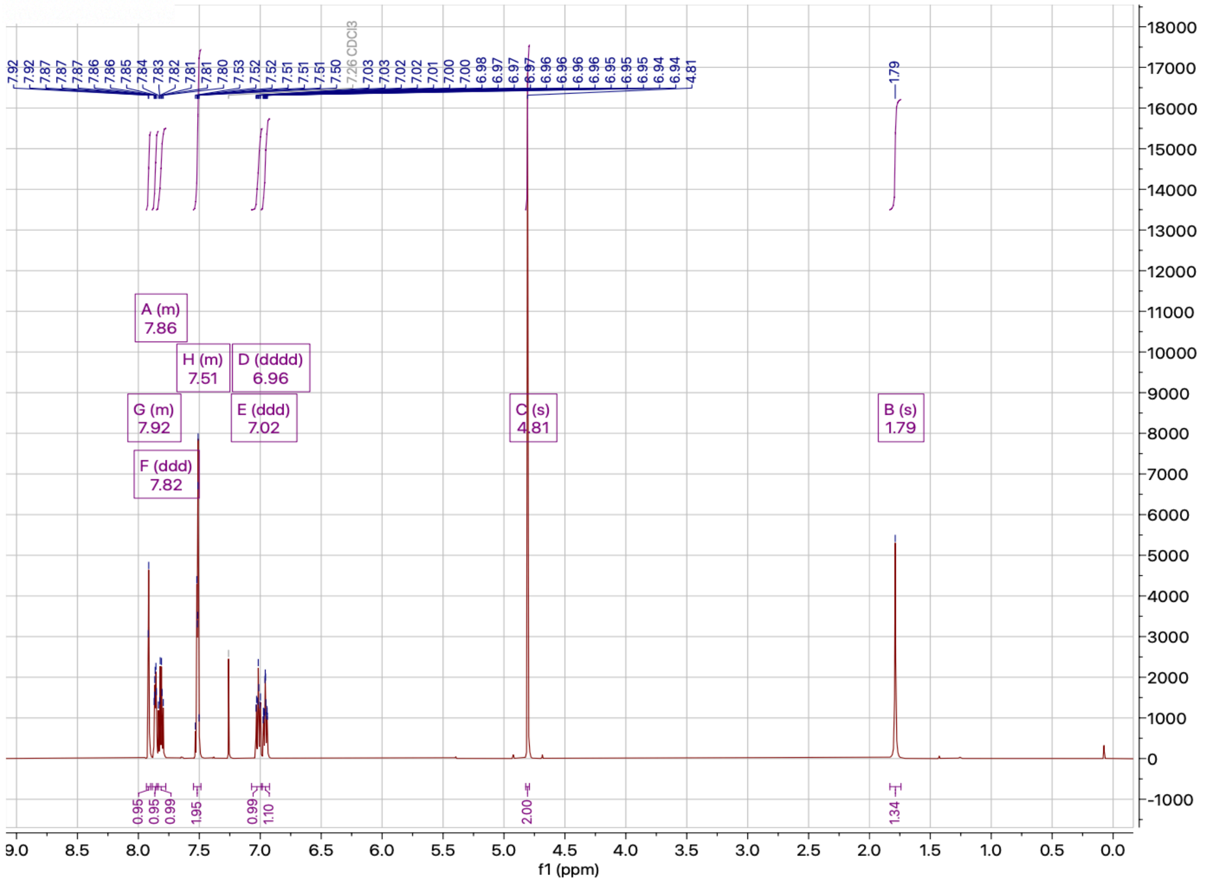


$^{13}C$ NMR spectrum of **14b**

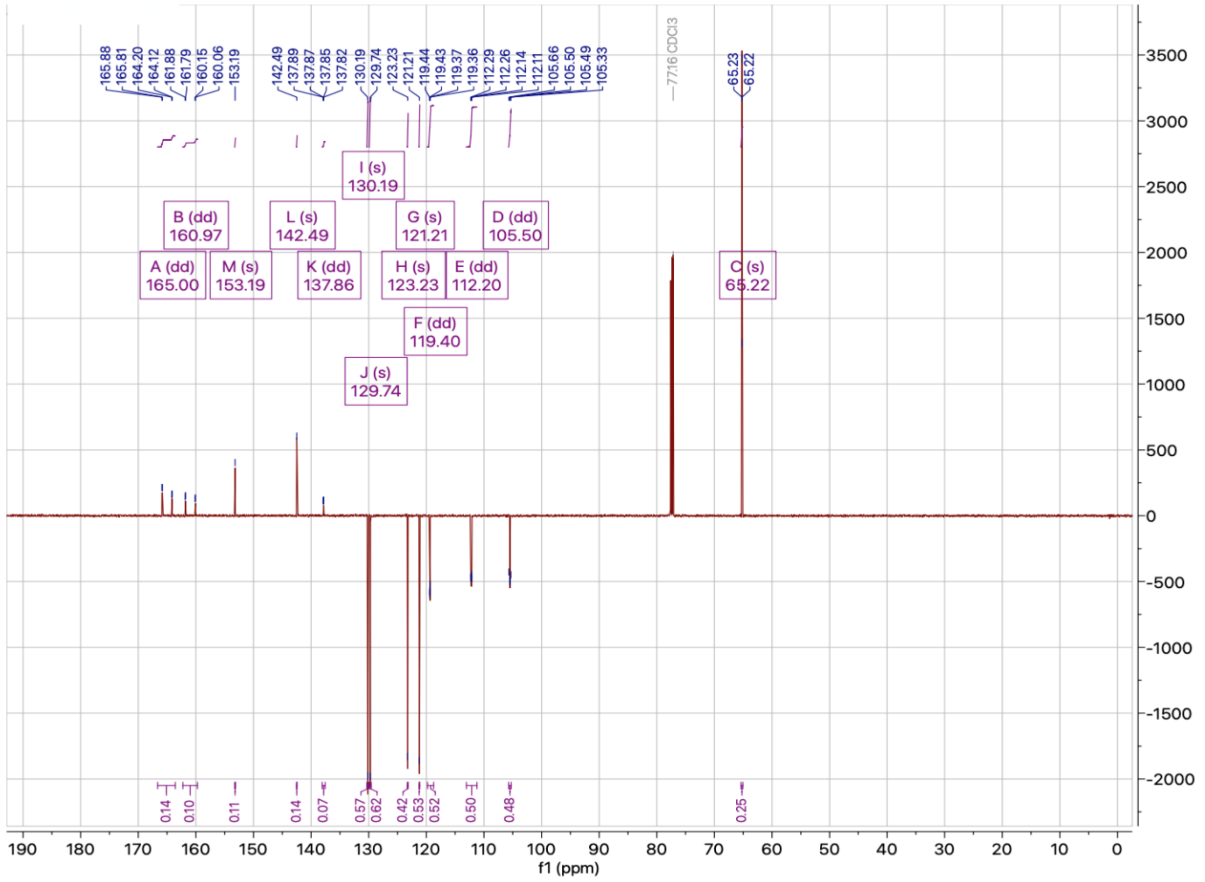

$^{1}H$ NMR spectrum of **15b**

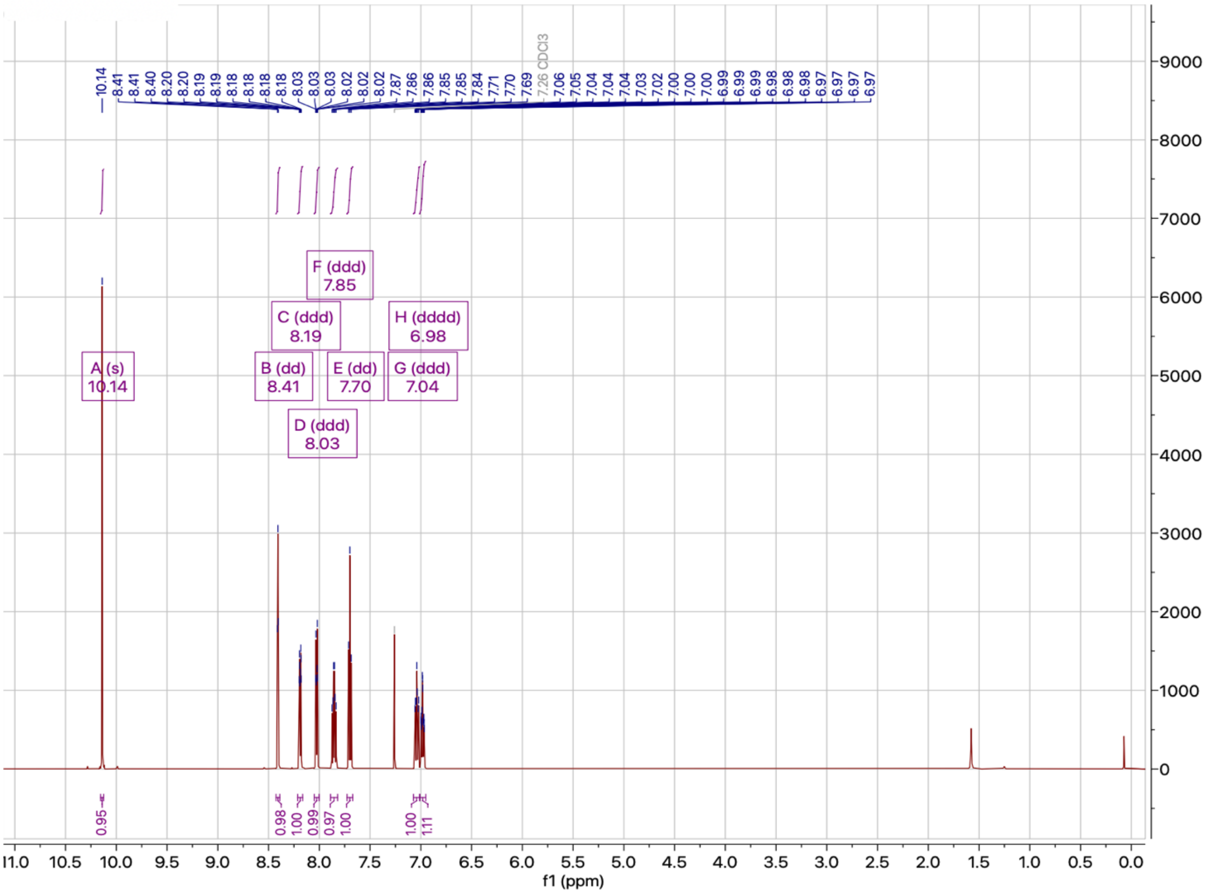

$^{13}C$ NMR spectrum of **15b**

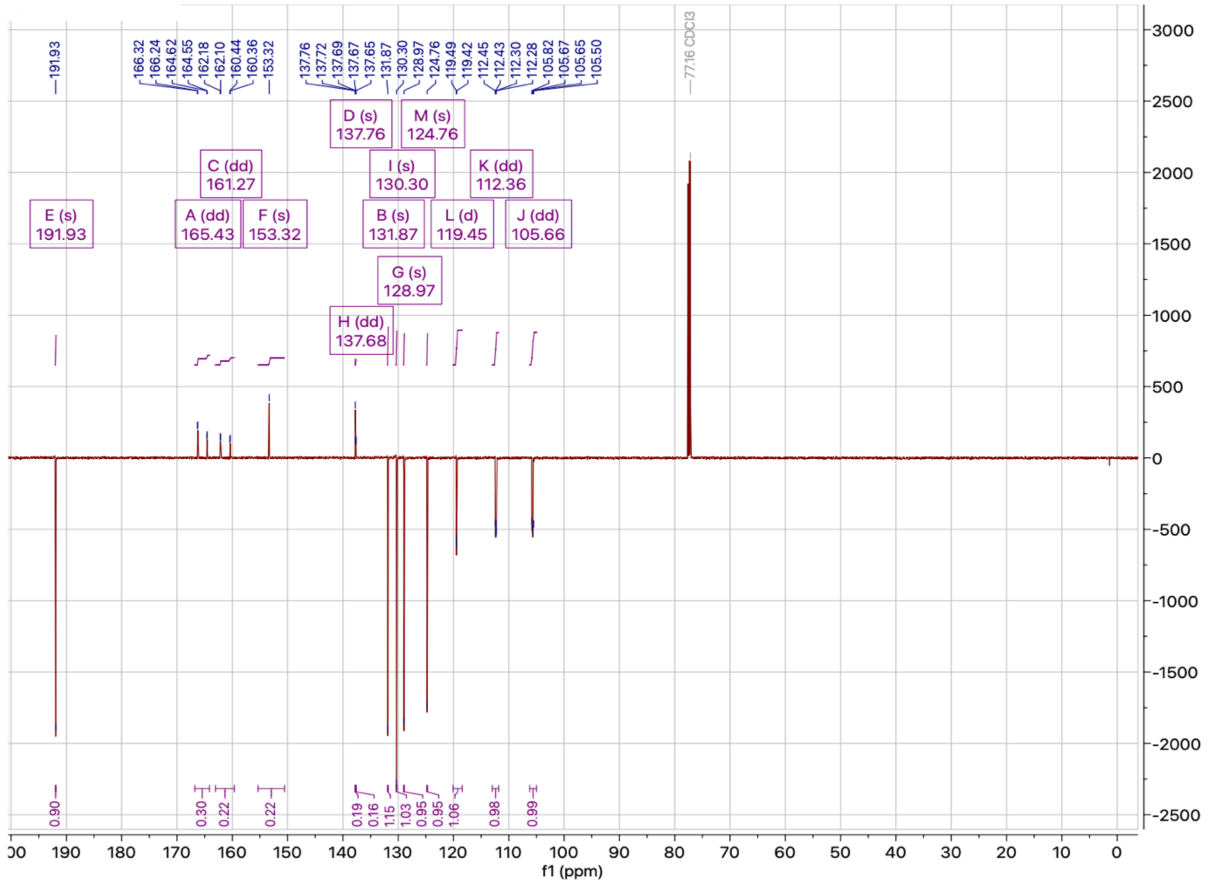

$^{1}$H NMR spectrum of **3a**

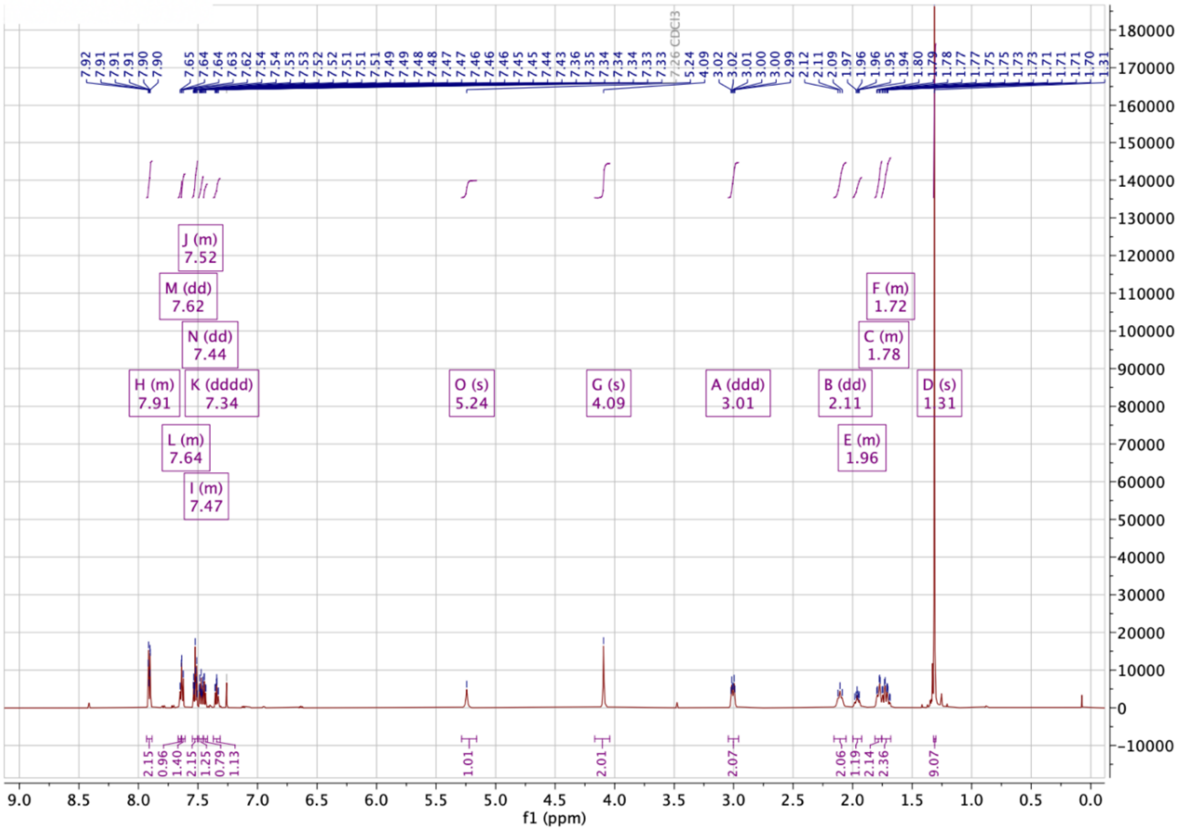


$^{13}$C NMR spectrum of **3a**

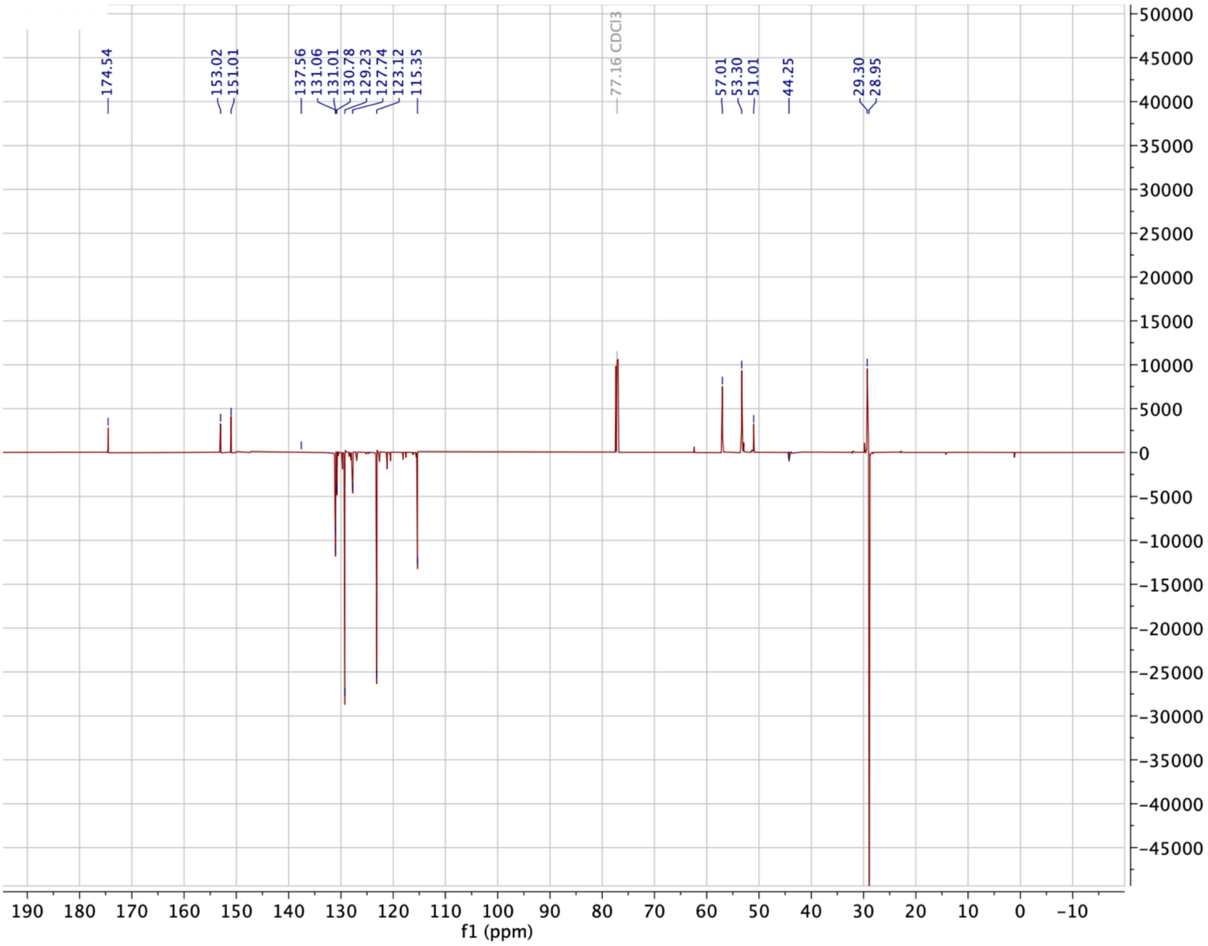

[1]H NMR spectrum of **3b**

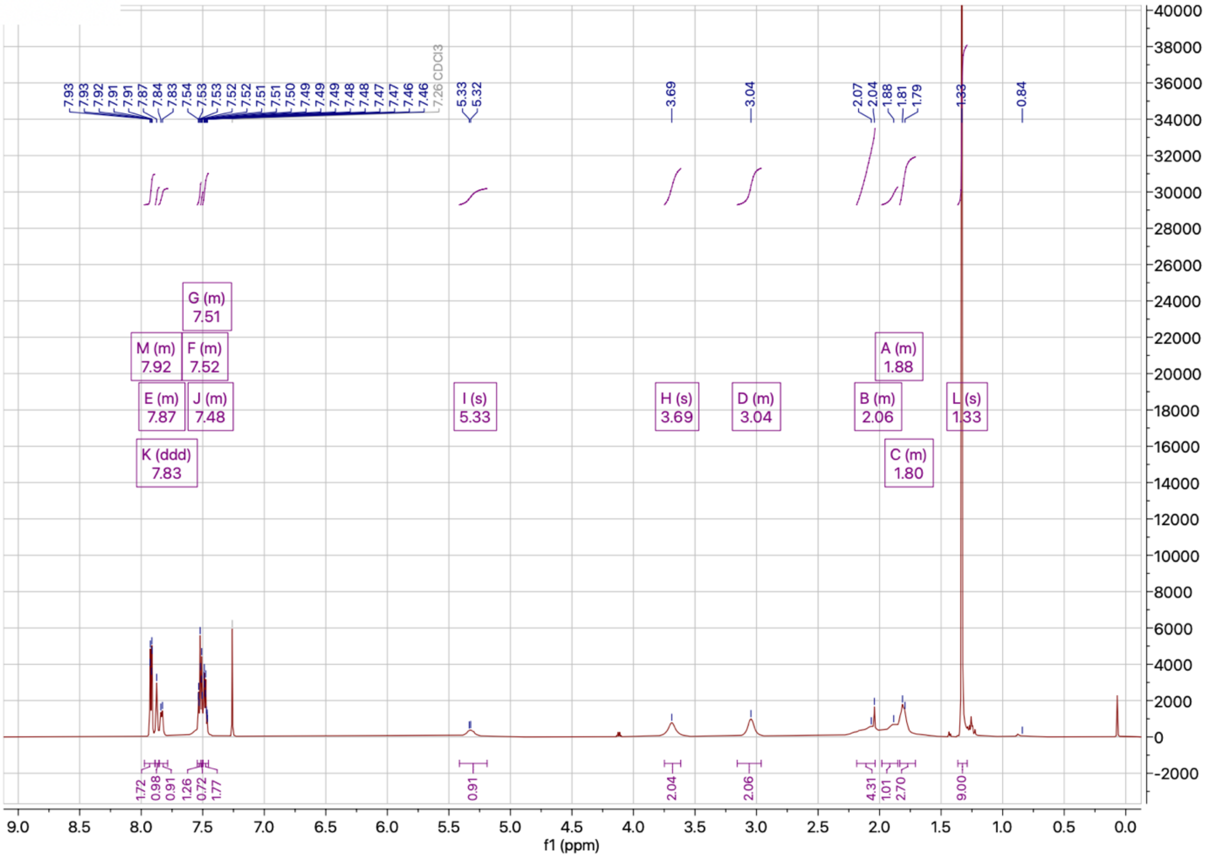

[13]C NMR spectrum of **3b**

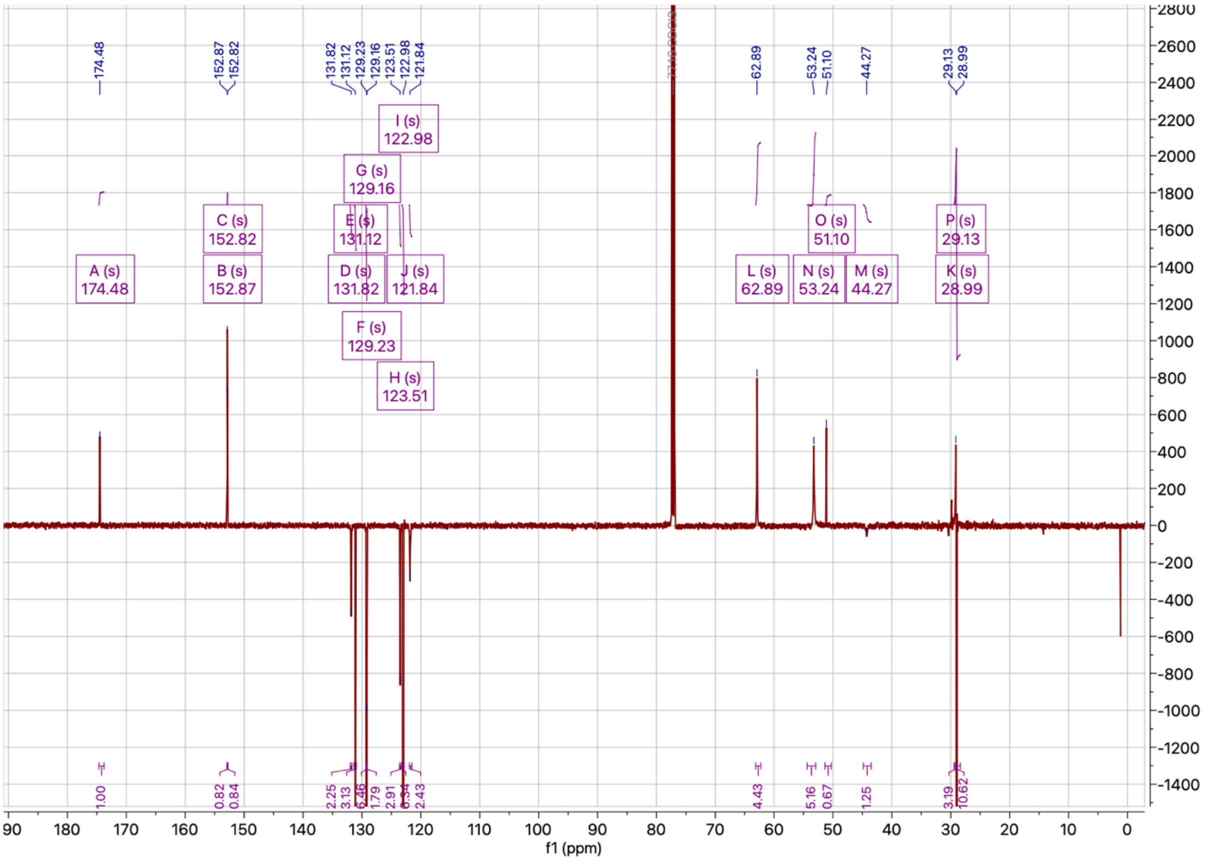

[1]H NMR spectrum of **3c**

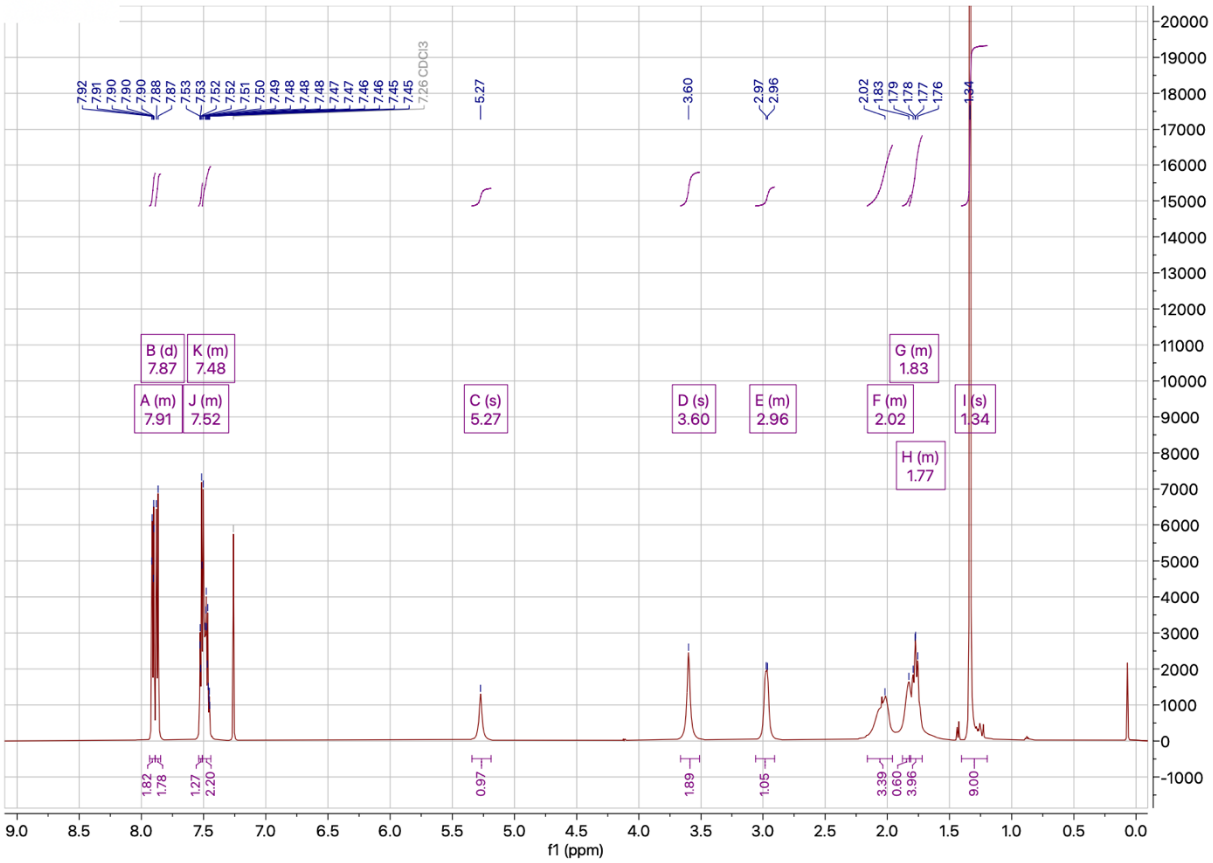

[13]C NMR spectrum of **3c**

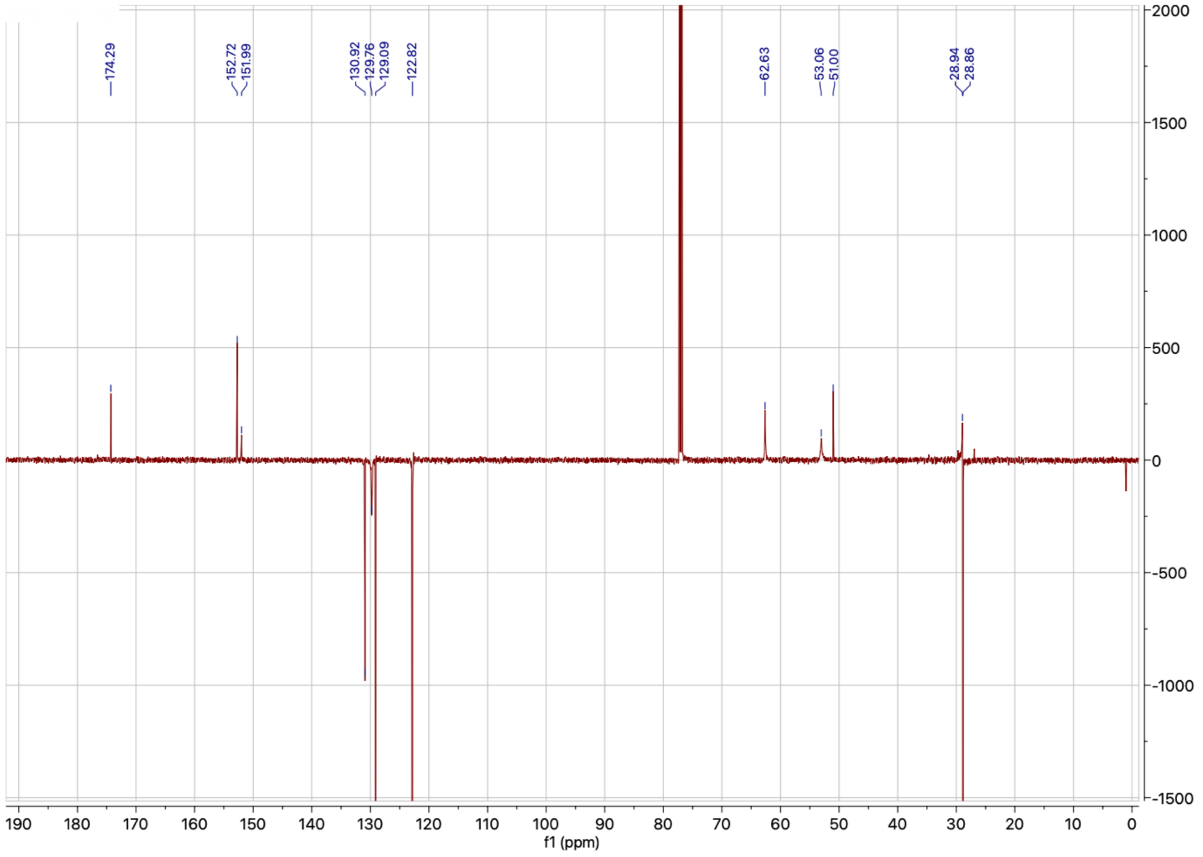

$^{1}$H NMR spectrum of **3d**

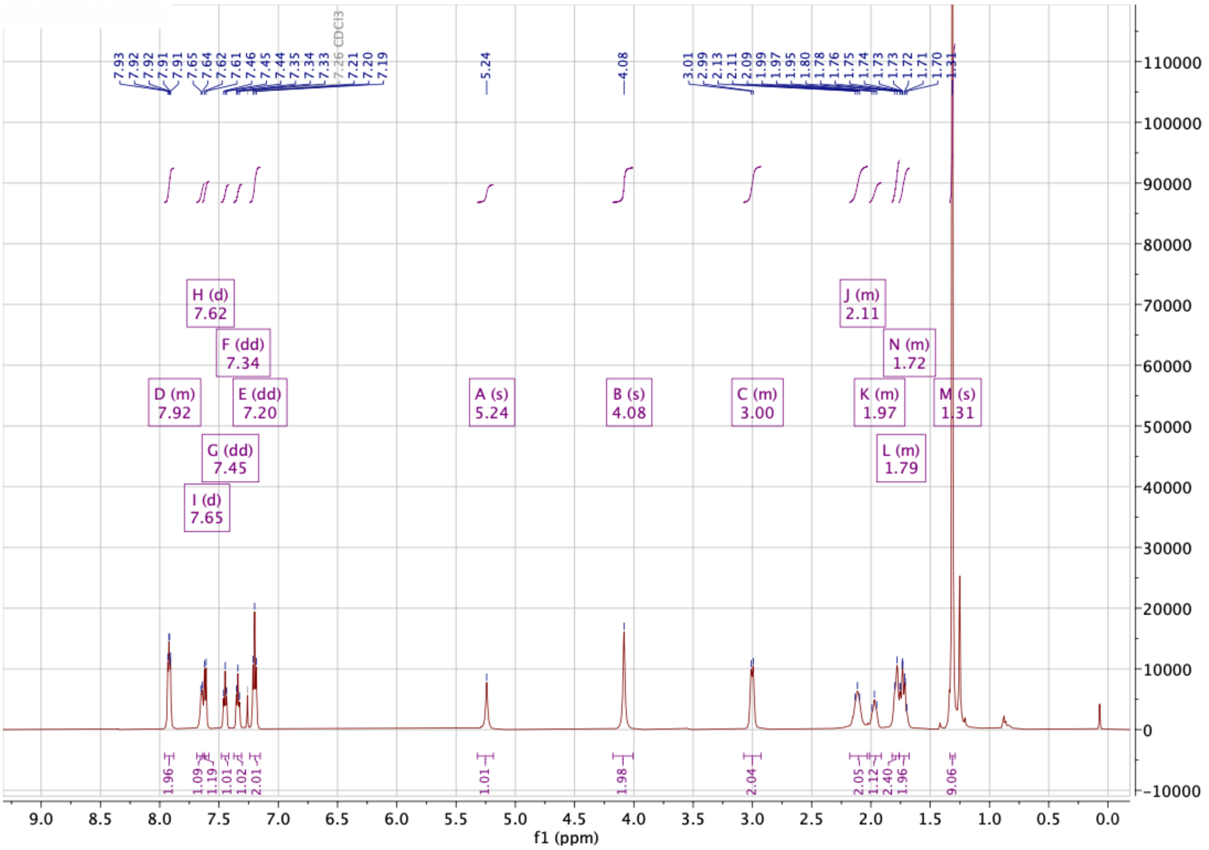

$^{13}$C NMR spectrum of **3d**

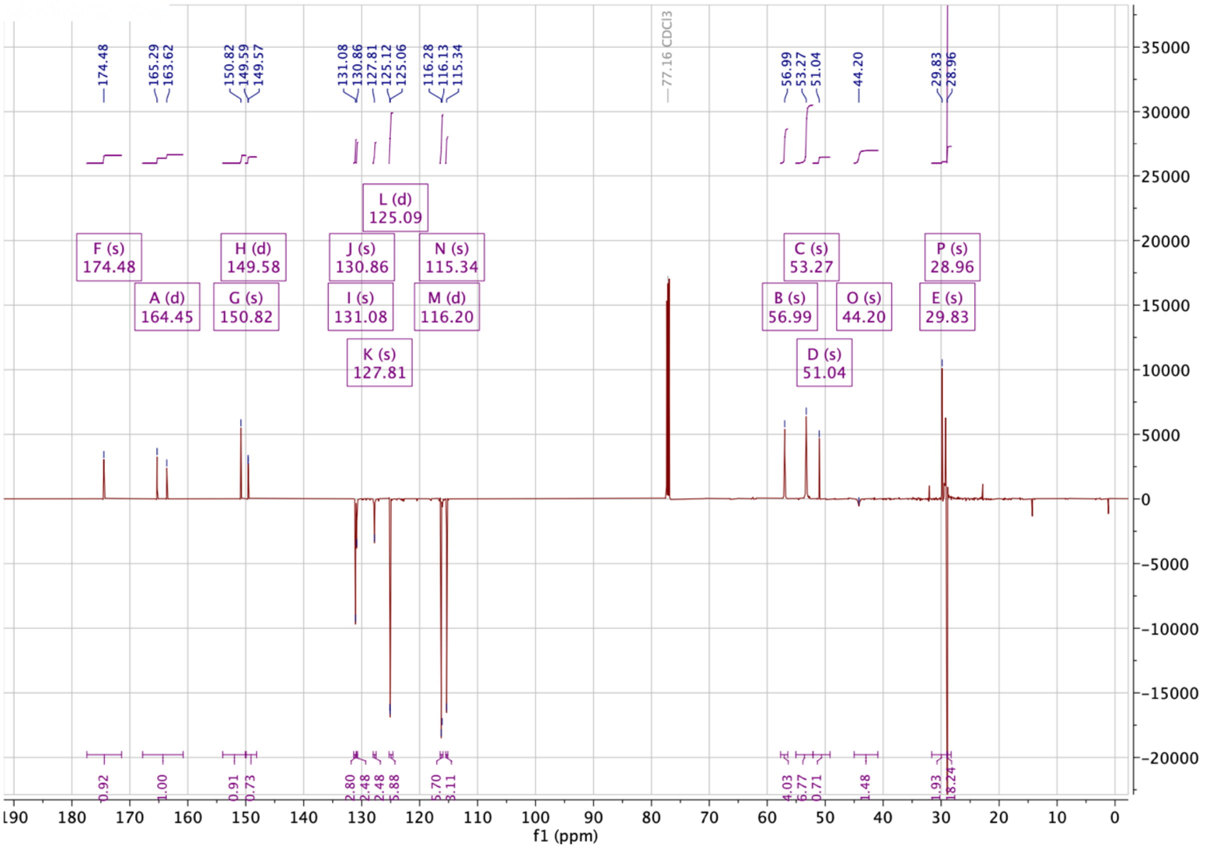

$^1$H NMR spectrum of **3e**

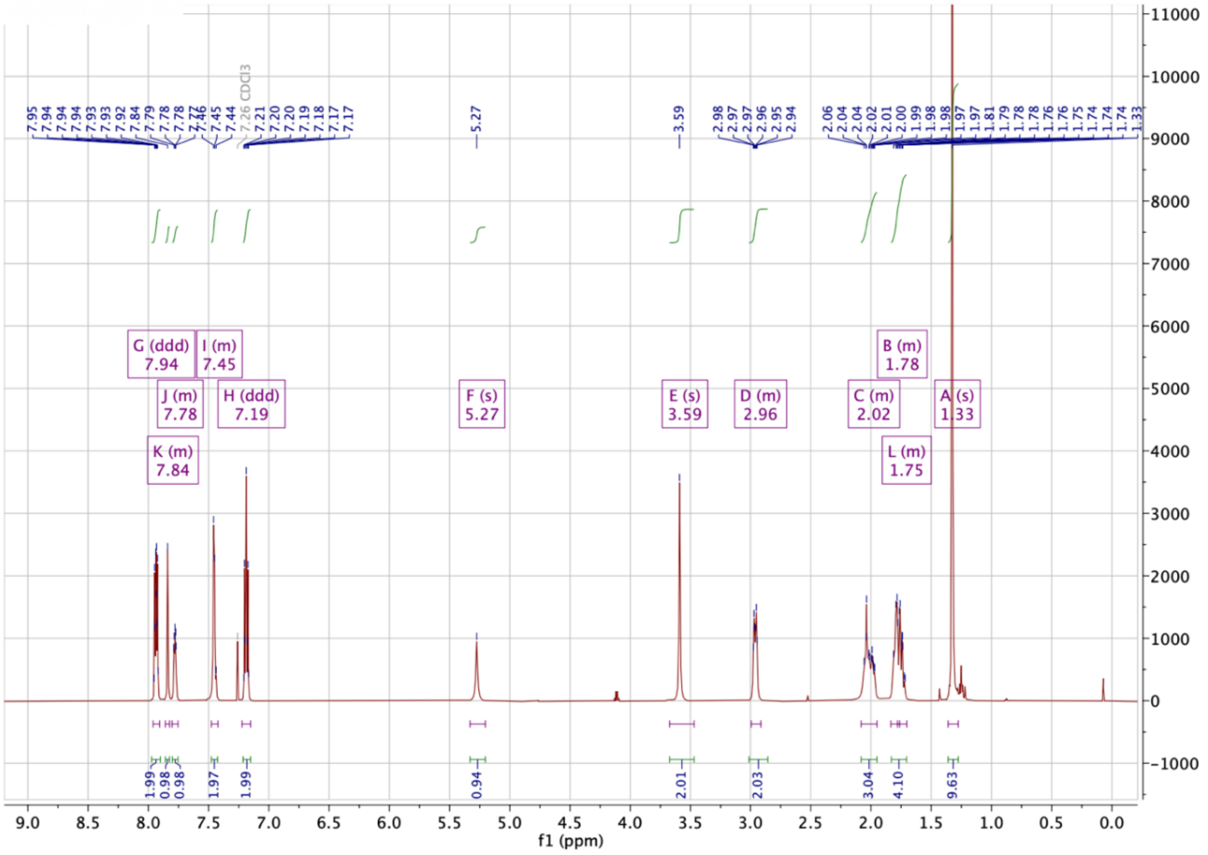


$^{13}$C NMR spectrum of **3e**

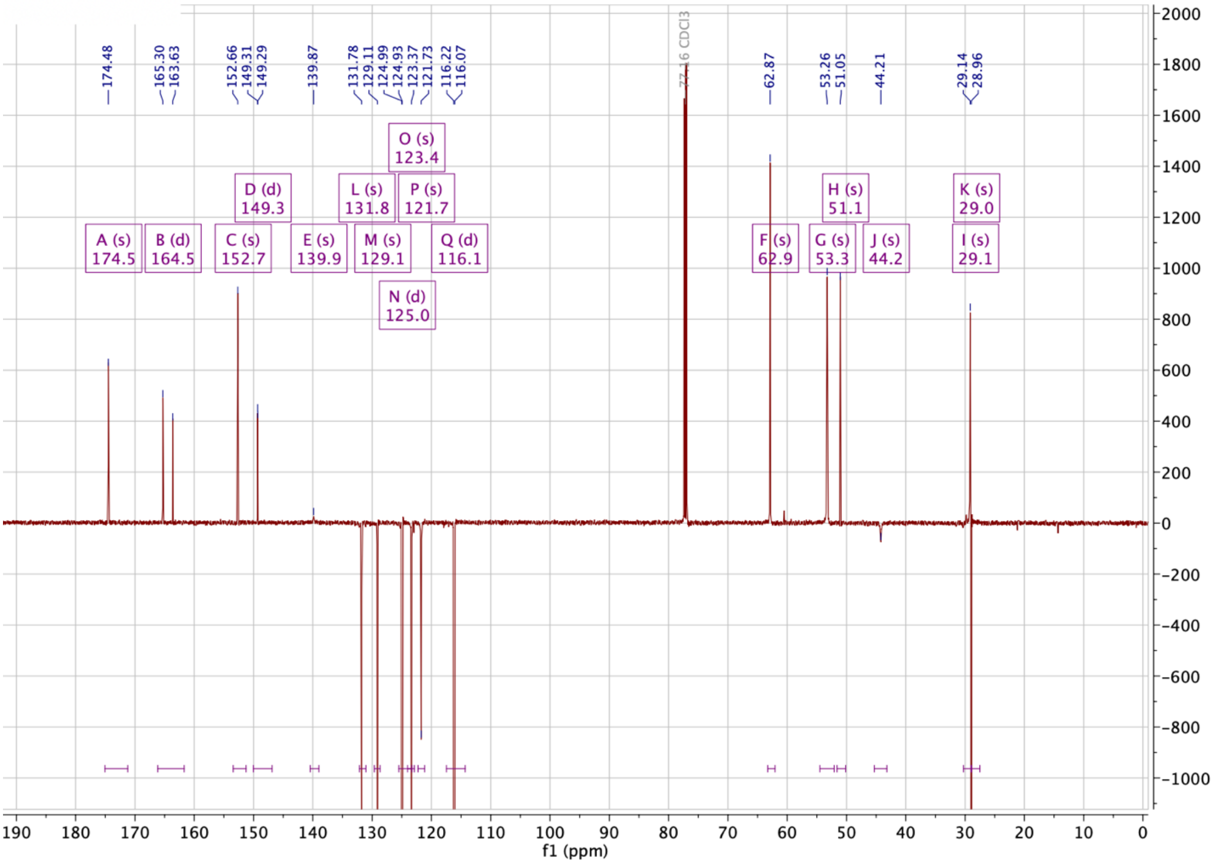

$^{1}$H NMR spectrum of **3f**

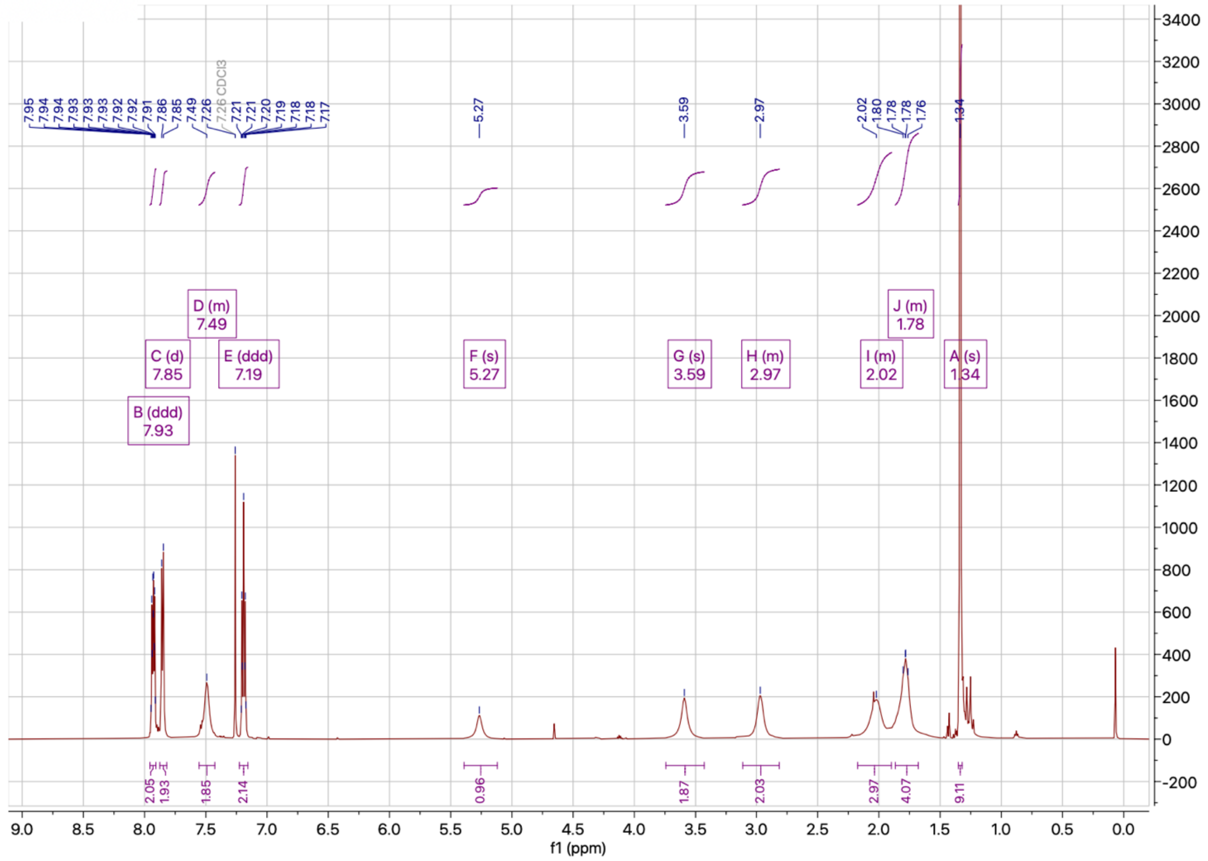


$^{13}$C NMR spectrum of **3f**

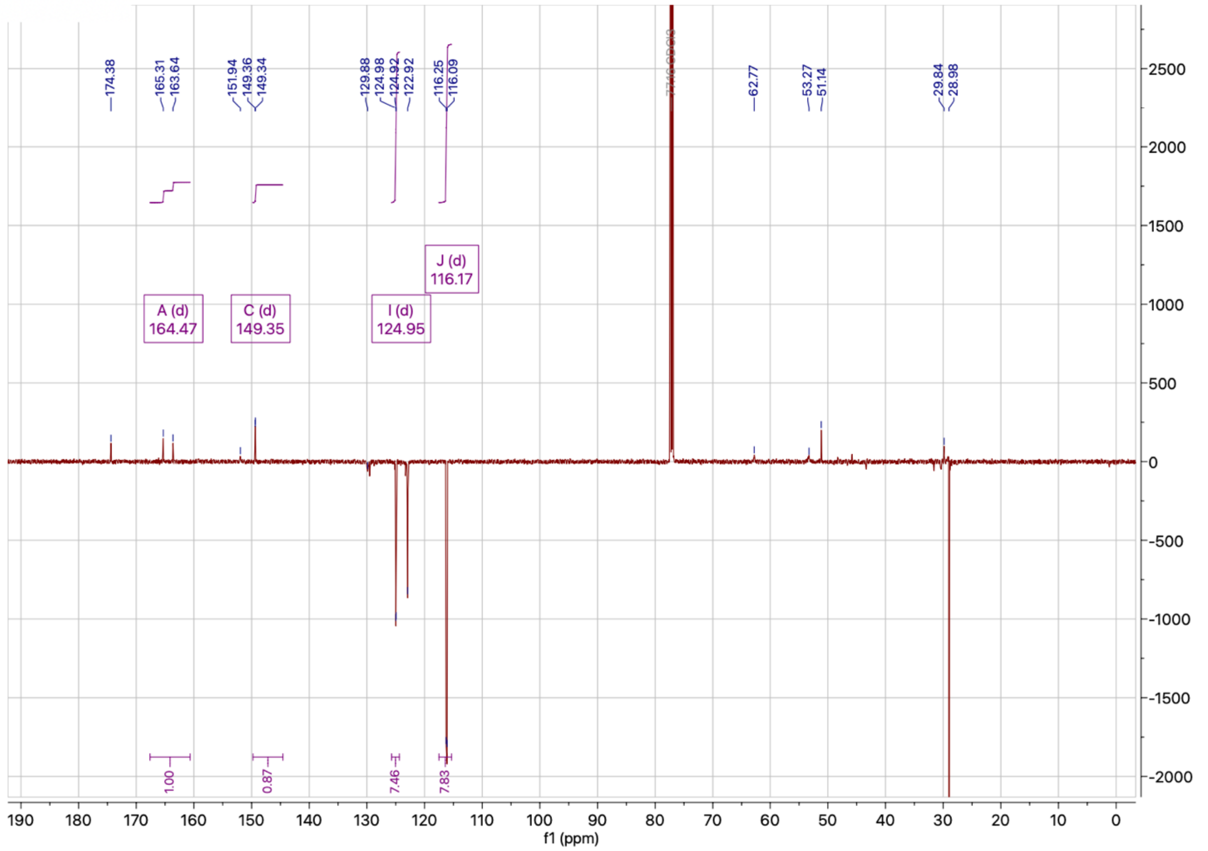

[1]H NMR spectrum of **3g**

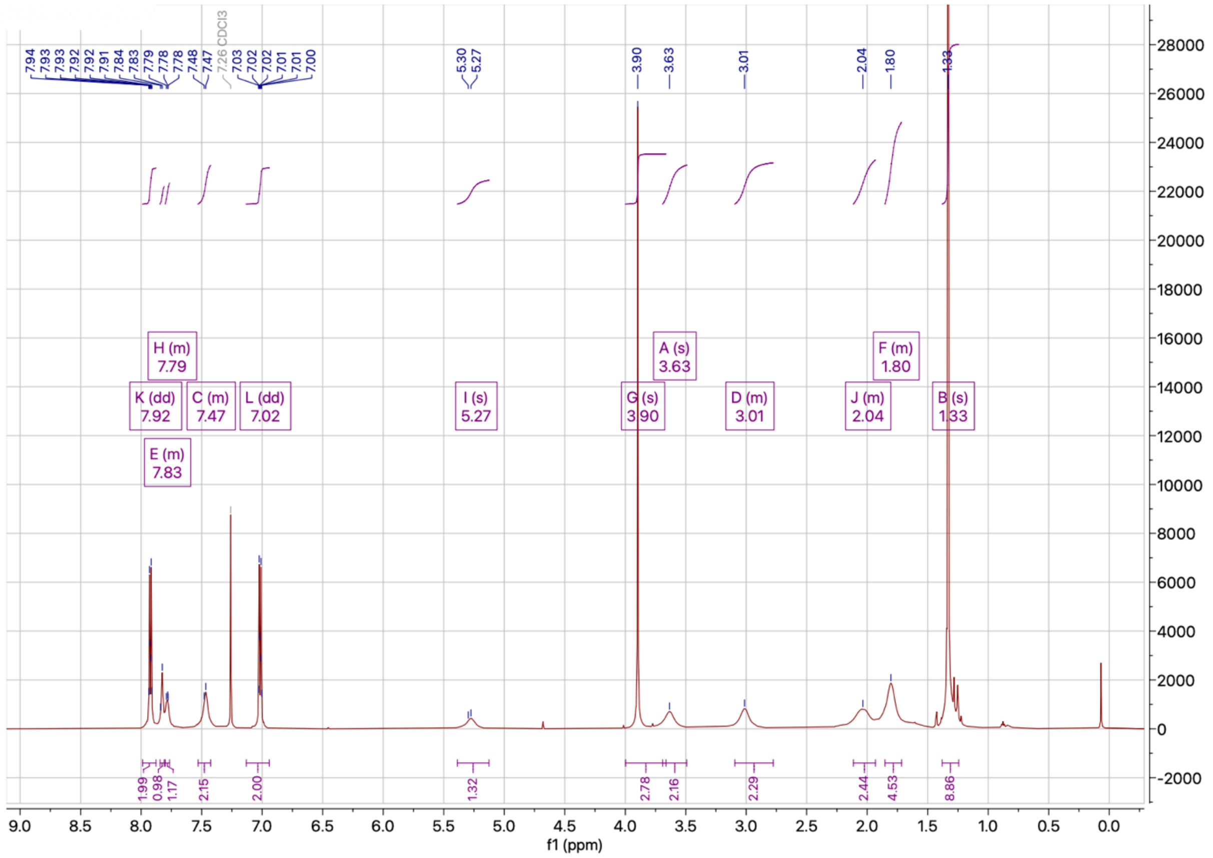


[13]C NMR spectrum of **3g**

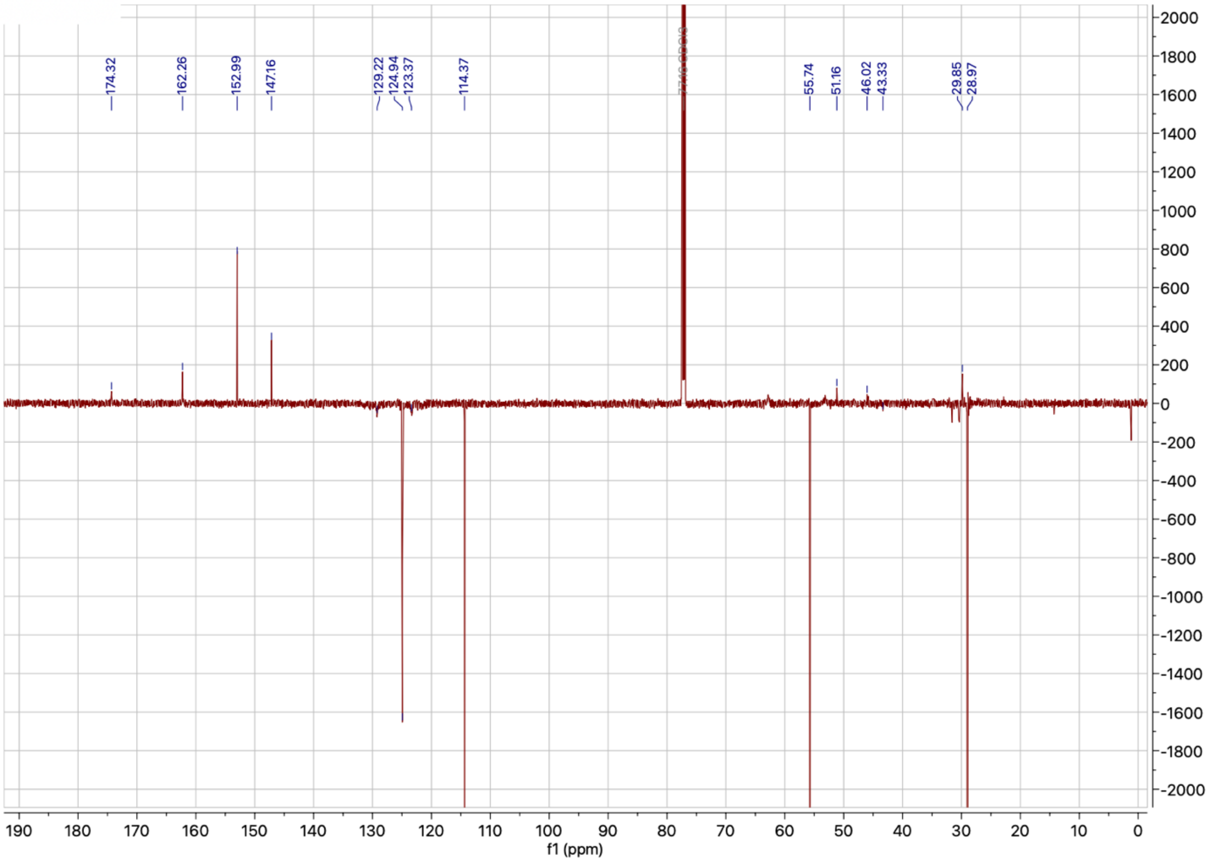

$^{1}$H NMR spectrum of **3h**

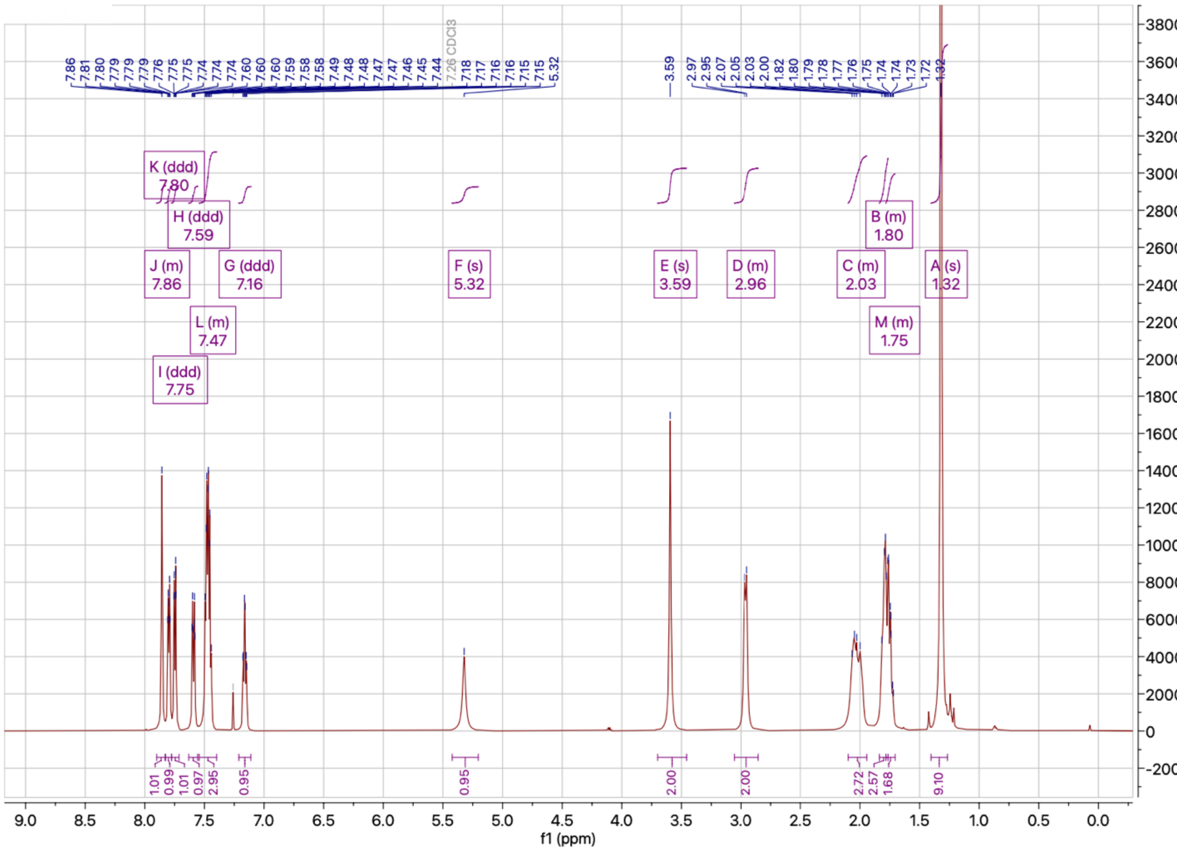

$^{13}$C NMR spectrum of **3h**

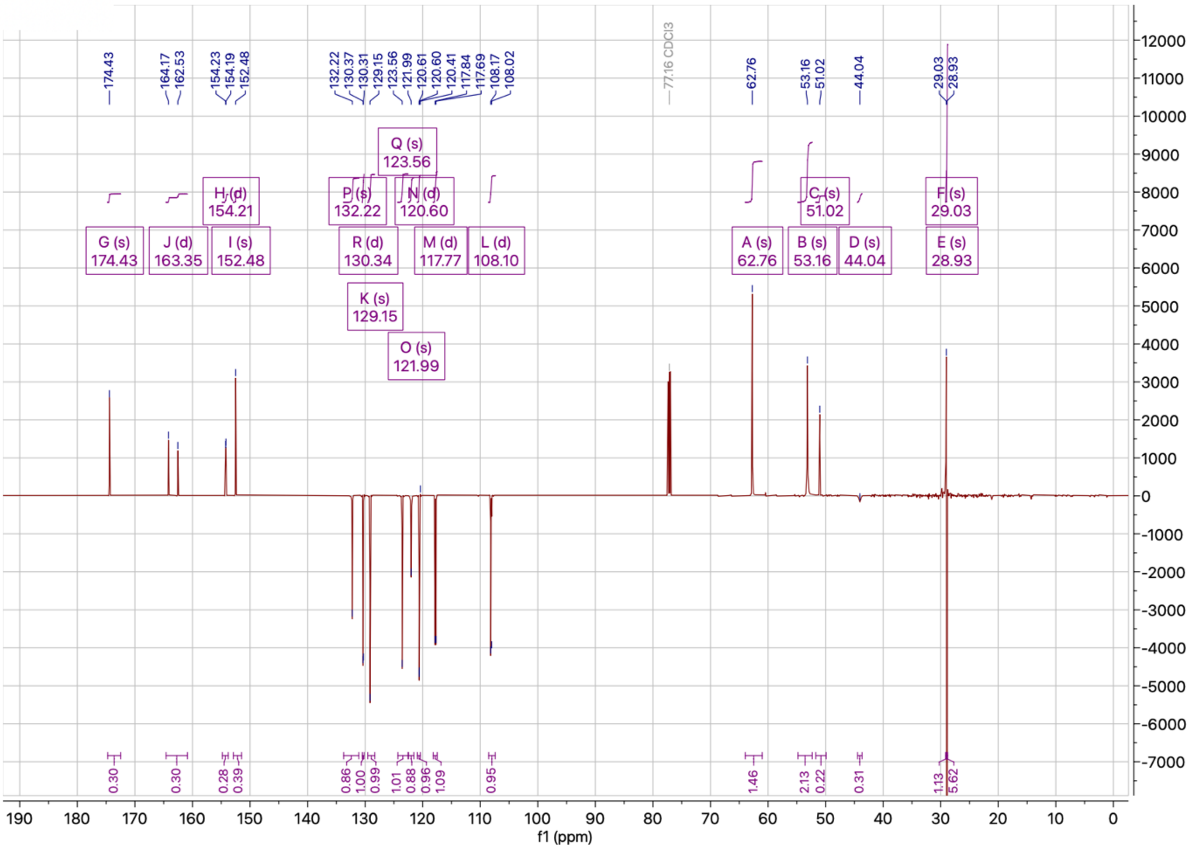

[1]H NMR spectrum of **3i**

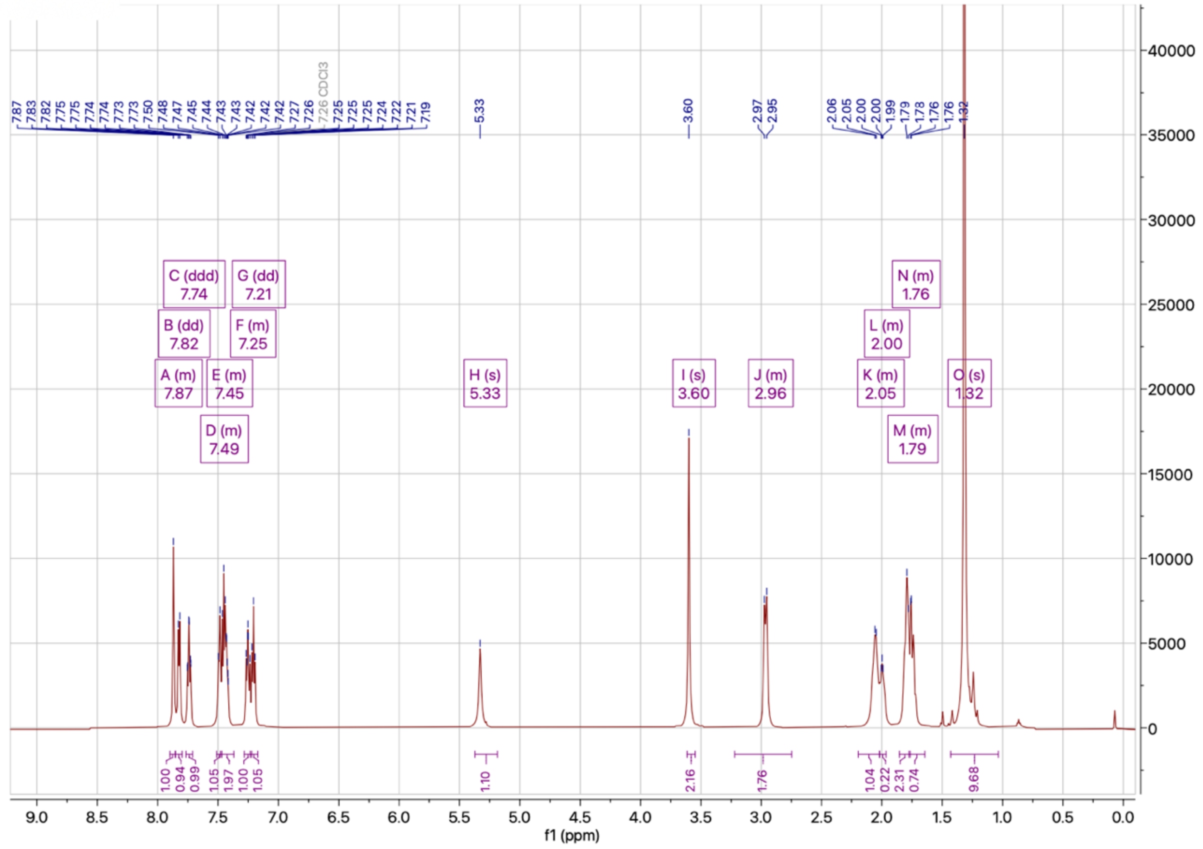


[13]C NMR spectrum of **3i**

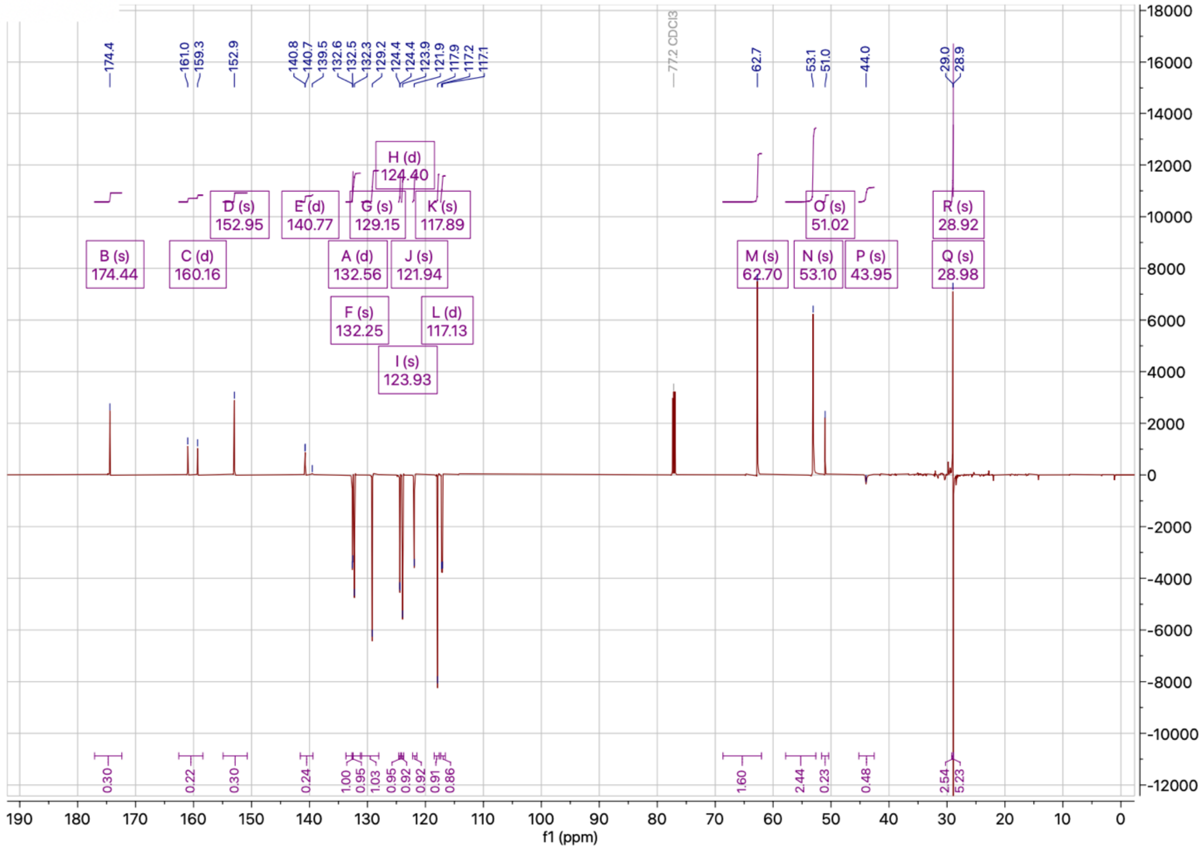

[1]H NMR spectrum of **3j**

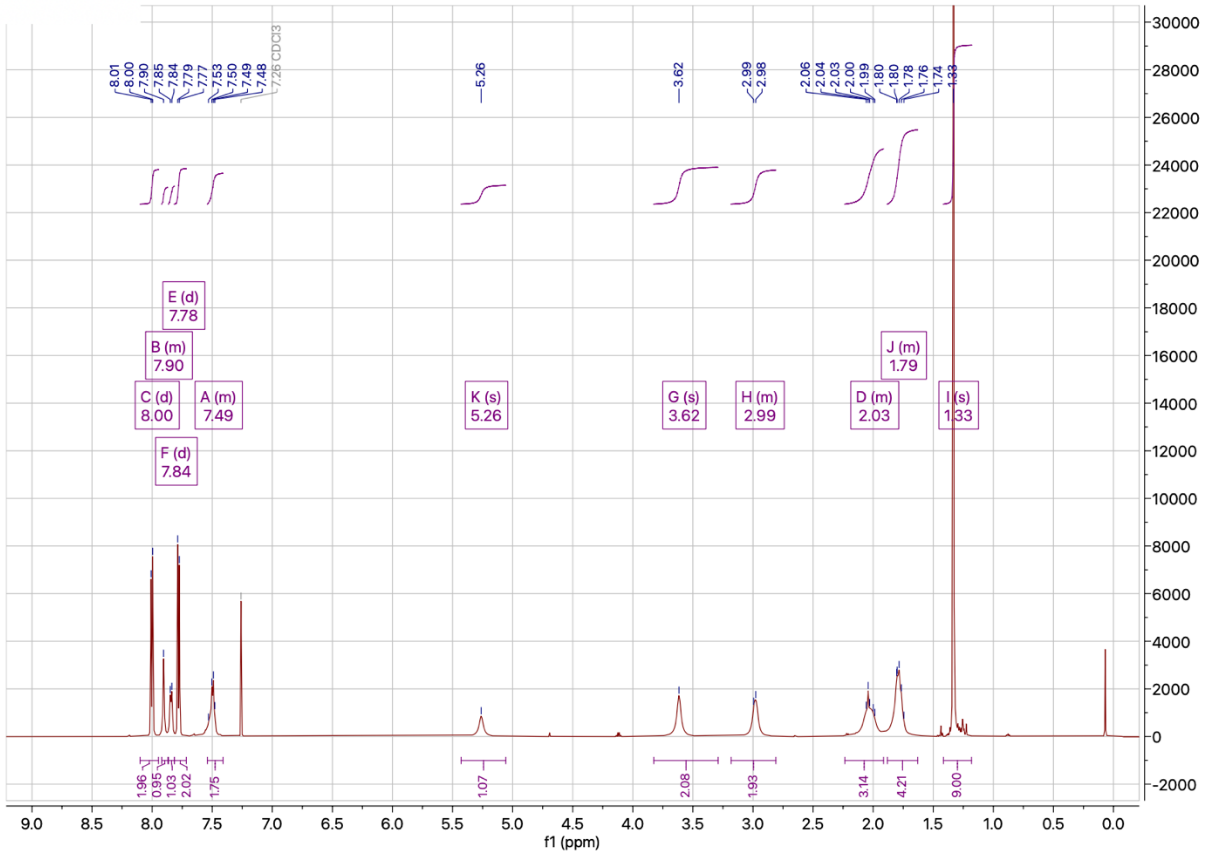

[13]C NMR spectrum of **3j**

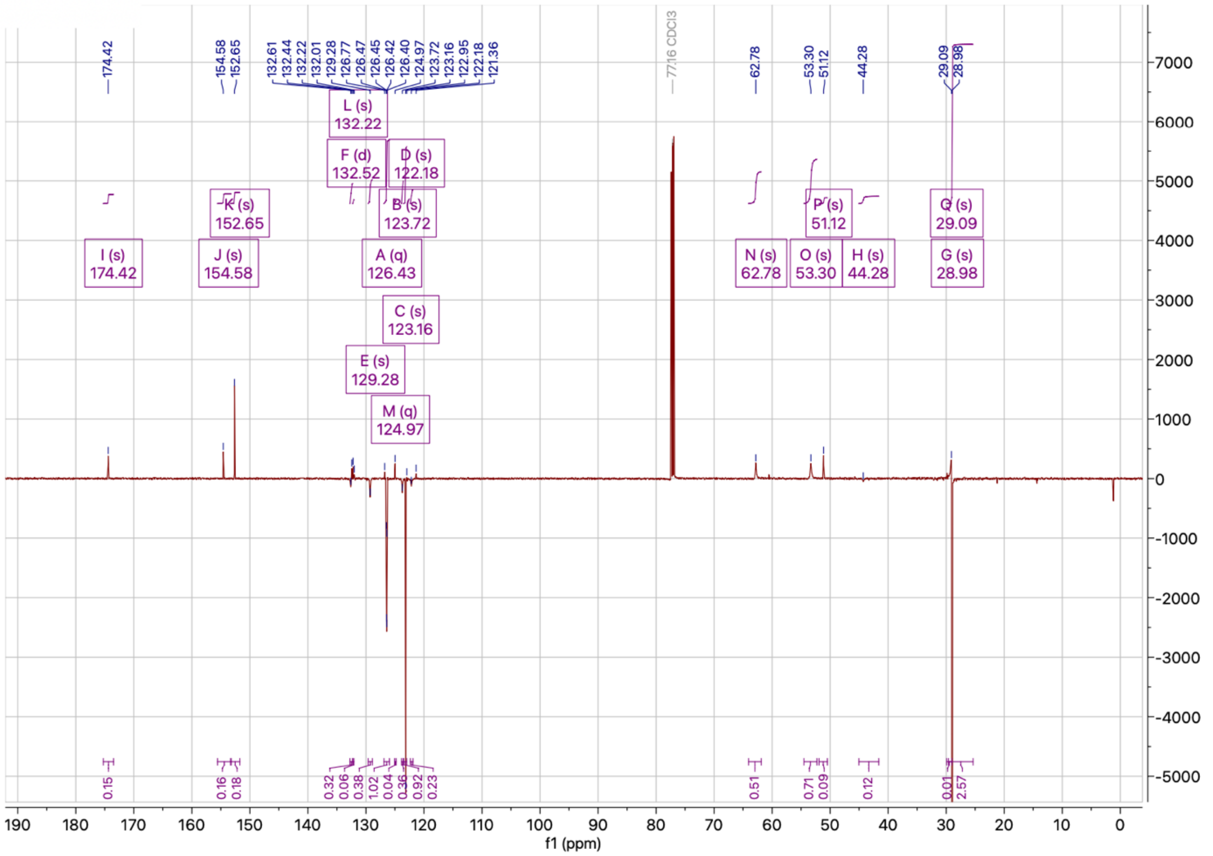

<sup></sup>

$^{1}$H NMR spectrum of **3k**

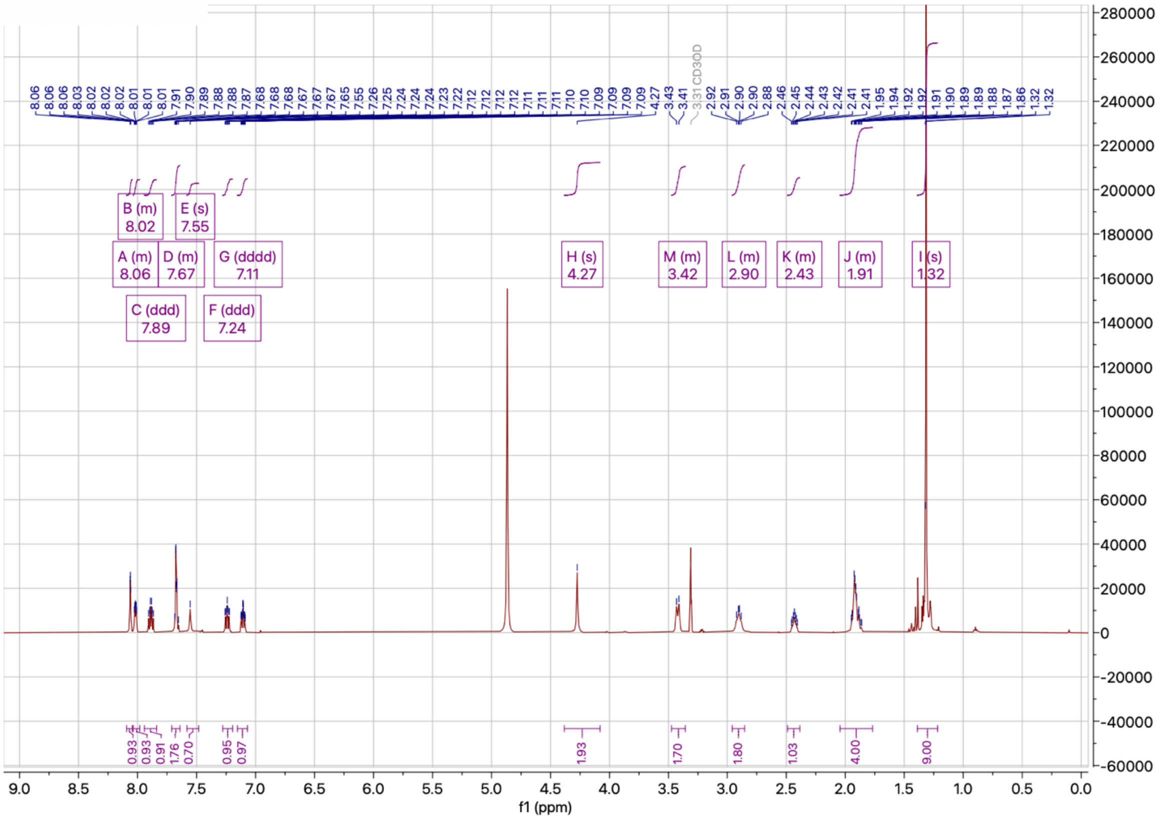

$^{13}$C NMR spectrum of **3k**

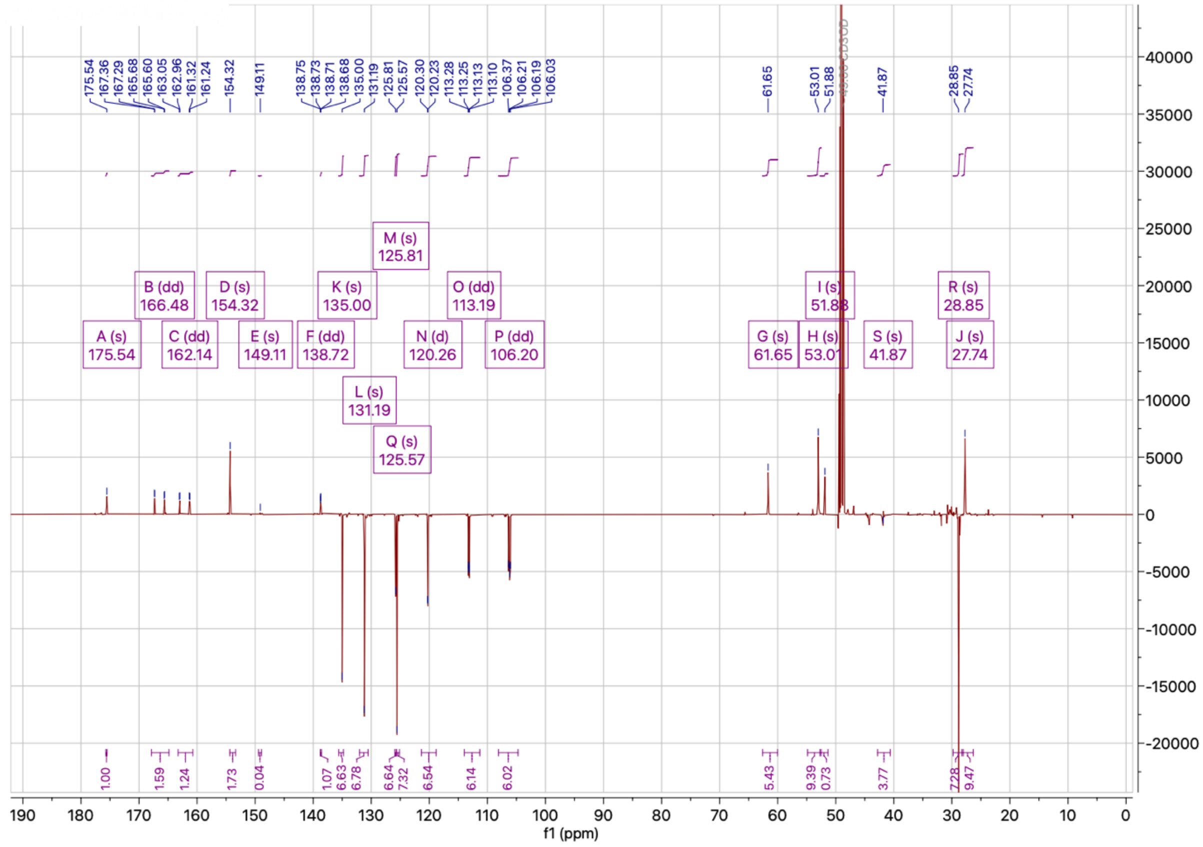

$^{1}$H NMR spectrum of **3l**

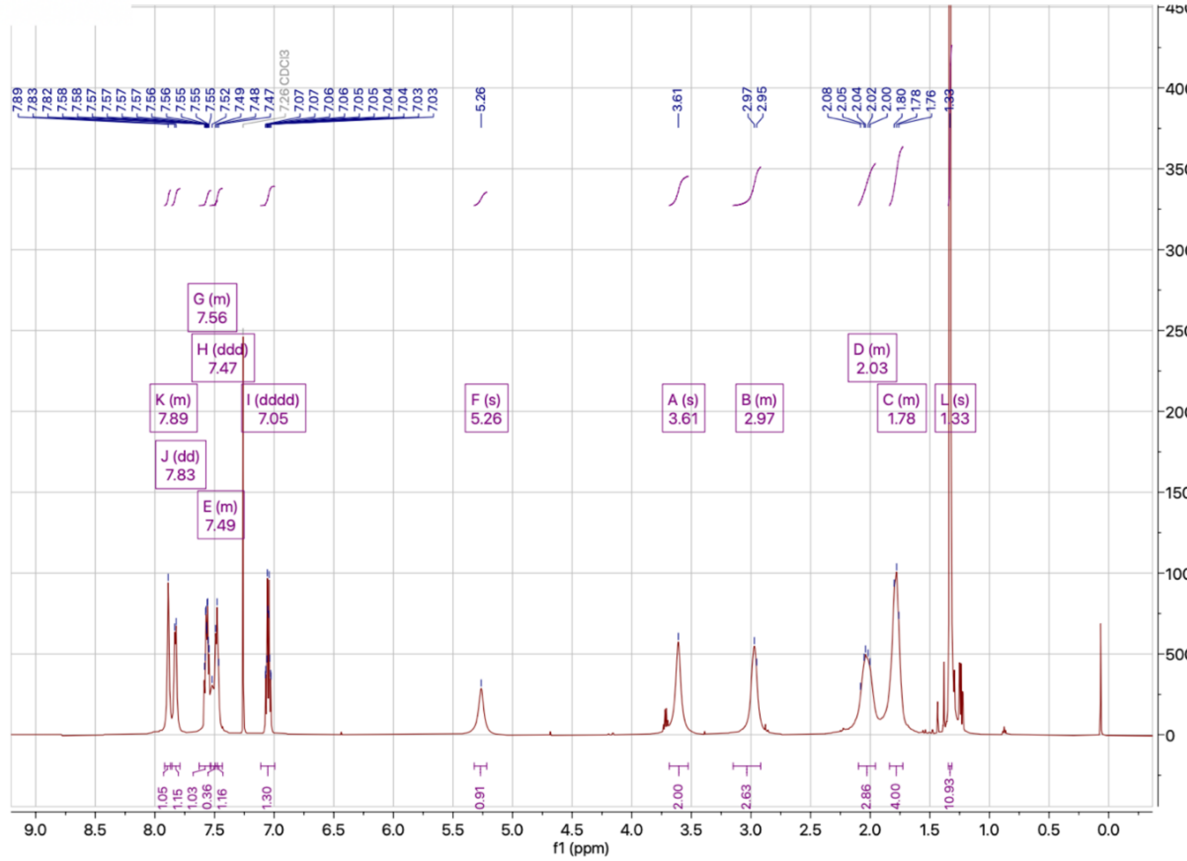

$^{13}$C NMR spectrum of **3l**

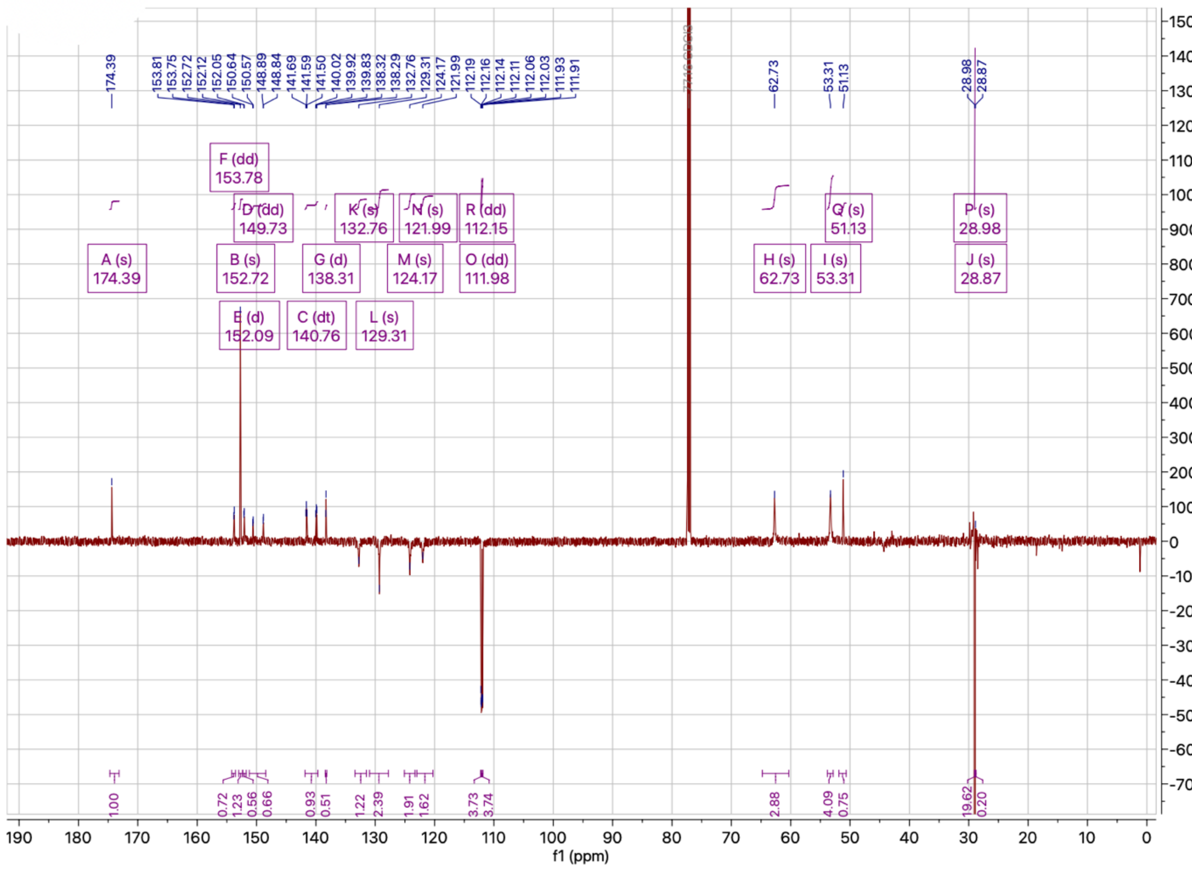

(8) MoKa. Molecular Discovery: Hertfordshire, UK.

(9) CORINA Classic—High-Quality 3D Molecular Models. MN-AM.

(10) Friesner, R. A.; Banks, J. L.; Murphy, R. B.; Halgren, T. A.; Klicic, J. J.; Mainz, D. T.; Repasky, M. P.; Knoll, E. H.; Shelley, M.; Perry, J. K.; Shaw, D. E.; Francis, P.; Shenkin, P. S. Glide: A New Approach for Rapid, Accurate Docking and Scoring. 1. Method and Assessment of Docking Accuracy. *J. Med. Chem.* **2004**, *47* (7). https://doi.org/10.1021/jm0306430.

(11) Halgren, T. A.; Murphy, R. B.; Friesner, R. A.; Beard, H. S.; Frye, L. L.; Pollard, W. T.; Banks, J. L. Glide: A New Approach for Rapid, Accurate Docking and Scoring. 2. Enrichment Factors in Database Screening. *J. Med. Chem.* **2004**, *47* (7). https://doi.org/10.1021/jm030644s.

(12) MacroModel. Schrödinger, LLC: New York, NY 2021.

(13) Groom CR, Bruno IJ, Lightfoot MP, W. S. The Cambridge Structural Database. *Acta Crystallogr B Struct Sci Cryst Eng Mater.* **2016**, *72* ((Pt 2)), 171–179. https://doi.org/10.1107/S2052520616003954.

(14) Bruno, I. J.; Cole, J. C.; Edgington, P. R.; Kessler, M.; Macrae, C. F.; McCabe, P.; Pearson, J.; Taylor, R. New Software for Searching the Cambridge Structural Database and Visualizing Crystal Structures. *Acta Crystallogr. Sect. B Struct. Sci.* **2002**, *58* (3 PART 1). https://doi.org/10.1107/S0108768102003324.

(15) Bruno, I. J.; Cole, J. C.; Kessler, M.; Luo, J.; Momerwell, W. D. S.; Purkis, L. H.; Smith, B. R.; Taylor, R.; Cooper, R. I.; Harris, S. E.; Orpen, A. G. Retrieval of Crystallographically-Derived Molecular Geometry Information. *J. Chem. Inf. Comput. Sci.* **2004**, *44* (6). https://doi.org/10.1021/ci049780b.

(16) Cottrell, S. J.; Olsson, T. S. G.; Taylor, R.; Cole, J. C.; Liebeschuetz, J. W. Validating and Understanding Ring Conformations Using Small Molecule Crystallographic Data. *J. Chem. Inf. Model.* **2012**, *52* (4). https://doi.org/10.1021/ci200439d.

(17) Bowers, K. J.; Chow, E.; Xu, H.; Dror, R. O.; Eastwood, M. P.; Gregersen, B. A.; Klepeis, J.

L.; Kolossvary, I.; Moraes, M. A.; Sacerdoti, F. D.; Salmon, J. K.; Shan, Y.; Shaw, D. E. Scalable Algorithms for Molecular Dynamics Simulations on Commodity Clusters. In *Proceedings of the 2006 ACM/IEEE Conference on Supercomputing, SC'06*; 2006. https://doi.org/10.1145/1188455.1188544.
